# In host evolution of *Exophiala dermatitidis* in cystic fibrosis lung micro-environment

**DOI:** 10.1101/2022.09.23.509114

**Authors:** Tania Kurbessoian, Daniel Murante, Alex Crocker, Deborah A. Hogan, Jason E. Stajich

## Abstract

Individuals with cystic fibrosis (CF) are susceptible to chronic lung infections that lead to inflammation and irreversible lung damage. While most respiratory infections that occur in CF are caused by bacteria, some are dominated by fungi such as the slow-growing black yeast *Exophiala dermatitidis*. Here, we analyze isolates of *E. dermatitidis* cultured from two samples, collected from a single subject two years apart. One isolate genome was sequenced using long-read Nanopore technology as an in-population reference to use in comparative single nucleotide polymorphism (SNP) and insertion-deletion (INDEL) variant analyses of twenty-three isolates. We then used population genomics and phylo-genomics to compare the isolates to each other as well as the type strain *E. dermatitidis NIH/UT8656.* Within the CF lung population, three *E. dermatitidis* clades were detected, each with varying mutation rates. Overall, the isolates were highly similar suggesting that they were recently diverged. All isolates were MAT 1-1, which was consistent with their high relatedness and the absence of evidence for mating or recombination between isolates. Phylogenetic analysis grouped sets of isolates into clades that contained isolates from both early and late time points indicating there are multiple persistent lineages. Functional assessment of variants unique to each clade identified alleles in genes that encode transporters, cytochrome P450 oxidoreductases, iron acquisition and DNA repair processes. Consistent with the genomic heterogeneity, isolates showed some stable phenotype heterogeneity in melanin production, subtle differences in antifungal minimum inhibitory concentrations, growth on different substrates. The persistent population heterogeneity identified in lung-derived isolates is an important factor to consider in the study of chronic fungal infections, and the analysis of changes in fungal pathogens over time may provide important insights into the physiology of black yeasts and other slow-growing fungi *in vivo*.

## Introduction

Cystic fibrosis (CF) is an autosomal recessive disorder caused by mutations in the cystic fibrosis transmembrane regulator (CFTR) gene that impair the balance of salts and water across epithelia. In the lungs, these ion transport defects cause viscous mucus which contributes to respiratory infections that cause most of the morbidity and mortality in CF populations (Riordan *et al*. 1989; Davis 2006; Ferec and Cutting 2012). Microbial colonization of mucosal plugs results in recurring infection and inflammation that cause irreversible lung damage and declining lung function (Turcios 2020). Bacteria, particularly *Staphylococcus aureus*, *Pseudomonas aeruginosa*, *Burkholderia cepacia* and *Stenotrophomonas maltophilia* are pathogens that frequently dominate CF respiratory infections (Burns *et al*. 1998; Mariani-Kurkdjian and Bingen 2003; Horré *et al*. 2004; Steinkamp *et al*. 2005; Tunney *et al*. 2008; Pihet *et al*. 2009; Zhao *et al*. 2012) and are sometimes isolated with various species of fungi. Clinically significant fungi in CF lung infections include *Exophiala dermatitidis*, *Scedosporium apiospermum*, *Aspergillus fumigatus*, *Candida albicans* and *Clavispora (Candida) lusitaniae* (Kusenbach *et al*. 1992; Cimon *et al*. 2000; Defontaine *et al*. 2002; Horré *et al*. 2004, 2009; Parize *et al*. 2014; Chen *et al*. 2018; Demers *et al*. 2018; Jong *et al*. 2020). The consequences of fungal infection on CF outcomes is not well understood but is influenced by the genotypes of both the host and microbes (Burns *et al*. 1998; Nagano *et al*. 2007; Pihet *et al*. 2009; Packeu *et al*. 2012).

*Exophiala dermatitidis*, previously named *Hormiscium dermatitidis* and *Wangiella dermatitidis*, are taxonomically classified in Phylum Ascomycota, Order Chaetothyriales, Family Herpotrichiellaceae. To date, about 40 species in the *Exophiala* genus have been identified, seventeen of which are known to cause disease in mammals. Among these, *E. dermatitidis* is the most clinically prevalent with reported mortality rates of 25–80% in systemic and invasive cases, even though fatal systemic cases are relatively rare (Kirchhoff *et al*. 2019). Clinical presentations of this fungus include phaeohyphomycosis, keratitis, chromoblastomycosis and even several neural diseases and meningitis (Revankar *et al*. 2002; Matos *et al*. 2002; Uijthof *et al*. 2009; Revankar and Sutton 2010; Seyedmousavi *et al*. 2014; Song *et al*. 2017; Kirchhoff *et al*. 2019; Lavrin *et al*. 2020). The first instance of *E. dermatitidis* to be isolated from a sputum culture procured from a cystic fibrosis patient was in 1990 (Haase *et al*. 1990; Kusenbach *et al*. 1992). Many studies have since isolated *E. dermatitidis* from CF sputum cultures (Rath *et al*. 1997; Diemert *et al*. 2001; Horré *et al*. 2004; Griffard *et al*. 2010; Packeu *et al*. 2012), some of which have led to the development of later state mycosis disease (Sudfeld *et al*. 2010; Kondori *et al*. 2011; Song *et al*. 2017; Grenouillet *et al*. 2018), treatment (Packeu *et al*. 2012; Mukai *et al*. 2014) or even death (Klasinc *et al*. 2019).

Exophiala species are black yeasts which are classified by three defining features. They produce melanin through 1-8 dihydroxynapthalene (DHN) biosynthesis pathway, exhibit morphological plasticity or meristematic growth (yeast cells, hyphae or even pseudohyphae), and have membrane associated carotenoids and an intracellular mycosporine-like amino acids (Hoog *et al*. 2000; Nosanchuk and Casadevall 2003, 2006; Saunte *et al*. 2012; Smith and Casadevall 2019). All of these properties likely contribute to the extreme resistance to environmental stresses including desiccation, UV or solar exposure, and ionizing radiation. These resistance traits may also contribute to success in growth in mammalian hosts and their ability to cause disease in susceptible hosts. The *E. dermatitidis* strain NIH/UT8656 genome was sequenced and assembled into 11 complete and contiguous chromosomes (Robertson *et al*. 2012; Chen *et al*. 2014; Schultzhaus *et al*. 2020; Malo *et al*. 2021) which has enabled comparative genomics and identification of the genes which may underlie its resilience (Robertson *et al*. 2012; Schultzhaus *et al*. 2020; Malo *et al*. 2021) and success in human host colonization (Kondori *et al*. 2011; Kirchhoff *et al*. 2019).

Reservoirs of *E. dermatitidis* have long been associated with hot and humid tropical origins (Sudhadham *et al*. 2008). *E. dermatitidis* are isolated from many man-made substrates found in humidifiers (Nishimura and Miyaji 1982), saunas (Matos *et al*. 2002), and dishwashers (Zupančič *et al*. 2016; Babič *et al*. 2018). Babic et al. (Babič *et al*. 2018) concluded that *Exophiala* tends to be found in locations with oligotrophic conditions or where rubber seals and humidity act as an enrichment or trapping mechanism which supports *Exophiala* persistence. *E. dermatitidis* and related species are also found in broad environmental niches including wasp nest (Conti-Díaz *et al*. 1977), healthy bats (Reiss and Mok 1979), lesions of toads (Frank *et al*. 1970), rotten wood (Dixon *et al*. 1980) which suggests human patients acquire infections from environmental exposure (Sudhadham *et al*. 2011).

Successful microbial invasions require iron, a critical growth-limiting factor, which must be sequestered from the surrounding environment through the release of iron chelating proteins called siderophores (Neilands 1995; Mossialos and Amoutzias 2009). For example, *Pseudomonas aeruginosa* produces the fluorescent siderophores, pyoverdine and pyochelin, which are used to sequester iron from the lung environment and as cofactors for respiratory proteins needed for surface motility and biofilm maturation (Haase *et al*. 1990; Banin *et al*. 2005; Matilla *et al*. 2007). Another example, the fungus *Aspergillus fumigatus* produces two hydroxamate-type siderophores: triacetylfusarinine (SidD) and ferricrocin (SidC), while also producing fusarinine and hydroxyferricrocin type siderophores (Schrettl *et al*. 2007; Haas 2014). In a study by Zajc et al., (Zajc *et al*. 2019), indicated 10 predicted siderophores in four *Exophiala dermatitidis* genomes, two of which include triacetylfusarinine and ferricrocin. Ferricrocin is an internal siderophore used to store iron and is essential for sexual development and contributes to oxidative stress resistance (Schrettl *et al*. 2007; Tyrrell and Callaghan 2016). Triacetylfusarinine is used to facilitate hyphal growth under iron-depleted conditions (Schrettl *et al*. 2007; Tyrrell and Callaghan 2016).

The *E. dermatitis* locus (HMPREF1120_07636) depicts the non-ribosomal peptide synthase SidC necessary for siderophore synthesis (Zajc *et al*. 2019; Malo *et al*. 2021). Polymicrobial infections persist in the CF lung environment through production and scavenging of extracellular siderophores which aid microbes in competition for resources (Tyrrell and Callaghan 2016; Sass *et al*. 2019; Yan *et al*. 2022). Microbes can use, obtain, and sequester iron in siderophores from hemoglobin found in red blood cells and lactoferrin contained in mucosal secretions. The competition and use of iron is an important dynamic in polymicrobial infections including fungal-bacteria competition within plant and animal hosts, and may sometimes assist in promoting the growth of their host (Crowley *et al*. 1991; Johnson 2008; Aznar *et al*. 2014; Kim 2018; Mochochoko *et al*. 2021; Pohl Carolina H. and Noverr Mairi C.).

In light of several studies that chronic CF-related fungal infections diversify over time (Kim *et al*. 2015; Demers *et al*. 2018; Jones *et al*. 2020; Ross *et al*. 2021), here we report both phenotypic and genomic diversity among *E. dermatitidis* isolates from a single individual. Population genomic analyses identified multiple lineages that persisted over two years. This data set allows us to test the hypothesis that, as seen in previous studies, the CF lung environment supports stably diverged populations of a clonally derived yeast.

## Methods

### Sputum-derived isolate cultures

Frozen sputum was obtained from a specimen bank in which samples were obtained in accordance with protocols approved by the Dartmouth-Hitchcock Institutional Review Board. Aliquots of sputum were plated onto Sabouraud Dextrose Agar (SAB) medium as described previously (Grahl *et al*. 2018). Individual isolates were obtained and banked; isolate identifiers are listed in **Supplemental Table 1.**

### DNA extraction and sequencing

*Exophiala* isolates were grown in Yeast Peptone Dextrose media (YPD) for approximately 24 hours in 5 ml roller-drum cultures at 37 °C. Cells were spun down for 5 minutes at 5000 RCF and washed thrice with deionized water. Genomic DNA was extracted from cell pellets using the MasterPure yeast DNA purification kit (Epicentre). Melanin was removed from genomic DNA using the *OneStep*^TM^ PCR Inhibitor Removal kit (Zymo Research). Genomic DNA was measured by Nanodrop and diluted to ∼20ng/µl. DNA extractions were sent to Novogene, (Novogene Corporation Inc., Cambridge, United Kingdom) for 2×150bp sequencing on an Illumina NovoSeq 6000. DNA from isolate DCF04 was also extracted and sequenced on Oxford Nanopore (ONT) platform with library preparation and sequencing following manufacturer’s directions (Oxford Nanopore, Oxford United Kingdom). Flow cell versions FAK67997 and FAK73296 were used along with base-calling using guppy (*v. 3.4.4+a296cb*) (Wick *et al*. 2019).

### Genome assembly and annotation

Genome assemblies were constructed for the twenty-three *E. dermatitidis* isolates from Illumina sequencing. One isolate, Ex4, was also re-sequenced using Oxford Nanopore technology. All genomes were *de novo* assembled with AAFTF pipeline (*v.0.2.3*) (Palmer and Stajich 2022) which performs read QC and filtering with BBTools bbduk (*v.38.86)* (Bushnell 2014) followed by SPAdes (*v.3.15.2*) (Bankevich *et al*. 2012) assembly using default parameters, followed by screening to remove short contigs < 200 bp and contamination using NCBI’s VecScreen. The BUSCO ascomycota_odb9 database (Manni *et al*. 2021) was used to determine how complete the assembly was for all 23 isolates of *E. dermatitidis.* A hybrid assembly of isolate DCF04/Ex4 was generated using MaSUrCa (*v.3.3.4*) (Zimin *et al*. 2013) as the assembler using both Nanopore and Illumina sequencing reads. General default parameters were used except: CA_PARAMETERS=cgwErrorRate=0.15, NUM_THREADS=16, and JF_SIZE=200000000. The updated genome was then scaffolded to strain NIH/UT8656 accession GCF_000230625 using Ragtag (*v.1.0.2*) (Alonge *et al*. 2019) which uses minimap2 (*v. 2.17-r941)* (Li 2018) to further link scaffolds based on shared co-linearity of these isolates’ genomes.

We predicted genes in this near complete genome assembly with Funannotate (*v1.8.1*) (Palmer and Stajich 2020). A masked genome was created by first generating a library of sequence repeats with the RepeatModeler pipeline (Smit and Hubley 2008). These species-specific predicted repeats were combined with fungal repeats in the RepBase (Bao *et al*. 2015) to identify and mask repetitive regions in the genome assembly with RepeatMasker (*v.4-1-1*) (SMIT A. F. A 2004). To predict genes, *ab initio* gene predictors SNAP (*v.2013_11_29*) (Korf 2004) and AUGUSTUS (*v.3.3.3*) (Stanke *et al*. 2006) were trained using the Funannotate ‘train’ command based on the full-length transcripts constructed by Genome-Guided run of Trinity (*v.2.11.0*) (Grabherr *et al*. 2011) using RNA-Seq from published E*. dermatitis* SRA accession SRS282040. The assembled transcripts were aligned with PASA (*v.2.4.1*) (Haas *et al*. 2008) to produce full-length spliced alignments and predicted open reading frames for training the *ab initio* predictors and as informant data for gene predictions. Additional gene models were predicted by GeneMark.HMM-ES (*v.4.62_lic*) (Brůna *et al*. 2020), and GlimmerHMM (*v.3.0.4*) (Majoros *et al*. 2004) that utilize a self-training procedure to optimize *ab initio* predictions. Additional exon evidence to provide hints to gene predictors was generated by DIAMOND BLASTX alignment of SwissprotDB proteins and polished by Exonerate (*v.2.4.0*) (Slater and Birney 2005). Finally, EvidenceModeler (*v.1.1.1*) (Haas *et al*. 2008) generated consensus gene models in Funannotate were constructed using default evidence weights. Non-protein-coding tRNA genes were predicted by tRNAscan-SE (*v.2.0.9*) (Lowe and Chan 2016).

The annotated genome was processed with antiSMASH (*v.5.1.1*) (Blin *et al*. 2021) to predict secondary metabolite biosynthesis gene clusters. These annotations were also incorporated into the functional annotation by Funannotate. Putative protein functions were assigned to genes based on sequence similarity to InterProScan5 (*v.5.51-85.0*) (Jones *et al*. 2014), Pfam (*v.35.0*) (Finn *et al*. 2014), Eggnog (*v.2.1.6-d35afda*) (Huerta-Cepas *et al*. 2019), dbCAN2 (*v.9.0*) (Zhang *et al*. 2018) and MEROPS (*v.12.0*) (Rawlings *et al*. 2018) databases relying on NCBI BLAST (*v.2.9.0+)* (Sofi *et al*. 2022) and HMMer (*v.3.3.2*) (Potter *et al*. 2018). Gene Ontology terms were assigned to protein products based on the inferred homology based on these sequence similarity analyses. The final annotation produced by Funannotate was deposited in NCBI as a genome assembly with gene model annotation.

Copy number variation (CNV) was examined by plotting window-based read coverage of the short-read alignments of each isolate. The depth of coverage was calculated using mosdepth (Pedersen and Quinlan 2018), and visualized with R using the ggplot2 package (Wickham 2016).

The Mating Type (MAT) locus was identified through searching for homologous MAT genes (HMPREF08862) and (HMPREF05727) in this study’s 23 *E dermatitidis* isolate genomes with cblaster (Gilchrist *et al*. 2021). A homothallic black yeast, *Capronia coronata* CBS 617.96 (AMWN00000000.1) (Teixeira *et al*. 2017), which has both MAT genes, was also incorporated into the analyses and visualization. The identified homologous regions were examined for their conserved synteny of the MAT locus using clinker (Gilchrist and Chooi 2021) and a custom Biopython script (Cock *et al*. 2009) (Kurbessoian 2022) to extract the annotated region of the genome which contained the locus.

Identification of telomeric repeat sequences was performed using FindTelomeres.py script (https://github.com/JanaSperschneider/FindTelomeres). Briefly, this searches for chromosomal assembly with a regular expression pattern for telomeric sequences at the 5’ and 3’ end of each scaffold. Telomere repeat sequences were also predicted using A Telomere Identification toolkit (tidk) (*v.0.1.5*) “explore” option (https://github.com/tolkit/telomeric-identifier).

### Identification of sequence variation

Sequence variation among isolates was assessed using the best practices of the Genome Analysis ToolKit GATK (*v. 4.0.4.0*) (McKenna *et al*. 2010; Franke and Crowgey 2020) to identify SNPs and Insertion/Deletions (INDEL). Illumina paired-end reads were aligned to isolate DCF04 assembly with BWA (*v.0.7.17*) (Li and Durbin 2010) and processed with Samtools (*v.1.8*) (Li *et al*. 2009) and Picard Toolkit (Institute; Toolkit) AddOrReplaceReadGroups and MarkDuplicates (*v.2.18.3*). The alignments were further improved by realigning reads near inferred INDELs using GATK tools RealignerTargetCreator, IndelRealigner, and PrintReads. Genotypes were inferred with the GATK Haplotype and GenotypeGVCF methods to produce a single VCF file of the identified variants. Low quality SNPs were further filtered using GATK VariantFiltration and finally SelectVariants was used with the parameters: mapping quality (score < 40), quality by depth (<2 reads), Strand Odds Ratio (SQR > 4.0), Fisher Strand Bias (> 200), and Read Position Rank Sum Test (< −20) to retain only high-quality polymorphisms. Finally, an additional stringent series of three filtering steps implemented in bcftools (*v. 1.12*)(Li et al. 2019) was used on the VCF file to remove calls that were below the 1000 quality score threshold, where any individual isolate had a “no call”, and where the standard deviation in read depth (DP) was above or below a standard deviation value of 1 for an individual SNP. SnpEff (*v.4.3r*) (Cingolani *et al*. 2012) was used to score the impact of the identified variants using the Funannotate annotated DCF04 genome GFF3 file.

Variant calling was performed on two sets of individuals, one limited to the twenty-three CF patient population isolates with reads aligned to the DCF04 isolate and one using the *E. dermatitidis* NIH/UT8656 type strain. Pairwise isolate comparisons of SNP and INDEL were counted to generate isolate correlation heatmaps for both variant types using a UPGMA clustering. A custom script make_diagonal.sh uses plink (*v.2.00a3LM_AVX2*) (Chang *et al*. 2015) to count all pairwise differences between individuals in the VCF files stratified by SNPs or INDELs. A custom Perl script transformed pairwise counts into a matrix of isolated differences observed for both SNP and INDEL variants. Counts were summarized as heatmaps with a R script. To summarize the matrix plots, a distance plot using regression statistics was applied on both SNP counts and INDEL counts. A regression plot and statistics for the slope of the progression line, Pearson’s R, R-squared and the *p*-value were computed with a R script. All scripts developed for this manuscript are available at the Github (Kurbessoian 2022) project linked in this paper.

A calculation of the population mutation rate was performed on each isolate based on the number of SNPs shared among a pair of isolates. The formula to calculate the mutation rate per year for each isolate is as follows: (SNP Pairwise Value) / (Adjusted Genome Length) / Pair / Year. The time between isolated collections was 22 months. The value used for the “Adjusted Genome Length” was collected from running the assembler AAFTF pipeline (*v.0.2.3*) (Palmer and Stajich 2022). A one-way ANOVA was run on the grouped calculated mutation rates for each isolate to determine significance.

Genome comparison with dot-plot was constructed with D-GENIES (Cabanettes and Klopp 2018) using minimap2 and default parameters through the website for the tool. Longer isoform proteins were extracted for each strain genome annotation in order to call a more accurate gene count. Using Orthofinder (Emms and Kelly 2019) and DIAMOND ultra-sensitive parameters (Buchfink *et al*. 2015), assessment of overlap in the predicted protein-coding gene sequences from the genomes of *E. dermatitidis DCF04* and *E. dermatitidis NIH/UT8656* protein genomes was generated.

### Phylogenetics relationships of the isolates

SNPs from polymorphic sites were extracted from the VCF files as multi-fasta files using BCFTools (Li *et al*. 2019) and a custom script make_strain_tree.sh. A Maximum Likelihood phylogenetic tree was constructed from the multi-fasta file using IQTree (*v. 2.0.4*) (Minh *et al*. 2020) and the model parameters [-m GTR+ASC]. The chosen nucleotide substation model was GTR+F+ASC selected based on Bayesian information criteria (BIC). Statistical support for the tree nodes was evaluated from 1000 bootstrap replicates using UFBoot ultra-fast bootstrapping approximation (Hoang *et al*. 2018). The tree was visualized using iTOL (Letunic and Bork 2016).

### Phenotype assays

*Exophiala* isolates were streaked from −80°C onto yeast extract peptone dextrose (YPD) plates (2% glucose, 2% yeast extract, 1% peptone) and allowed to grow for 48 hours at 37 °C. Overnight cultures were started from YPD patches inoculated into 5 ml of liquid YPD and grown for approximately 24 hours in 5 ml rolling barrel cultures at 30 °C. For MIC assays, cultures were spun down for 5 minutes at 5000 RCF and washed thrice in deionized water. Cells were counted on a hemocytometer and added to a final concentration of 1000 colony forming units per well in a 96-well flat-bottom dish, then grown at 37°C for 72 hours before measuring final MIC.

## Results

A molecular and culture-based analysis of a series of sputum samples identified an individual with CF with a chronic lung infection caused by *E. dermatitidis* (Grahl *et al*. 2018)*. E. dermatitidis* isolates were recovered from banked sputum samples, collected two years apart. *Staphylococcus aureus* and *Candida albicans* were also identified in clinical cultures from the patient in the intervening years between the two timepoints (**Figure 1**). The subject’s antimicrobial use history included Aztreonam, Azithromycin, Tobramycin, Ciprofloxacin, and Doxycycline and the patient’s lung function, measured by percent predicted forced expiratory volume (%FEV1), ranged between 80 and 49% during this time period. While *E. dermatitidis* was not detected in the first clinical microbiological analysis, perhaps due to its extremely slow growth out of clinical samples (Grahl *et al*. 2018) or suppression by bacteria, it was detected in the second clinical analysis. Twenty-three isolates (eleven from the early time point and twelve from the late timepoint) were selected for further population genomic study.

**Figure 1.**
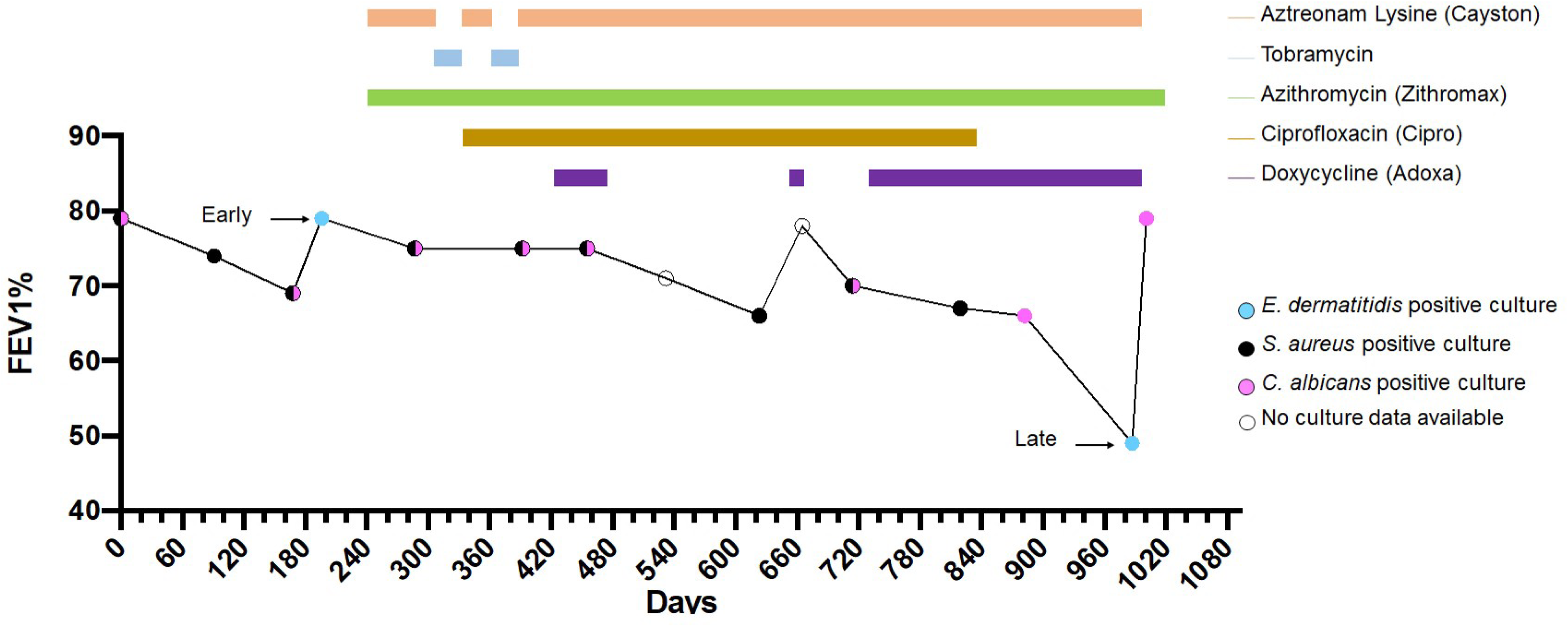
FEV1 pulmonary function data. FEV1 pulmonary function data was tested over the course of three years, and sputum cultures were acquired at each indicated time point and assessed for the presence of pathogens other than the mixed bacteria that compose normal upper respiratory flora. Colored bars indicate the duration of treatment for each listed antimicrobial given during infection.

### Sequencing and assembly of E. dermatitidis isolates

To gain information on the genetic variation and potential population structure for the recovered *E. dermatitidis* isolates, we sequenced and assembled the genomes of the twenty-three isolates (**Supplemental Table 2**). The depth of coverage ranged from 11-47x coverage across all 23 Illumina sequenced samples. The BUSCO (Benchmarking Universal Single Copy Orthologs) is another tool used to determine genome assembly completeness; within our dataset it ranged from 99-99.3 % complete, 98.3-99.3 single copies present, 0-1 ranged in duplicates, with a range of 0-0.3 in missing BUSCOs. Contig counts ranged from 44-1422, while the average genome assembly size is 26,706,323 Mbp, L50 about 1,078,277 and N50 of 9.

To further examine the fine-scale variation within the CF isolates we sought to generate a within population high-quality reference genome. The isolate DCF04 was sequenced using Oxford Nanopore (ONT) long reads to a genome depth of coverage at 11x and a hybrid genome assembly constructed from both ONT and Illumina reads (**Table 1; Supplemental Table 2).** The 26.6 Mb assembly contained only 3.19% of identified repetitive elements. The candidate telomeric repeat units “TTTAGGG/CCCTAA” were identified as repeat arrays at both ends of five scaffolds, but also found as single pairs in the remaining 4 scaffolds, as would be expected for 9 complete chromosomes (**Supplemental Table 3**). Sixty-four tRNA models were predicted from the genome. While the total number of genes predicted was 10030, 9599 of which are protein coding. 15 secondary metabolite clusters, 37 biosynthetic enzymes, and 49 small COGs were predicted with antiSMASH.

**Table 1.**
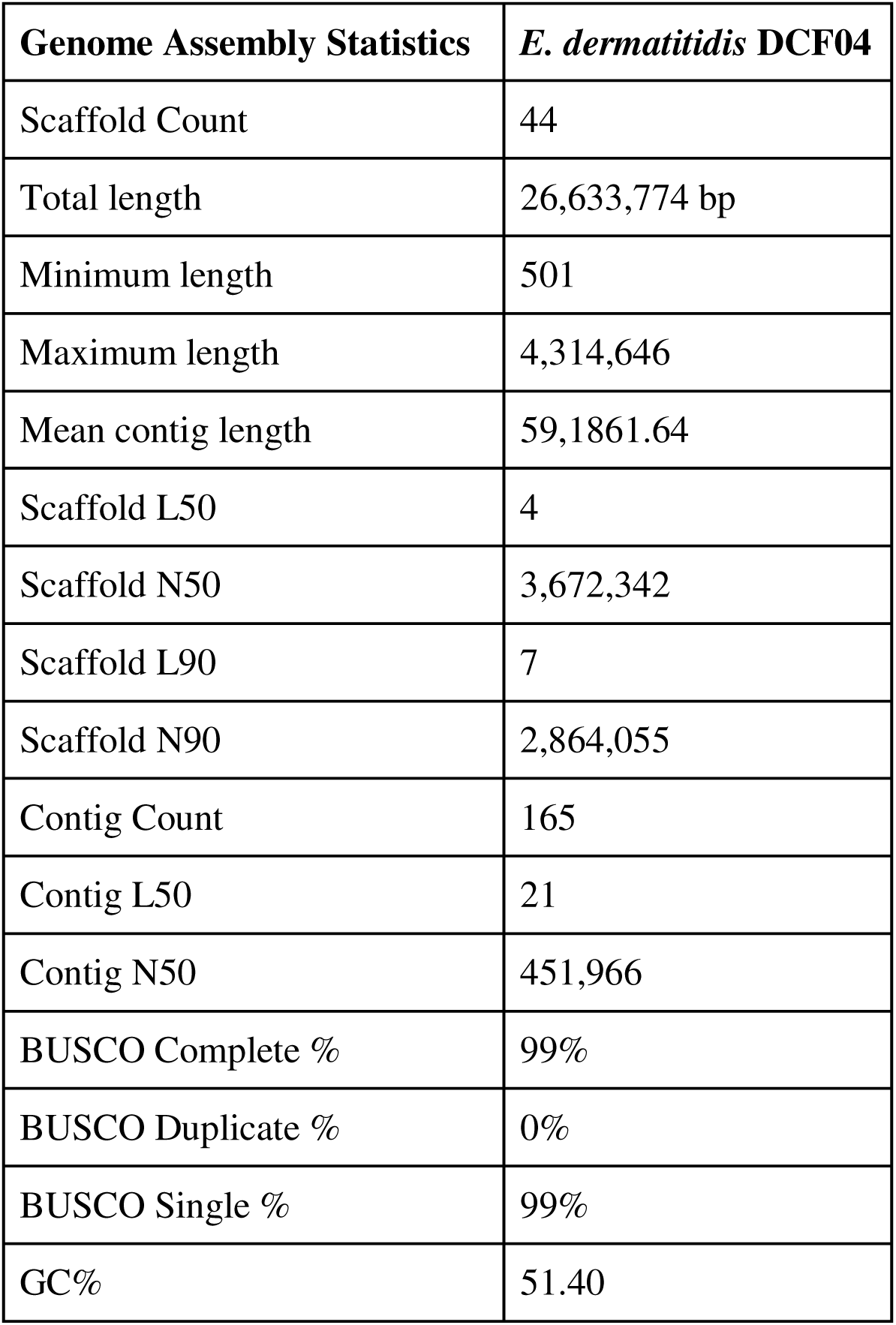
Genome assembly summary statistics for reference isolate *E. dermatitidis* DCF04. Table indicates assembly summary statistics of the hybrid assembly using Illumina and Nanopore sequencing of isolate DCF04. Summary statistics of contig and scaffold lengths are presented with genome completeness data calculated with BUSCO using the ascomycota_odb9 database.

**Table 2.** Genes with SNP variants found stratified by clade. Isolates collected in the early (red) and late (blue) time points are labeled in the comparisons. The functions of genes of interest found to have nonsynonymous mutations among entire clade of comparisons are reported with the total count of nonsynonymous and synonymous (NonSyn/Syn) variants found in each clade to identify the relative frequency of these changes and general functional differences in the types of genes with variants across the population clades.

| Clade | Isolates | Nonsynonymous mutations of interest in clade | Number of NonSynonymous/Synonymous SNPs unique to clade |
| --- | --- | --- | --- |
| I | Ex1, Ex2, Ex12, Ex14 | MADS box Transcription factor, G4 quadruplex binding protein, MFS transporter SP family sugar:H+ symporter, Bud site selection Bud4, DNA polymerase alpha subunit A, GTPase-activating protein | 59/7 |
| II | a) Ex3, Ex4, Ex8 | a. All hypothetical. | 56/8 |
|  | b) **Ex22, Ex17, Ex16, Ex7, Ex19, Ex23**<br>c) **Ex6, Ex10** | b. <u>Ex7 - Ex22</u> GO terms indicate Signal Transduction Mechanism and Transcription Factor, <u>Ex7-Ex17</u> Zinc finger binding protein TFIIIA, <u>Ex16</u> queuine tRNA-ribosyltransferase, MFS transporter SP family sugar:H<sup>+</sup> symporter, NAD-dependent histone deacetylase SIR2 and cytochrome P450, family 7, subfamily B (oxysterol 7-alpha-hydroxylase)<br>c. None. |  |
| III | a) **Ex9, Ex13, Ex11**<br>b) **Ex18, Ex15, Ex20, Ex21, Ex5** | a. Ex13 & Ex9, Ex11 Ap-3 complex subunit delta, DNA repair protein RAD50, and Regulator of nonsense transcripts 1-like protein.<br>b. Ex18 & Ex15, Ex20 an MFS transporter DHA2 family methylenomycin A resistance protein, sulfite reductase (ferredoxin), glycerol ethanol-ferric requiring protein, polyketide synthase, MFS transporter SP family solute carrier family 2 and a DNA repair protein RAD20 | 125/20 |

**Table 3.**
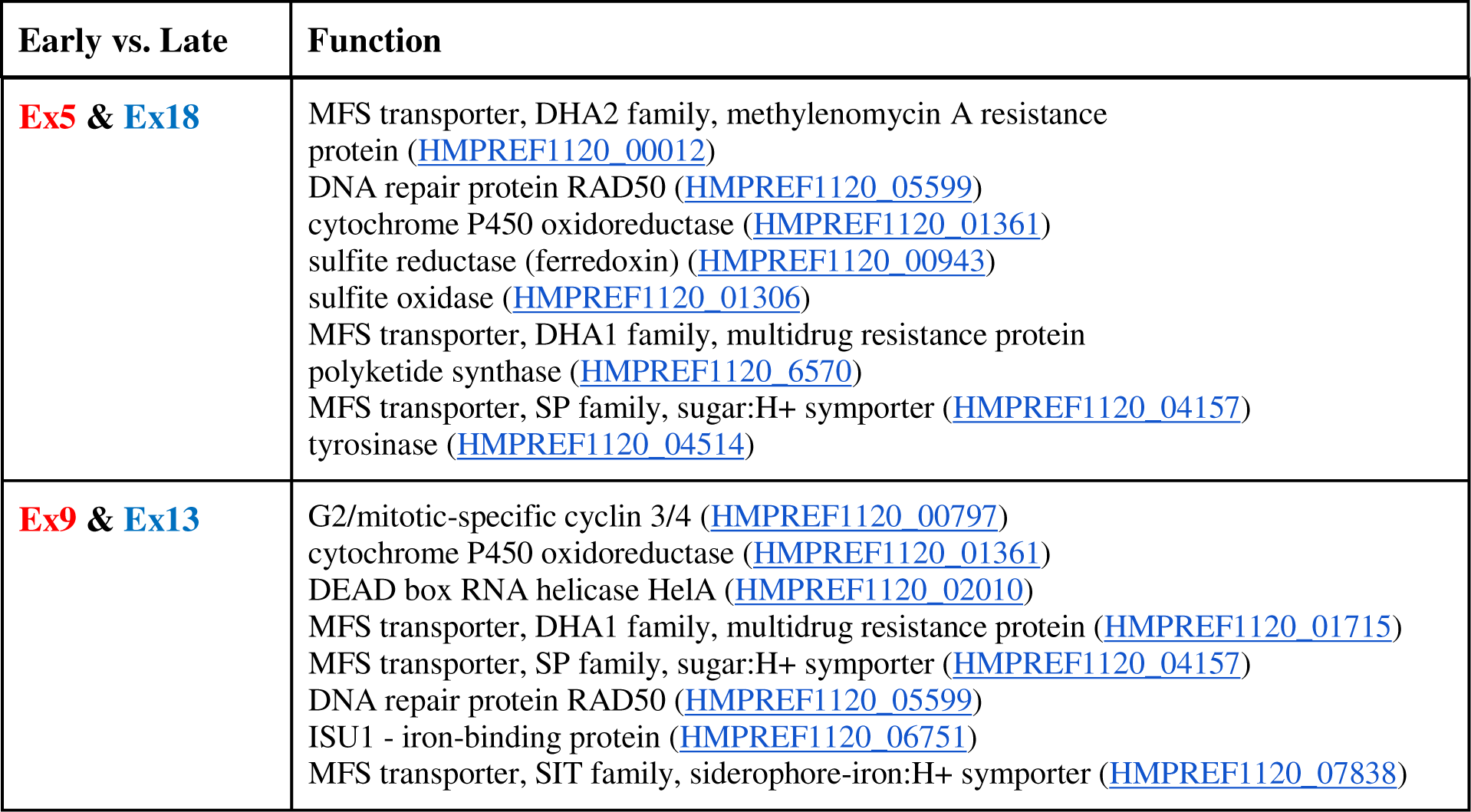
Genes with nonsynonymous SNP variants of interest observed in pairwise comparisons of *E. dermatitidis* CF lung isolates. Genes with notable non-synonymous differences identified in pairwise comparisons of early and late time point isolates. Full results of all pairwise differences are in **Supplemental** Table 7. For continuity, gene locus names are in parentheses and refer to the locus names in the reference strain *E. dermatitidis* NIH/UT8656.

### Comparing genome assembly and annotation of DCF04 to NIH/UT8656

Our DCF04 isolate is genetically close to the public strain *Exophiala dermatitidis NIH/UT8656* sequenced with Sanger sequencing technology (BioProject: PRJNA225511, Assembly: GCF_000230625.1). To assess the differences between the two genome assemblies we compiled summary statistics (**Table 1** and **Supplemental Table 2**) and interrogated the predicted gene content of both genomes. The DCF04 assembly had 165 contigs linked into 44 scaffolds, while NIH/UT8656 comprised 238 contigs linked into 10 scaffolds. Note that DCF04 scaffolds were derived by a comparative assembly against the NIH/UT8656 assembly to achieve best assembly after checking for rearrangements. The total length of the genome assembly is nearly the same for both DCF04 at 26.6Mb and NIH/UT8656 at 26.4Mb. The summary statistics for scaffold L50 and N50 are also nearly identical for DCF04 at 4 and 3.7Mb and NIH/UT8656 were 4 and 3.6Mb. A dot-plot comparing the two genome assemblies revealed minimal rearrangements or discontinuity suggesting high similarity of the two isolated genomes (**Supplemental Figure 1**).

The genome content was further compared using OrthoFinder (Emms and Kelly 2019) (**Supplemental Table 4).** OrthoFinder identified 8,256 orthologous groups or 17,640 orthologous protein-coding genes between the two genomes. Both strains had a number of unassigned genes that were given orthogroups, 705 for DCF04 and 475 for NIH/UT8656, along with 58 genes that were assigned, 34 DCF04 genes and 24 NIH/UT8656 genes. Of the 34 isolate-specific assigned genes in DCF04, 21 had identifiable fungal homologs with NIH/UT8656 including ABC multidrug transporters, AAT family amino acid transporter, 5-oxoprolinase and hypothetical protein/P-loop containing nucleoside triphosphate hydrolase protein. Two of the 13 assigned protein-coding genes resulted in a ribonuclease HI protein, while the remaining assigned 11 resulted in hypothetical protein matches. 4 out of the 5 orthogroups specific to NIH/UT8656 (22 protein-coding genes) were identical to the 4 seen in DCF04, while 1 orthogroup matched to a hypothetical protein/DUF300-domain containing protein. When analyzing the unassigned genes, these functionally had no known paralog on NCBI. We believe these results reflect differences in gene prediction pipelines as much as it could be due to gene content differences (Weisman *et al*. 2022).

### All Ex CF isolates are MAT1-1 mating type

The small-scale genome synteny evaluation tool clinker was used to visualize slices of the genome adjacent to the identified MAT loci. All the lung isolates including DCF04/Ex4 *E. dermatitidis* isolate encoded a MAT1-1 locus (**Figure 2, Supplemental Figure 2, Supplemental Table 5**). For all other *E. dermatitidis* isolates and *Capronia* species analyzed, *SLA2*, a SRC-like adapter protein, and *APN2*, apurinic-apyrimidinic endonuclease 2, flanked the MAT loci in all isolates, and a hypothetical protein between SLA2 and MAT1-1-4 was also present across isolates. The MAT locus found in strain NIH/UT8656 have the MAT1-2 mating type, while our clinical isolates have the MAT 1-1 mating type. Consistent with MAT loci described in *E. dermatitidis* (Metin *et al*. 2019) DCF04 had both MAT1-1-4 and MAT1-1-1 genes (**Figure 2**). The homothallic outgroup *Capronia coronata CBS617.96* genome (Teixeira *et al*. 2017) contains both MAT1-1 and MAT1-2 genes.

**Figure 2.**
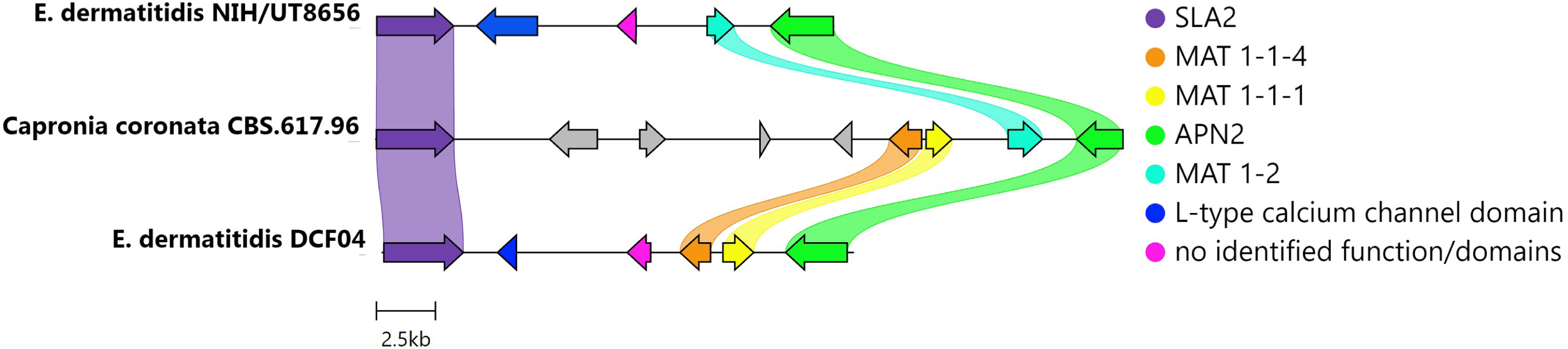
Mating-type determination of 4 clinical isolates of *E. dermatitidis.* Gene content, order and orientation of the MAT locus from *E. dermatitidis* DCF04 and NIH/UT8656 strains and the Chaetothyriales black yeast *Capronia coronata CBS 617.98*. The locus is flanked by two genes SLA2 (purple) and APN2 (green). The MAT 1-1 genes, MAT 1-1-4 (orange) and MAT 1-1-1 (yellow) are observed in DCF04 strains while the reference strain NIH/UT8656 has a MAT 1-2 (teal) gene. In addition, two genes are predicted in the interval between SLA2 and the MAT genes. One is a L-type calcium channel domain (blue), which is only found in the *E. dermatitidis* strains, and a gene with no identified function or domains (pink) which is syntentically adjacent to the MAT genes in all strains and species.

### Chromosome copy number variation across E. dermatitidis isolates

Copy number variation of full or partial chromosomes was evaluated by calculating depth of coverage using 10kb sliding windows (**Figure 3).** The read depths of windows across *E. dermatitidis* isolate genomes from this study were compared across all 9 chromosomes. Visual scanning of the plots identified an anomaly of at least 1.5x higher coverage on chromosome 5 in Ex3 (**Figure 3A**). A similar but much smaller region of chromosome 5 appears to have 2-2.25x coverage and may be duplicated as an aneuploid in Ex13. Additional partial 1.25-1.5x coverage for part of the left arm of chromosome 2 in isolate Ex18 is also observed.

**Figure 3.**
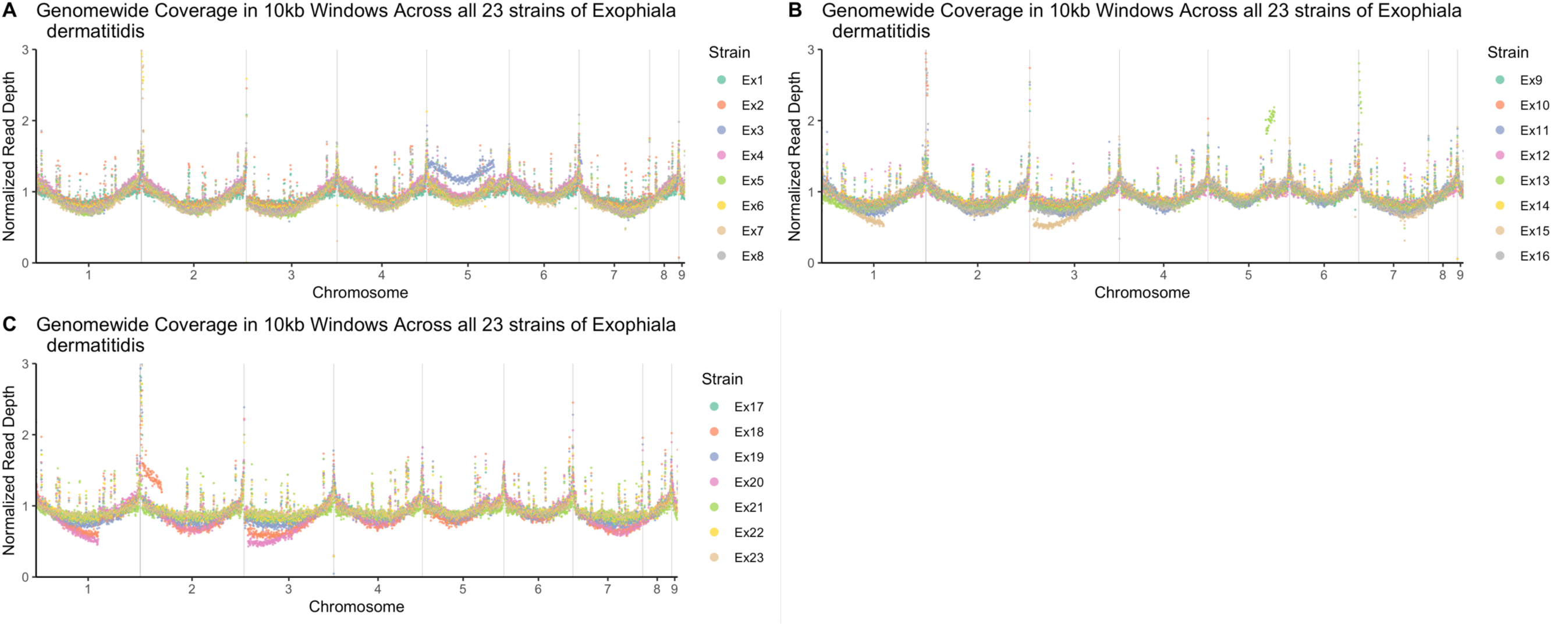
Genome sequencing depth coverage visualization to test for copy number variation across isolates. Visualization of depth of coverage was generated by plotting a normalized read depth across chromosomes for isolates **A)** Ex 1-8, **B)** Ex9-16, and **C)** Ex17-23.

The plots of isolates Ex15, Ex18 and Ex20 indicate 0.5x less normalized coverage in chromosomes 1 and 3 (**Figure 3B and Figure 3C**), a possible sign of segmental chromosomal aneuploidy event. The continuation of the coverage pattern between chromosomes 1 & 3 seem to complement each other for these isolates. Haploid organisms, like *Exophiala dermatitidis*, could benefit from genomic plasticity through expansion or contraction to enable adaptation in a new environment (Selmecki *et al*. 2010; Legrand *et al*. 2019). Noting that Ex3 was isolated in the early time point, this genome copy may contribute to an adaptive mechanism to support colonization and persistence in the human host. Other isolates sharing potential aneuploidies (Ex15, Ex18, Ex20) are from the late time point and also have the highest mutation rate among this population as observed from the phylogenetic tree and mutation rate calculation (**Figure 4 and Table 4**).

**Figure 4:**
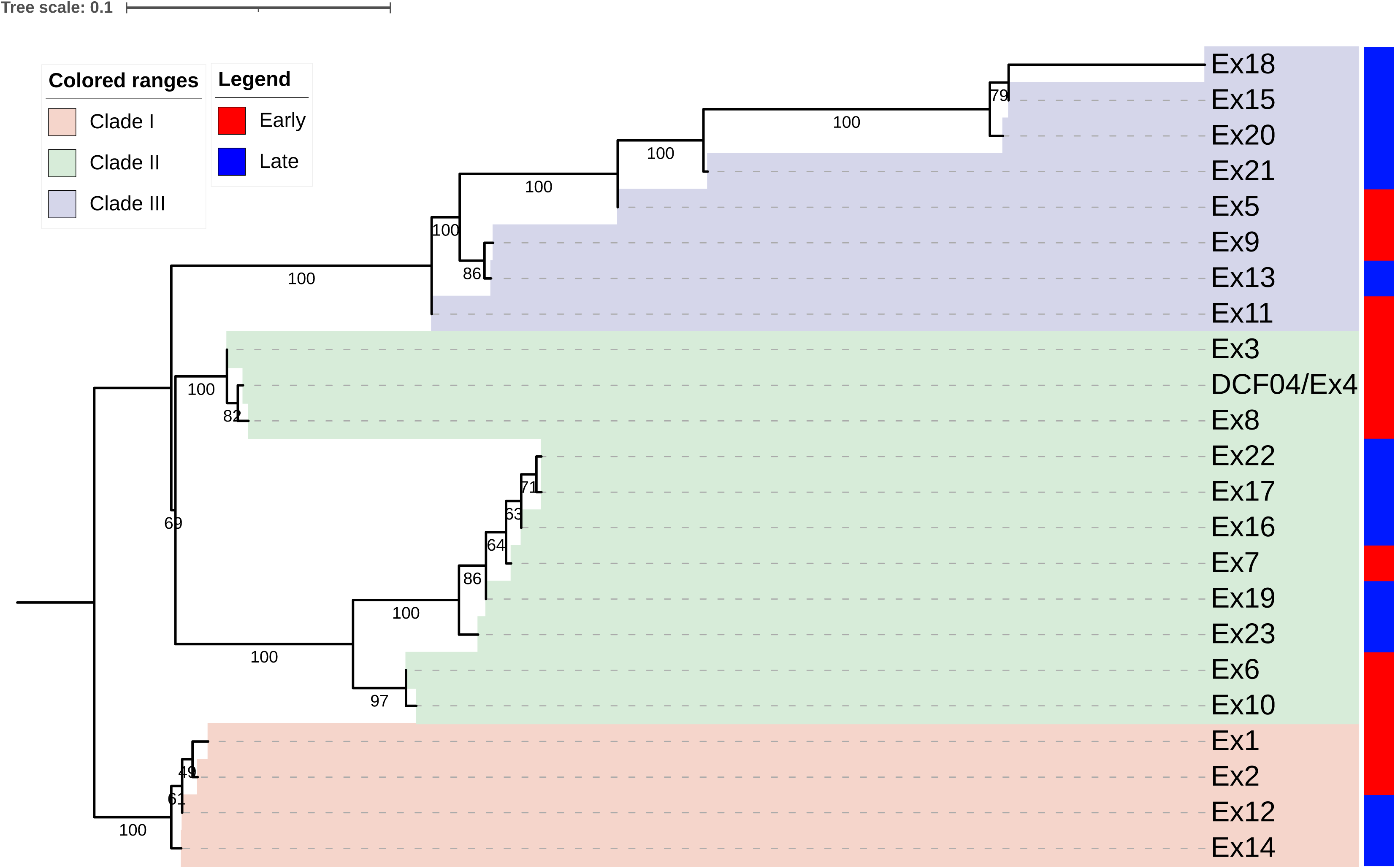
Phylogenetic tree of 23 isolates. A Maximum-Likelihood phylogenetic tree constructed from the Single Nucleotide Variants by IQTREE2 identified from the isolate resequencing data. Tree is rooted with the Clade I branch based on additional analyses that included NIH/UT8656 as an outgroup. Isolates are labeled as one of three clades based on the phylogenetic relationships and the isolation time point is indicated with a red (early) or blue (late) colored box.

**Table 4.**
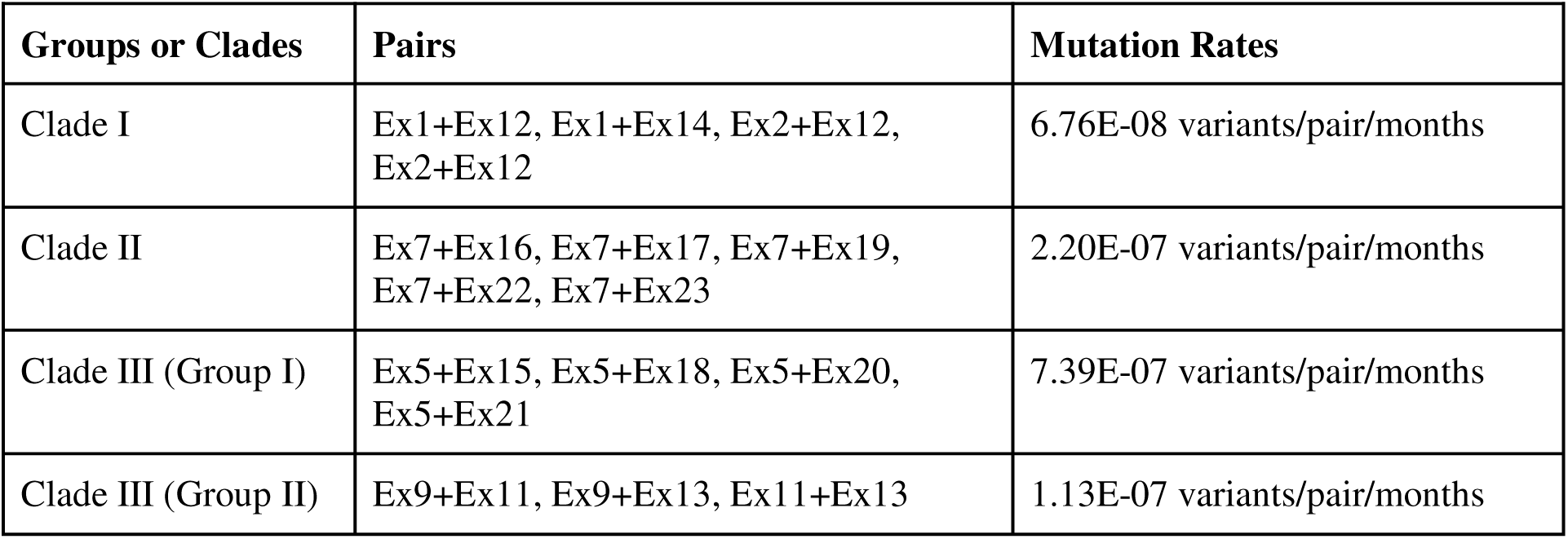
Average mutation rates for each clade/group. Mutation rates calculated for each clade of isolates and two sub-groups of Clade III. Calculations were summarized as the median of all pairs of isolates within a group or clade.

### SNP genotyping and SNP-based phylogenetic analysis of 23 E. dermatitidis isolates

We used the DCF04/Ex4 isolate as a reference for variant identification within the 23 CF isolate collection. The DCF04/Ex4 VCF file generated 441 variants between twenty-two isolates and the DCF04 reference isolate. A phylogenetic tree was generated using the filtered SNP data results and rooted with the earliest diverging group containing Ex1, Ex2, Ex12 and Ex14 (**Figure 4**). When repeated for out-population reference strain NIH/UT8656 the analysis resulted in about ∼11,000 variants between the NIH/8656 strain and the CF isolates (Kurbessoian 2022). Another phylogenetic tree was generated using a secondary dataset created with the NIH strain to be used as a root (**Supplemental Figure 4**).

Three clades (Clade I, Clade II, and Clade III) were identified based on the tree. Clade I is composed of two early collected and two late collected isolates of *E. dermatitidis* Ex1, Ex2, and Ex12, and Ex14. Clade II is composed of three subgroups, one of which contains only early isolates and a second group with both an early isolate and late isolates. The third group in Clade II contains two early collected isolates Ex6 and Ex10. Clade III has two main groups composed of both early and late isolates. The subgroup of Clade III had a long branch length suggesting more divergence. Interestingly, the CNV plot (**Figure 3**) showed these three isolates, Ex15, Ex20 and Ex18 contained similar CNV differences in chromosome 1 and 3 when compared to the other isolates.

### Non-synonymous and synonymous SNP and INDEL pairwise differences

A dissimilarity matrix was constructed comparing the overlaps of SNPs and INDELs collated for all pairs of isolates. As expected, isolates that are closest to each other in the SNP-based phylogenetic tree had fewer differences in their SNP composition (**Figure 5A**). Our analysis was further supported by the observation that the number of SNP differences correlated with the number of INDELs detected in pairwise analyses (**Figure 5B**). Within Clade I and Clade II, isolates had few SNP and INDEL differences, indicating the evolutionary distance between them was small. The SNP and INDEL counts within Clade III were much higher indicating a higher divergence within this group of isolates as compared to other groups.

**Figure 5.**
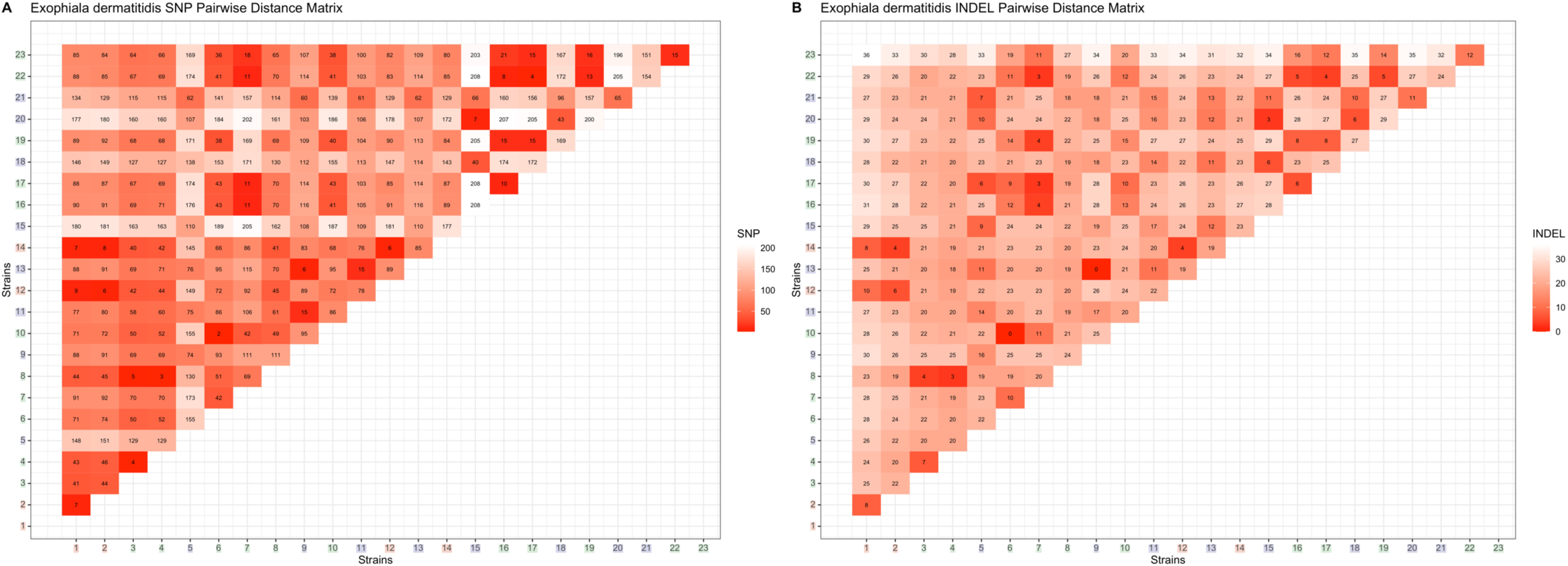
Matrix of *E. dermatitidis* CF isolates SNP and INDEL pairwise dissimilarities. The number of SNPs **(A)** and **(B)** INDELs that differ among pairs of isolates. The more similar isolate pairs have lower (darker red) numbers and the color approaches white indicating more dissimilar isolates. Clade I and Clade II isolates generally differ by very few SNP and INDELs (noting a few exceptions) consistent with their inferred near phylogenetic relationships. Within Clade III isolate pairs differ by more SNPs which may indicate a higher mutation rate within these isolates.

### Non-synonymous SNP Differences seen between clades

The human-readable snpEff table which best described the variant analysis along with functional gene annotation can be found in **Supplemental Table 6**. Clade I is composed of four isolates: two (Ex1, Ex2) and two isolates from the late time point (Ex12, Ex14). Two genes with mutations are of note, a MADS-box transcription factor (HMPREF1120_06786) and a G4 quadruplex nucleic acid binding protein (HMPREF1120_02174). The MADS-box transcription factor is part of the MADS-box proteins with a highly conserved 56 amino acid DNA-binding domain, some containing a weakly conserved K-box domain that is involved in the dimerization of transcription factors (Shore and Sharrocks 1995). In fungi, isolates with knocked-out MADS-box genes have reduced virulence (Damveld *et al*. 2005; Qu *et al*. 2014; Xiong *et al*. 2016). Mutations in *E. dermatitidis* MADS-box regions may increase their pervasiveness and tolerance of the lung environment. Previous studies have found that G4 quadruplex nucleic acid binding proteins. The helical complex is formed through guanine rich nucleic acid sequences and is found at the telomeric regions of chromosomes.

Clade II contains 11 isolates, six from the early time point and five from the later time point, that fall into three different subgroups. Ex3, Ex4/DCF04, and Ex8 fell into one group, Ex22, Ex17, Ex16, Ex7, Ex19 and Ex23 fell into the second group, and Ex6 and Ex10 formed a third group. Group one and group three both contain only early isolates, while the second group contains the majority of the later isolates.

Clade III, which contains eight isolates, also divided into two groups: one composed of Ex18, Ex15, Ex20, Ex21, and the other group of Ex9, Ex13 and Ex11. Five are later isolated while the other three are from the early isolations. The most significant group is the second consisting of Ex9, Ex13 and Ex11. Four genes were found to have distinct mutational differences when comparing Ex13 (a late isolate) to the early isolates Ex9 and Ex11. These four genes are the Ap-3 complex subunit delta (HMPREF1120_00143), DNA repair protein RAD50 (HMPREF1120_05599), a regulator of nonsense transcripts 1-like protein (HMPREF1120_06837) which is similar to helicase-RNA complex, and a gene with no identified function (HMPREF1120_02556). The second group had three isolates (Ex18, Ex15 and Ex20) with a higher mutation rate than *E. dermatitidis* in Clade III. There are about 13 instances of hypothetical proteins, while the other 12 instances are predicted genes. Only six of the twelve are genes of note: an MFS transporter DHA2 family methylenomycin A resistance protein (HMPREF1120_00012), sulfite reductase (ferredoxin) (HMPREF1120_00943), glycerol ethanol-ferric requiring protein (HMPREF1120_01007), polyketide synthase (HMPREF1120_03173), MFS transporter SP family solute carrier family 2 (HMPREF1120_06771) and a DNA repair protein RAD50 (HMPREF1120_05599). Research on polyketide synthases in micro-colonial fungi and *E. dermatitidis* have been found to impact phenotypes and adjust the melanin synthesis pathway, resistance or susceptibility to antifungals and extreme environment adaptability (Paolo *et al*. 2006).

### Analysis of SNP differences between clades

Finally, a single non-synonymous mutation in a gene orthologous to *S. cerevisiae MRS4* (HMPREF1120_06597), a mitochondrial iron transporter. One allele of *MRS4* allele encoded a protein that was identical in sequence to the allele in the reference NIH strain UT8656, present in the isolates in Clade I (Ex1, Ex2, Ex12, Ex14), and a subclade of clade II (Ex3, Ex4/DCF04, Ex8). The remainder of the isolates had a second allele with a non-synonymous mutation in the 40^th^ amino acid position, converting a glutamic acid residue to glycine. The functional consequences and differences of these Mrs4 alleles will be described in a separate manuscript.

### Testing for enrichment of evolutionary patterns within clades

A pairwise comparison of the synonymous and nonsynonymous SNP differences was performed on fifteen pairs of isolates identified as early and late members of the same lineage. **Table 3** summarizes the significant results with a focus on candidate genes that may relate to lung pathogenicity. A comprehensive list of gene differences among all pairwise comparisons is in **Supplemental Table 7** and includes hypothetical proteins with no identified function. This analysis tested for differences between early and late isolates found in the same clade to focus on changes that may have occurred within the host.

The analysis considered all pairwise combinations between the early Ex1 and Ex2 and the late Ex12 and Ex14 isolates from Clade I. These isolates show very little genetic differentiation (about 6-9 variant SNPs and 6-10 variant INDELs). The contrast of early Ex1 vs Ex2 isolates to the late isolates (Ex12,Ex14) found variants in three genes: AFG2-an ATPase (HMPREF1120_06104), a DNA polymerase (HMPREF1120_07994), GYP1-a GTPase activating protein (HMPREF1120_08601), and MFS transporter (HMPREF1120_04157) homologs. These variants are an interesting observation and warrants extra analysis in the future.

Pairwise comparisons of the Clade II isolates contrasted the early Ex7 with each of the four late isolates Ex16, Ex17, Ex19, and Ex23. There was more variation among these isolates (about 11-169 variant SNPs and 3-16 variant INDELs) than observed in Clade I isolates. Isolate Ex23 also appeared to have substantially more differences from all others indicating it may be more distantly related or where its corresponding ancestral early strain was not sampled. A variant of note, NAD-dependent histone deacetylase SIR2, is involved with chromosomal remodeling specifically with phenotype transcription modification (Freire-Benéitez *et al*. 2016).

Comparison of Clade III group 1 members Ex5 (early) to Ex15,18,20,21 identified the highest number of variants as compared to the other clades (from 62-138 variant SNPs and 7-10 variant INDELs). Clade III group 2 members Ex9 (early), Ex11 (early) and Ex13 had a similar number of variants as Clades I & II (6-15 variant SNPs and 0-17 variant INDELs). The five pairwise comparisons for group 1 revealed a non-synonymous mutation in the RAD50-DNA repair protein (HMPREF1120_05599). Group 1 members in the phylogenetic tree also appear to have a long branch length indicating a potentially higher diversification rate than the other clades. It may be that the RAD50 changes could have an impact on DNA repair and contribute to the higher mutation rate observed. In addition, non-synonymous mutations were identified in a cytochrome P450 oxidoreductase (HMPREF1120_01361), MFS transporter sugar to H+ symporter (HMPREF1120_04157) and GTPase activating proteins (HMPREF1120_08601). SNP variants with predicted functional impact on interactions with the host and CF lungs were found in the sulfite oxidase (HMPREF1120_01306), sulfite reductase (HMPREF1120_00943), polyketide synthase (HMPREF1120_03173), and tyrosinase (HMPREF1120_04514) genes.

Interestingly, the group 2 (Clade III) isolates Ex9, Ex11 and Ex13 had more variants in iron-binding (HMPREF1120_06751) and siderophore related (HMPREF1120_07838) genes than group 1. We took a candidate gene approach and tested if the siderophore NPRS SidC (HMPREF1120_07636) had accumulated any specific mutations in these lineages, but we failed to identify any non-synonymous variants in this gene across the CF lung isolates. Though, the presence of SNP variants in iron-binding and siderophore transporters indicates there are other means *E. dermatitidis* is obtaining the elusive iron molecules from its environment. Further analysis of these variants will point to supporting phenotypic differences allowing for effective colonization of CF lung environments.

### Mutation rate calculation to test for different rates of evolution between clades

We calculated a mutation rate for each of the three clades (**Table 4**) by taking an average of all the pairwise comparisons with a clade. The Clade I average of the four pairwise calculations for all combinations of early and late isolates was 6.78E-08 variants/pair/2 years. The average rate for the five comparisons of early to late isolates in Clade II was 2.20E-07 variants/pair/2 years. The average rate for the six comparisons of early to late isolates in Clade III the first group values were averaged and determined to be 7.39E-07 variants/pair/2 years, while the second group values were averaged and determined to be 1.13E-07 variants/pair/2 years. The Clade III group 1 isolates appear to be a faster evolving group and may have acquired variants allowing improved adaptation to the lung environment. One-way ANOVA test revealed a statistically significant difference between all Clades (F-value = 7.0711, p-value = 0.00104) and a Tukey post-hoc test across the clades indicated Clade III (Group 1) has a significantly different mutation rate. Calculations underlying the mutation rate values are detailed in **Supplemental Table 8**.

### Heterogeneous phenotypes observed in the E. dermatitidis isolates

The twenty-three *E. dermatitidis* isolates (Ex1-11 from the early time point and isolates Ex12-23 from the late time point) were heterogeneous for traits in a number of ways including pigmentation, antifungal sensitivity, and auxotrophy. We noted differences in melanin pigmentation across isolates (**Figure 6A**) that did not correlate with either time point or phylogenetic clade. Within Clade 1, Ex1, Ex2, and Ex12 had light melanization, but isolate Ex14 was one of the strongest melanin producers (**Figure 6B**). In a search for genetic determinants responsible for this phenotypic range, we identified three candidate intergenic mutations. Of particular interest is a mutation in the 5’ UTR of a putative iron-dependent biphenyl-2,3-dioxygenase potentially involved in the degradation of phenolic compounds. Melanin is synthesized through the oxidation of various aromatic compounds, and thus disruption or enhancement of biphenyl oxidation may be a mechanism by which melanin production was enhanced in Ex14.

**Figure 6.**
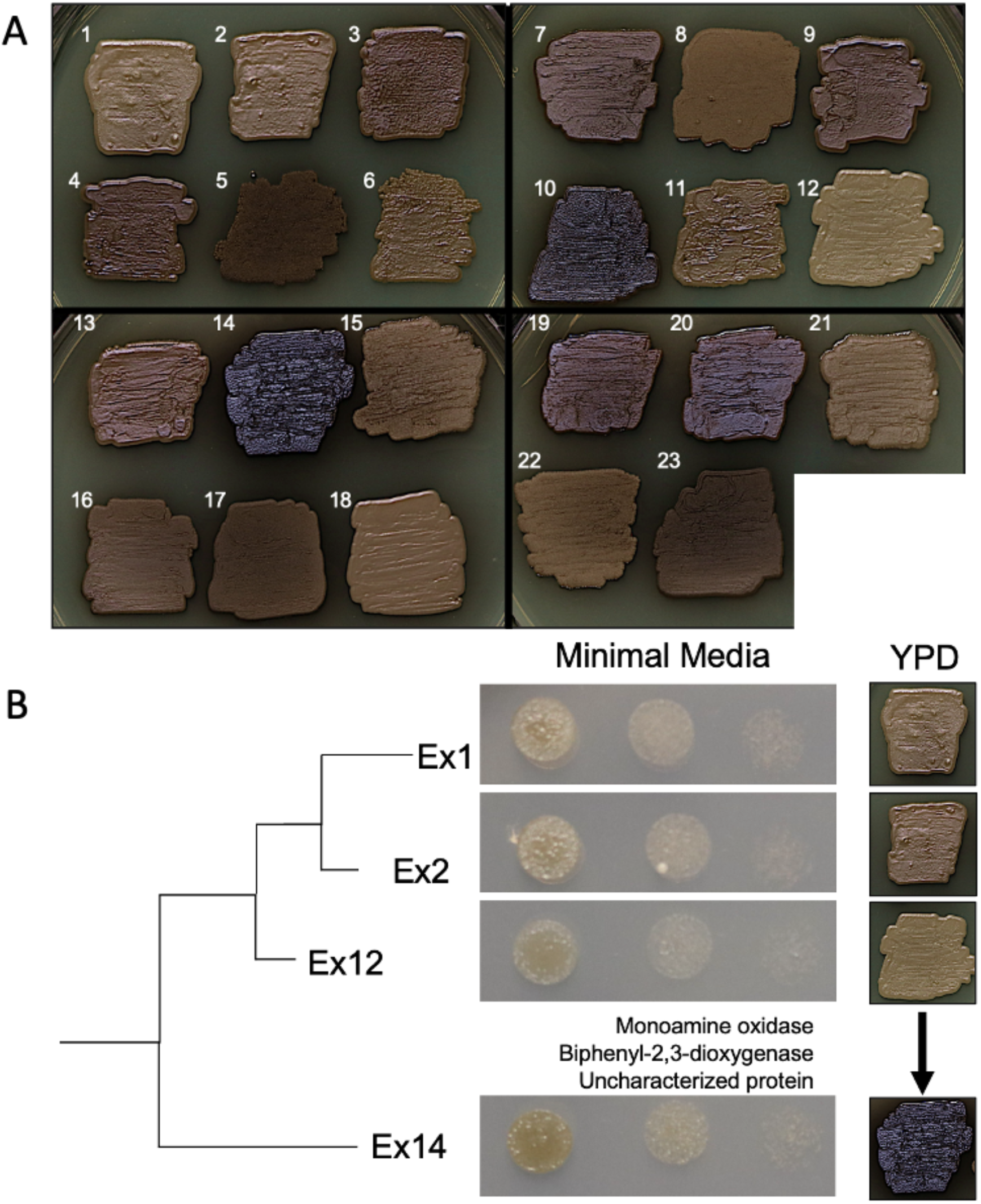
*E. dermatitidis* clades and their comparisons to melanin phenotype and MIC. **(A)** Isolates were struck from freezer stock onto YPD plates and imaged at 120 h at 37°C to compare melanin accumulation. **(B)** Strong differences in melanin production between related isolates may be due to intergenic mutations in the 5’ UTR of a biphenyl-2,3-dioxygenase encoding gene.

The 23 isolates also varied in sensitivity to itraconazole, a recommended treatment for *E. dermatitidis* infections (Fothergill *et al*. 2009; Mukai *et al*. 2014), over a ∼10-fold range (minimum inhibitory concentrations (MIC) from 0.0625-0.5 µg/ml) (**Figure 7**). Heterogeneity in amino acid auxotrophy, scored as no growth on a minimal medium that was rescued by supplementation of amino acids (**Figure 7**), was also observed. Lastly, there were stable differences in filamentation across isolates in two of the three clades; closely related strains did not have similar morphologies (**Figure 8**).

**Figure 7.**
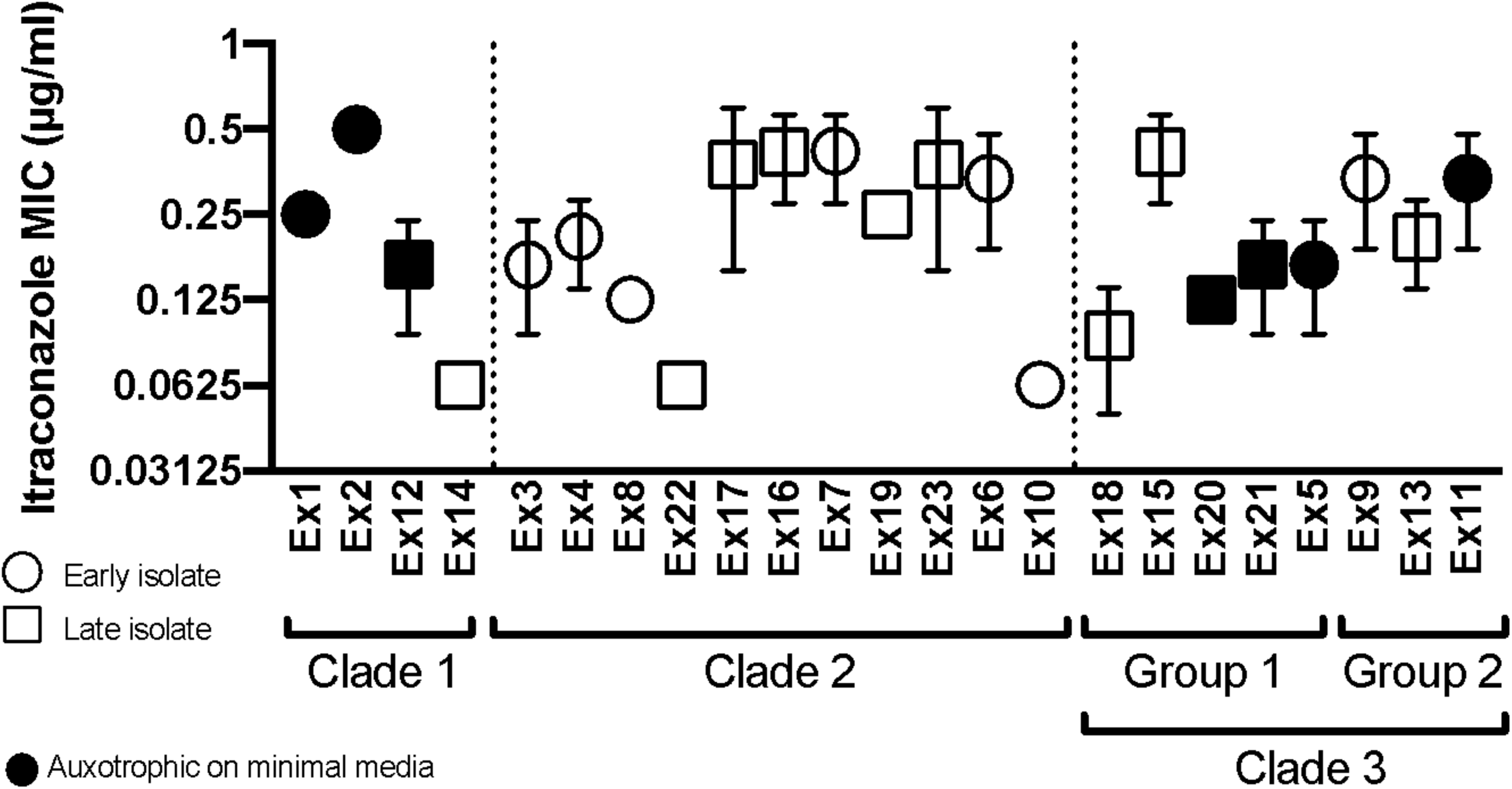
Itraconazole MIC testing on 23 isolates of *E. dermatitidis,* while comparing clades. Isolates were grown for ∼24 hours in liquid YPD at 30°C in a rolling barrel culture, then inoculated into 96-well flat bottom plates at a concentration of 1000 CFU/well, and allowed to grow for 72 hours at 37° before determining MIC, which is represented on the figure as the mean of three biological replicates. For determination of auxotrophy, cells were grown as described above, CFU equilibrated, and spotted onto rich media and minimal media (YNB) with and without the addition of casamino acids. isolates with decreased growth on YNB relative to YPD, and could be rescued with the addition of amino acids are noted as auxotrophic.

**Figure 8.**
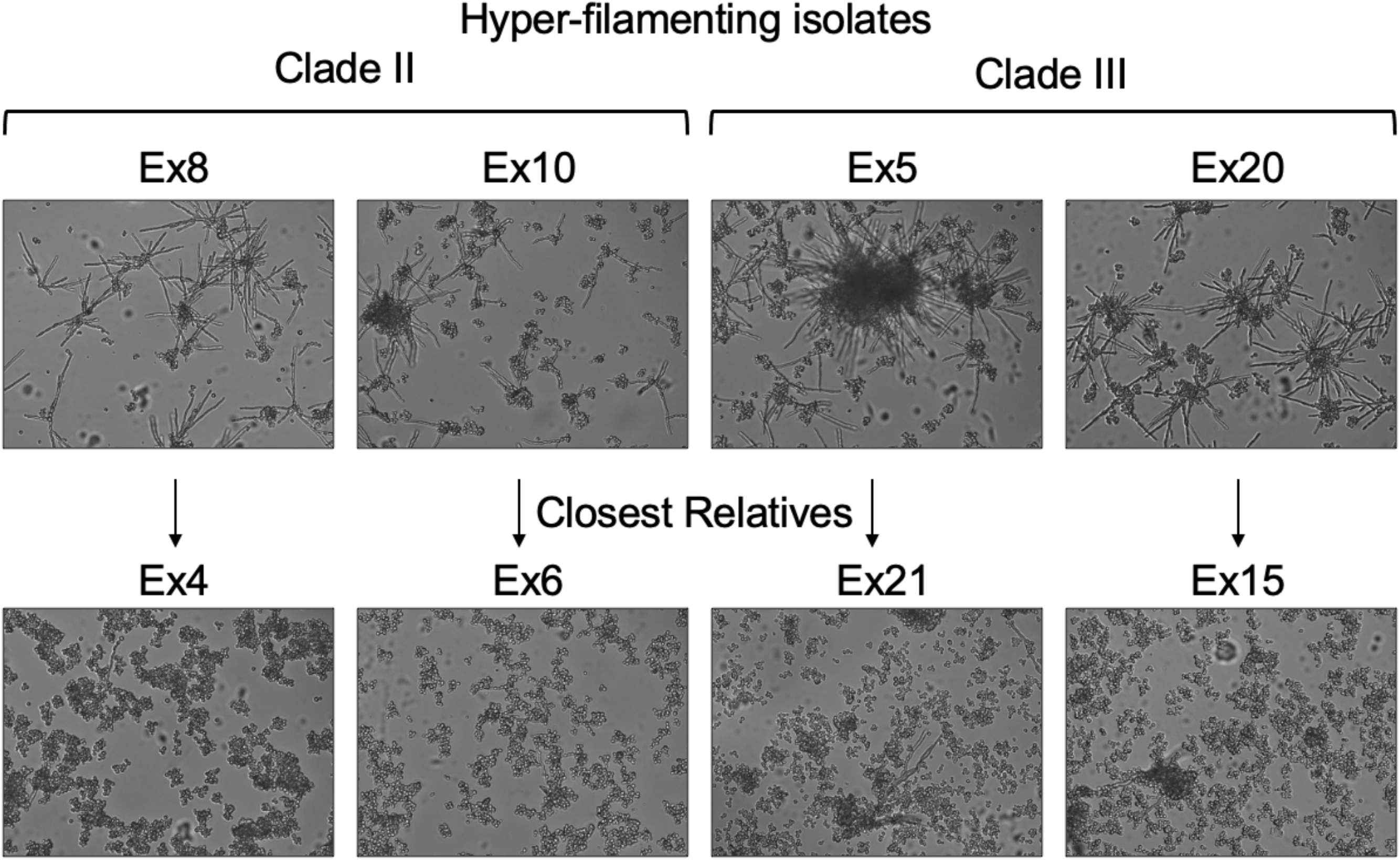
Cell phenotype microscopy. Isolates were grown for 18 h in RPMI, at 37° in 5% CO, and imaged at 63X using DIC microscopy_2_. The first row of images depicts isolates displaying hyper-filamentous attributes, and are distributed throughout their respective clades. The second row depicts the closest related isolate to each respective isolate in the first row, which lack similar phenotypes.

## Discussion

### Fungus-dominant population

While opportunistic fungal pathogens are often culturable from the sputum of patients with cystic fibrosis, they do not commonly present as the dominant microbe or as a risk for infection. In this paper we have presented a clinical case where a population of the black yeast *E. dermatitidis* was the predominant microbe concomitant with a lung exacerbation event. Other reported clinical cases have shown that *E. dermatitidis* has an underappreciated role as a CF pathogen (Haase *et al*. 1990; Kusenbach *et al*. 1992; Kondori *et al*. 2014). Previous sputum isolates revealed the presence of this uncommon fungi two years prior, indicating that it persisted in low concentrations before having the opportunity to dominate the lung microbiome. Because *E. dermatitidis* is relatively slow growing (Rath *et al*. 1997; Sudhadham *et al*. 2011; Malo *et al*. 2021), it may be easily missed during routine clinical microbiological identification. To better understand the CF disease and the fungal infections associated with a long-term disease such as this one, it is important to consider testing for slow growing microorganisms such as black yeasts.

### Phenotypic heterogeneity; Melanin production and drug resistance

Identification of key evolutionary strategies is crucial to the understanding of microbial pathogenesis in clinically relevant settings. Developing methods for evaluating population structure and heterogeneity for these human-associated microbes will aid in future studies. Initial observations of these isolates indicated a striking amount of heterogeneity in melanin production, which propelled an investigation into the genotypic diversity of the population. Population heterogeneity is an important factor to consider in treatment, and determining adaptive responses through genomics can help identify selective stimuli in chronic infections (Demers *et al*. 2018). We expected that differing levels of melanin, which comprises part of the cell wall in black yeasts, would have an effect on antifungal resistance. While drug resistance to clinically relevant antifungals Itraconazole and Voriconazole varied slightly between isolates, there was no consistent pattern of evolution that corresponded either with time of isolation or melanin production (**Figure 7**). We found amino acid auxotrophs in the population, and the isolation of auxotrophs in CF infections has been previously described in bacterial isolates, such as *P. aeruginosa* (Barth and Pitt 1995; Thomas *et al*. 2000). This repeated occurrence of filamenting phenotypes in phylogenetically diverse isolates may indicate evolution towards hyper-filamentation or a non-mutation-based difference between isolates such as “switching” or phase variation, or epigenetic regulation.

### Short and long read sequencing, in-population vs outside reference

While short-read sequencing provided us with the basic and high quality read depth for each CF isolate, using the Oxford Nanopore long-read technology sequencing to build on top of a short read assembly provided us with a well assembled genome. This genome was then used as an in-population reference allowing for a better recovery of variants due to the read mapping to a closer isolate. When using our in-population reference, compared to the NIH/UT8656 strain (Robertson *et al*. 2012; Chen *et al*. 2014; Schultzhaus *et al*. 2020; Malo *et al*. 2021), our quantification of population specific number of variants is higher (due to higher sensitivity) while still maintaining relevance in our study system.

### Temporal resolution of variation accumulation in a fungal CF lung population

Contrasting mutations accumulating in isolates collected from two time points allowed comparison in mutation rate differences among three different genetic lineages of *E. dermatitidis* and evaluated if any changes could suggest adaptations that enabled lineages to survive the lung environment. Average mutation rates within a Clade tended to be similar indicating common diversification events. Statistical comparisons of mutation rates between Clades indicate significance seen between Clade I and Clade III group 1, Clade II and Clade III group 1 and Clade III group 1 and group 2. The main difference here is the greatly increased diversification in Clade III group 1 which is shown with the phylogenetic tree in Figure 4, chromosomal aneuploidies seen in Figure 3, the very high SNP and INDEL counts seen in Figure 5, the increased mutation rates seen in Table 4 and hyper-filamentous phenotypes see in Figure 8. We propose the mutations found in *RAD50* may have contributed to the increased diversification seen in this group and future studies will test this hypothesis (Goldman *et al*. 2002; Krogh and Symington 2004).

These clades persisted over a two-year span. Are there physical separations disallowing for SNPs in these sub-populations to mix? When observing the functional assessment of variants unique to each clade, we identified alleles in genes that encode for transporters, cytochrome P450 oxidoreductases, iron acquisition and DNA repair processes. Iron acquisition can be a possible virulence method *E. dermatitidis* could be using to persist in the lung environment as seen in *Aspergillus fumigatus* and *Pseudomonas aeruginosa* (Neilands 1995; Matilla *et al*. 2007; Schrettl *et al*. 2007). Mutations in transporters could be an evolutionary move to adapt to antibiotics or antifungal treatments. The finding of a non-synonymous mutation, *MRS3/4* a mitochondrial iron transporter, may also help point into the pathogenicity and virulence (Murante, 2022, In preparation). Though, the presence of certain SNP variants in iron-binding and siderophore transporters suggests that there could be other means by which *E. dermatitidis* is obtaining the elusive iron molecules from its environment. It would be important to observe the type of variants (e.g. stop codons or a loss of function) produced in these iron-related genes to begin searching for other virulence genes. Further analysis of these variants will point to supporting phenotypic differences allowing for effective colonization of CF lung environments. It’s clear that this mutation is not found in the rapidly diverging Clade III and only in the Clade I cluster with very few variants. The isolates we have sequenced only contain one mating type, indicating the low likelihood that even if the isolates have close proximity to each other they are unable to mate. As these isolates were collected from sputum plates, it could be certain isolates have co-localized within different lobes of the lung and could have nutritional or other ecological partitioning that could be driving these clades to diverge and remain diverged. Our results suggest that there may have been a diversification event that occurred early perhaps during the initial colonization. The clades then stabilized once co-localizing into their respective niches, or there may have been a second ‘inoculation’ event from a common, stably heterogeneous environmental source at some point in the two year timeline (Warren *et al*. 2011).

### Conclusions

As it has become increasingly clear, collecting a single strain and using it as a metric to assess a single environment and moment of time is inaccurate and a more population approach must be used to best assess microbial infection (Demers *et al*. 2018). Having a closer reference strain to assess variants is also necessary to observe true variants and not a result of years of strain formation. Our results indicate the CF lung environment supports stably diverged populations of clonally derived yeasts.

## Supporting information

Supplemental Figure 1. Genome dot plot of E. dermatitidis DCF04 and E. dermatitidis NIH/UT8656.

Supplemental Figure 2. Mating-type determination of 23 clinical isolates of E. dermatitidis.

Supplemental Figure 3. Phylogenetic tree of 24 isolates

Supplemental Table 1. Collection and MIC values for CF patient derived E. dermatitidis isolates.

Supplemental Table 2. Genome assembly statistics for 23 CF isolates.

Supplemental Table 3. Telomere Recovery for DCF04.

Supplemental Table 4. OrthoFinder summary comparing DCF04 and NIH/UT8656.

Supplemental Table 5. Mating-type determination locus name descriptions of E. dermatitidis.

Supplemental Table 6. Functional impact of identified variants.

Supplemental Table 7. All functional SNP and INDEL results for early and late pairs.

Supplemental Table 8. All 23 mutation rates calculated.

## Data availability

Sequence data generated for isolates with Illumina and Oxford Nanopore technology are deposited in NCBI Sequence Read Archive linked under BioProject PRJNA628510. The assembled genomes of each CF isolate (Ex1-23) are available under accessions listed in **Supplemental Table 2**. The assembled and annotated genome of the in-population reference isolate (DCF04) is available at accession JAJGCF000000000. All analysis pipelines, custom scripts used for data analysis, and raw variant data in the variant call format are available in Github repository https://github.com/tania-k/CF_Exophiala_dermatitidis and archived in Zenodo (Kurbessoian 2022).

## Acknowledgments

This work was supported by NIH grants R01 AI127548 (D.A.H. and J.E.S.) and T32 AI007519 (D.R.M.). JES was partially supported by R01 AI130128. This project was also supported by DartCF (NIH grant P30-DK117469) and the Cystic Fibrosis Foundation Research Development Program (STANTO19R0). Computational analyses were performed at the University of California-Riverside HPCC supported by grants from the National Science Foundation (MRI-1429826) and NIH (S10OD016290). JES is a CIFAR Fellow in the Fungal Kingdom: Threats and Opportunities program.

---

**Supplemental Figure 1. Genome dot plot of *E. dermatitidis* DCF04 and *E. dermatitidis* NIH/UT8656.** D-GENIES web application was used to generate a dot plot representation to compare genome assembly content and test for structural changes or rearrangements. Both iterations of the plot indicate a majority of the genomes are nearly identical with each other.

**Supplemental Figure 2. Mating-type determination of 23 clinical isolates of *E. dermatitidis.*** Gene content, order and orientation of the MAT locus from 23 CF isolated *E. dermatitidis*, NIH/UT8656 strain and the Chaetothyriales black yeast *Capronia coronata CBS 617.98*. The locus is flanked by two genes SLA2 (purple) and APN2 (green). The MAT 1-1 genes, MAT 1-1-4 (orange) and MAT 1-1-1 (yellow) are observed in all 23 CF strains while the reference strain NIH/UT8656 has a MAT 1-2 (teal) gene. In addition, two genes are predicted in the interval between SLA2 and the MAT genes. One is a L-type calcium channel domain (blue), which is only found in the *E. dermatitidis* strains, and a gene with no identified function or domains (pink) which is syntentically adjacent to the MAT genes in some strains.

**Supplemental Figure 3. Phylogenetic tree of 24 isolates.** A Maximum-Likelihood phylogenetic tree constructed from the Single Nucleotide Variants by IQTREE2 identified from the isolate resequencing data. Tree is rooted with NIH/UT8656 as an outgroup. Isolates are labeled as one of three clades based on the phylogenetic relationships and the isolation time point is indicated with a red (early) or blue (late) colored box.

**Supplemental Table 1. Collection and MIC values for CF patient derived *E. dermatitidis* isolates.** Information on the 23 CF isolates cultured from one patient sputum across three years. The table summarizes phylogenetic clade designation, itraconazole MIC, date of collection, and classification as an Early or Late.

**Supplemental Table 2. Genome assembly statistics for 23 CF isolates**. Assembly statistics for scaffolded assemblies of isolates and NIH/UT8656 previously published genome. Table indicates assembly summary statistics of the assembly using Illumina sequencing of all 23 CF isolates. Summary statistics of scaffold lengths are presented with genome completeness data calculated with BUSCO using the ascomycota_odb9 database.

**Supplemental Table 3. Telomere Recovery for DCF04.** Table depicting telomere recovery results for DCF04 CF *E. dermatitidis*. Candidate telomeric repeat units “TTTAGGG/CCCTAA” were identified as repeat arrays at both ends of five scaffolds, but also found as single pairs in the remaining 4 scaffolds, as would be expected for 9 complete chromosomes.

**Supplemental Table 4. OrthoFinder summary comparing DCF04 and NIH/UT8656.** Comparison of the shared and unique orthogroups found in the annotated proteomes of *E. dermatitidis* strains NIH/UT8656 and DCF04. A majority of orthogroups (8,256; 99%) had members from both strains, of these 15 were single-copy orthogroups containing a single protein-coding gene from each strain. There were 10 orthogroups unique to DCF04 encompassing 34 protein-coding genes and 5 orthogroups unique to NIH/UT8656 made up of 24 protein-coding genes. DCF04 contained 705 unassigned genes, while NIH/UT8656 had 475.

**Supplemental Table 5. Mating-type determination locus name descriptions of *E. dermatitidis*.** This table describes all the proteins depicted in Figure 2 of the gene content, order and orientation of MAT locus from *E. dermatitidis* isolates in this study, strain NIH/UT8656 along with the Chaetothyriales black yeast *Capronia coronata CBS 617.98*.

**Supplemental Table 6. Functional impact of identified variants.** Filtered human readable snpEff tabular results for all CF 23 *E. dermatitidis*. Annotations have been added to the final list to better help assess the function of each protein while also observing variants detected from GATK.

**Supplemental Table 7. All functional SNP and INDEL results for early and late pairs.** Results indicated in this table include all hypothetical or undescribed results along with results described in Tables 2 and 3. Proteins listed include Protein ID numbers to better facilitate identification.

**Supplemental Table 8. All 23 mutation rates calculated.** Mutation rates for each 23 CF *E. dermatitidis* isolates calculated using formula described in methods. One-way ANOVA was run to detect significance (p-value= 0.00104, F-value = 7.0711) along with Tukey multiple comparisons of means indicating Clade III had the highest significance among the six pairwise comparisons.

## Notes

https://doi.org/10.5281/zenodo.7106110

https://github.com/tania-k/CF_Exophiala_dermatitidis

### Competing Interest Statement

The authors have declared no competing interest.

