## Supplementary figures and images for "In host evolution of *Exophiala dermatitidis* in cystic fibrosis lung micro-environment"

### Supplemental Figure 1. Genome dot plot of E. dermatitidis DCF04 and E. dermatitidis NIH/UT8656.

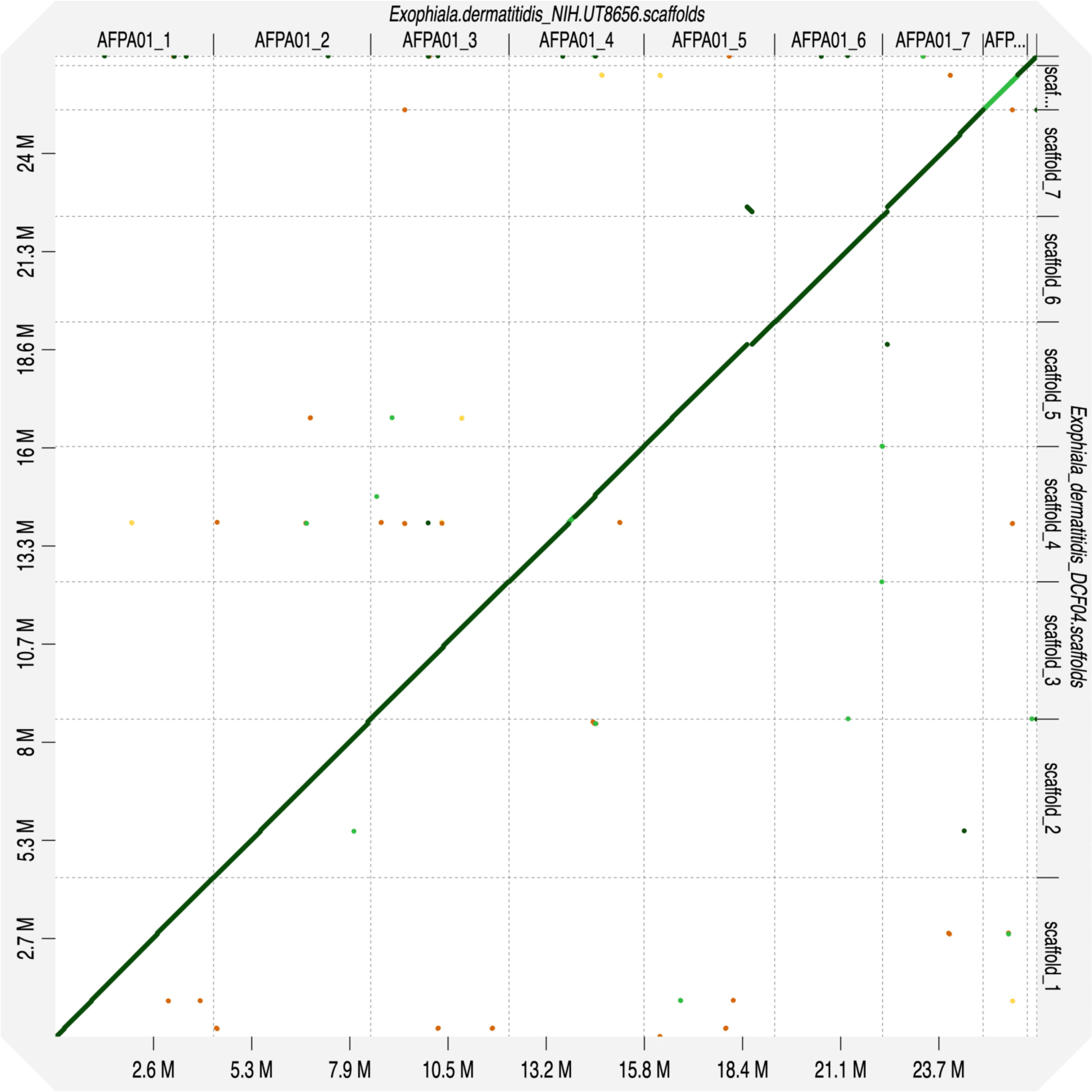

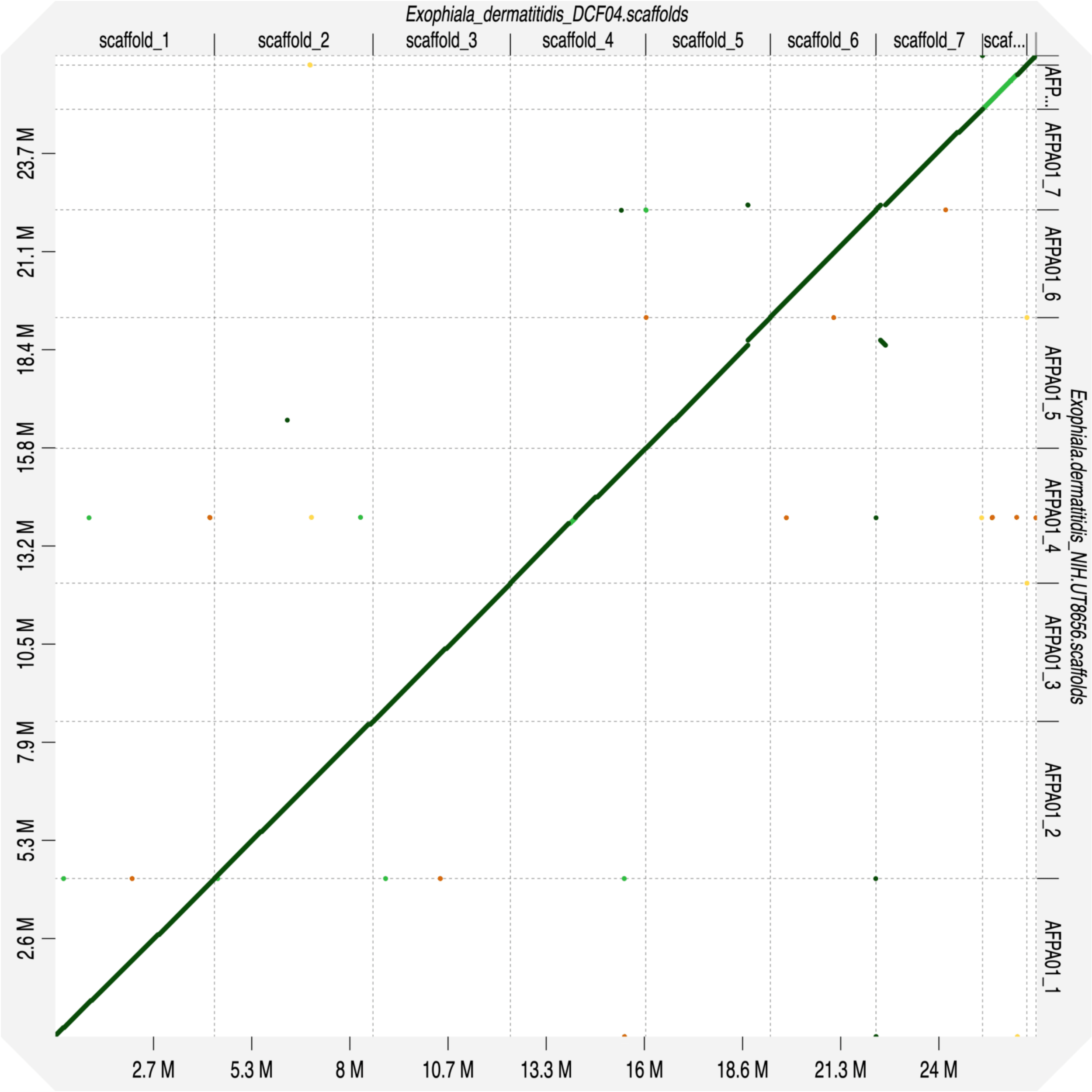

### Supplemental Figure 2. Mating-type determination of 23 clinical isolates of E. dermatitidis.

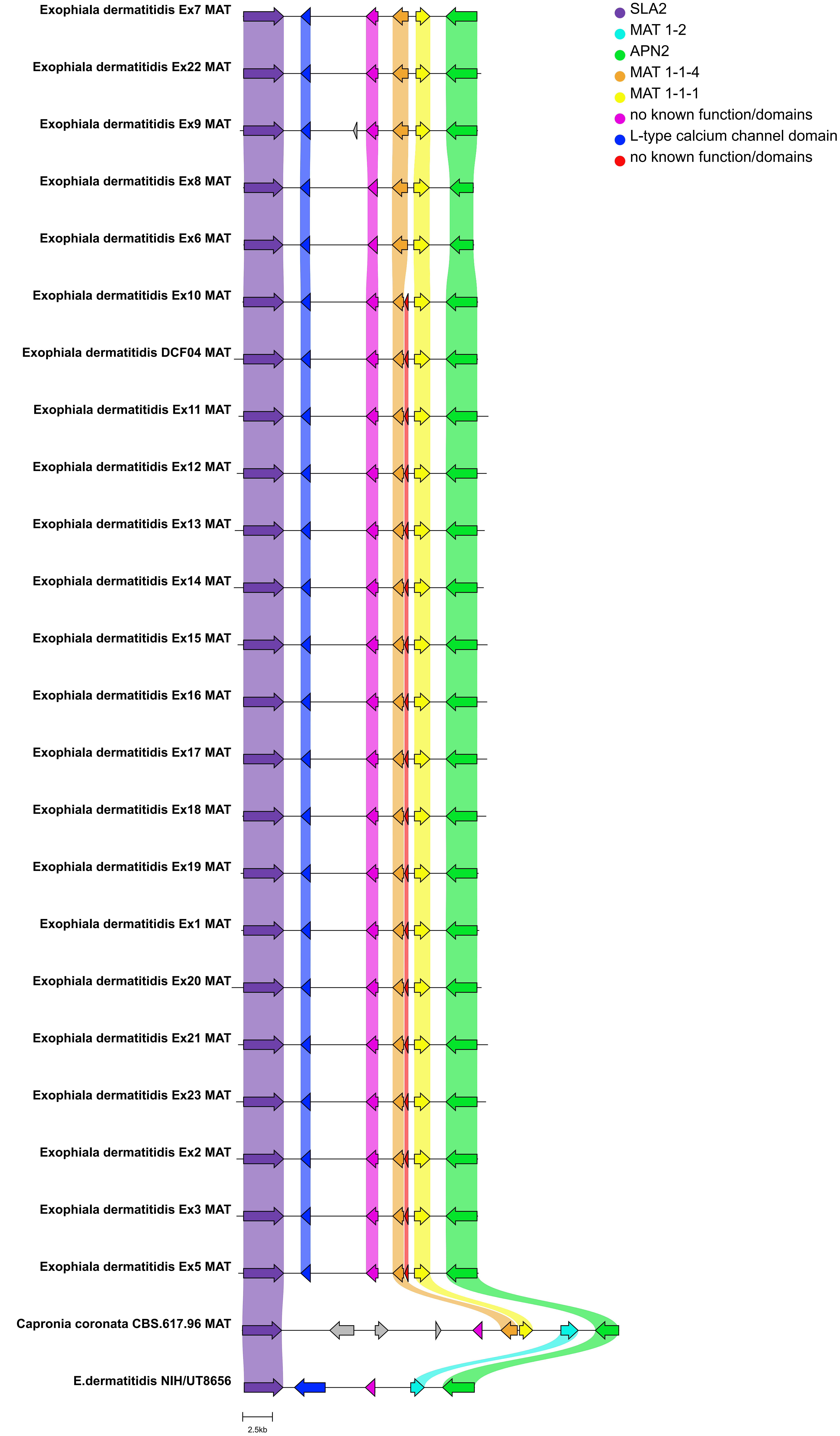

### Supplemental Figure 3. Phylogenetic tree of 24 isolates

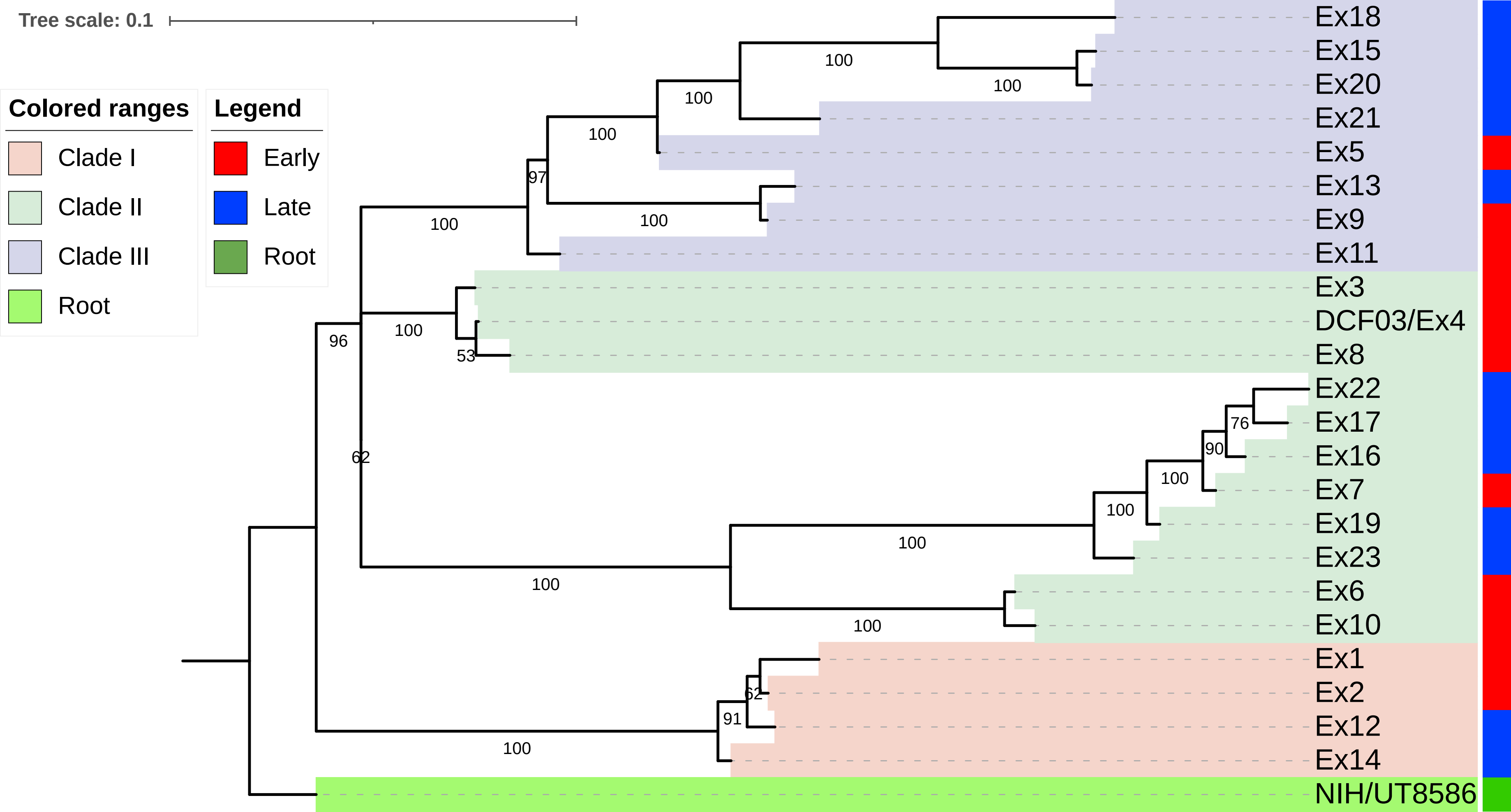
