## Supplemental Table 1. Collection and MIC values for CF patient derived E. dermatitidis isolates. for "In host evolution of *Exophiala dermatitidis* in cystic fibrosis lung micro-environment"

| Strain ID | Clade Node | MIC ng/ml | Date Isolated | Early or Late |
| --- | --- | --- | --- | --- |
| Ex2 | I | 500 | October 2, 2014 | Early |
| Ex1 | I | 250 | October 2, 2014 | Early |
| Ex12 | I | 250 | August 4, 2016 | Late |
| Ex14 | I | 62.5 | August 4, 2016 | Late |
| Ex6 | II | 250 | October 2, 2014 | Early |
| Ex10 | II | 62.5 | October 2, 2014 | Early |
| Ex22 | II | 62.5 | August 17, 2016 | Late |
| Ex17 | II | 250 | August 4, 2016 | Late |
| Ex16 | II | 500 | August 4, 2016 | Late |
| Ex7 | II | 500 | October 2, 2014 | Early |
| Ex19 | II | 250 | August 4, 2016 | Late |
| Ex23 | II | 250 | August 17, 2016 | Late |
| Ex3 | II | 125 | October 2, 2014 | Early |
| Ex8 | II | 125 | October 2, 2014 | Early |
| Ex4/DCF04 | II | 250 | October 2, 2014 | Early |
| Ex15 | III | 250 | August 4, 2016 | Late |
| Ex18 | III | 62.5 | August 4, 2016 | Late |
| Ex20 | III | 125 | August 17, 2016 | Late |
| Ex21 | III | 125 | August 17, 2016 | Late |
| Ex5 | III | 250 | October 2, 2014 | Early |
| Ex9 | III | 500 | October 2, 2014 | Early |
| Ex13 | III | 250 | August 4, 2016 | Late |
| Ex11 | III | 250 | October 2, 2014 | Early |
