## Supplemental Table 4. OrthoFinder summary comparing DCF04 and NIH/UT8656. for "In host evolution of *Exophiala dermatitidis* in cystic fibrosis lung micro-environment"

| **OrthoFinder Results** | ***E. dermatitidis* DCF04** | ***E. dermatitidis* NIH/UT8656** |
| --- | --- | --- |
| Number of genes | 9535 | 9285 |
| Number of genes in orthogroups | 8830 | 8810 |
| Number of unassigned genes (species-specific) | 705 | 475 |
| Number of genes in species-specific orthogroups | 34 | 24 |
| Total number of genes specific to each species | 739 | 499 |

| **Summary Statistics** |  |
| --- | --- |
| Number of genes | 18,820 |
| Number of genes in orthogroups | 17,640 |
| Number of unassigned genes | 1,180 |
| Number of orthogroups | 8,271 |
| Number of orthogroups with both taxa present | 8,256 |
| Number of single-copy orthogroups | 7,818 |

**Lineage-specific paralogs**

| **Orthogroup** | **DCF04** | **NIH/UT8656** | **Total** | **Function** |
| --- | --- | --- | --- | --- |
| OG0000035 | 0 | 8 | 8 | [ABC multidrug transporter](https://blast.ncbi.nlm.nih.gov/Blast.cgi#alnHdr_XP_009152512) [[Exophiala dermatitidis]](https://blast.ncbi.nlm.nih.gov/Blast.cgi#alnHdr_XP_009156165) |
| OG0000038 | 0 | 7 | 7 | [AAT family amino acid transporter [Exophiala dermatitidis NIH/UT8656]](https://blast.ncbi.nlm.nih.gov/Blast.cgi#alnHdr_XP_009154427) |
| OG0000108 | 0 | 5 | 5 | [5-oxoprolinase (ATP-hydrolysing) [Exophiala dermatitidis NIH/UT8656]](https://blast.ncbi.nlm.nih.gov/Blast.cgi#alnHdr_XP_009159207) |
| OG0008269 | 0 | 2 | 2 | [hypothetical protein HMPREF1120_05993 [Exophiala dermatitidis NIH/UT8656] DUF300-domain-containing protein](https://blast.ncbi.nlm.nih.gov/Blast.cgi#alnHdr_XP_009158434) |
| OG0008270 | 0 | 2 | 2 | [hypothetical protein HMPREF1120_08910 [Exophiala dermatitidis NIH/UT8656] P-loop containing nucleoside triphosphate hydrolase protein](https://blast.ncbi.nlm.nih.gov/Blast.cgi#alnHdr_XP_009161429) |
| OG0000020 | 8 | 0 | 8 | [ABC multidrug transporter [Exophiala dermatitidis]](https://blast.ncbi.nlm.nih.gov/Blast.cgi#alnHdr_XP_009156165) |
| OG0000043 | 6 | 0 | 6 | [AAT family amino acid transporter [Exophiala dermatitidis]](https://blast.ncbi.nlm.nih.gov/Blast.cgi#alnHdr_XP_009154601) |
| OG0000101 | 5 | 0 | 5 | [5-oxoprolinase (ATP-hydrolysing) [Exophiala dermatitidis]](https://blast.ncbi.nlm.nih.gov/Blast.cgi#alnHdr_XP_009156070) |
| OG0000020 | 8 | 0 | 8 | [ABC multidrug transporter [Exophiala dermatitidis]](https://blast.ncbi.nlm.nih.gov/Blast.cgi#alnHdr_XP_009156165) |
| OG0000043 | 6 | 0 | 6 | [AAT family amino acid transporter [Exophiala dermatitidis]](https://blast.ncbi.nlm.nih.gov/Blast.cgi#alnHdr_XP_009154601) |
| OG0000101 | 5 | 0 | 5 | [5-oxoprolinase (ATP-hydrolysing) [Exophiala dermatitidis]](https://blast.ncbi.nlm.nih.gov/Blast.cgi#alnHdr_XP_009156070) |
| OG0000377 | 3 | 0 | 3 | [hypothetical protein A1O5_09736 [Cladophialophora psammophila]](https://blast.ncbi.nlm.nih.gov/Blast.cgi#alnHdr_XP_007748505) |
| OG0001397 | 2 | 0 | 2 | [hypothetical protein KCU88_g6189 [Aureobasidium melanogenum]](https://blast.ncbi.nlm.nih.gov/Blast.cgi#alnHdr_KAG9772090) |

**Lineage-specific unassigned paralogs**

Full list can be found in Zenodo as OrthoFinder/Orthogroups_UnassignedGenes.tsv

doi: [10.5281/zenodo.7106110](https://doi.org/10.5281/zenodo.7106110)
