## Supplemental Table 5. Mating-type determination locus name descriptions of E. dermatitidis. for "In host evolution of *Exophiala dermatitidis* in cystic fibrosis lung micro-environment"

| **Species** | **SLA2 (purple)** | **L-domain (blue)** | **No known function/domain (pink)** | **MAT 1-2 (teal)** | **MAT 1-1-4 (orange)** | **MAT 1-1-1 (yellow)** | **APN2 (green)** |
| --- | --- | --- | --- | --- | --- | --- | --- |
| **NIH/UT8656** | HMPREF1120_08859 | HMPREF1120_08860 | HMPREF1120_08861 | HMPREF1120_08862 | x | x | HMPREF1120_08863 |
| **DCF04** | HRR96_009253 | HRR96_009234 | HRR96_009235 | x | HRR96_009236 | HRR96_009237 | HRR96_009238 |
| **CBS 617.98** | A1O1_07962 | x | A1O1_07968 | A1O1_07971 | A1O1_07969 | A1O1_07970 | A1O1_07972 |
