## Supplemental Table 7. All functional SNP and INDEL results for early and late pairs. for "In host evolution of *Exophiala dermatitidis* in cystic fibrosis lung micro-environment"

| *Pairwise Difference (Early vs Late)* | *Function* |
| --- | --- |
| *Ex1 & Ex12*  *15* | *hypothetical protein* [*HMPREF1120_00373*](https://www.ncbi.nlm.nih.gov/protein/EHY52156.1) *[Exophiala dermatitidis NIH/UT8656]*  *hypothetical protein* [*HMPREF1120_01565*](https://www.ncbi.nlm.nih.gov/protein/EHY53371.1) *[Exophiala dermatitidis NIH/UT8656]*  *MADS-box transcription factor* [*HMPREF1120_04857*](https://www.ncbi.nlm.nih.gov/protein/EHY56791.1) *[Exophiala dermatitidis NIH/UT8656]*  *G4 quadruplex nucleic acid binding protein [Exophiala dermatitidis NIH/UT8656]*  *hypothetical protein* [*HMPREF1120_02735*](https://www.ncbi.nlm.nih.gov/protein/EHY54567.1) *[Exophiala dermatitidis NIH/UT8656]*  *mitogen-activated protein kinase kinase kinase* [*HMPREF1120_04310*](https://www.ncbi.nlm.nih.gov/protein/EHY56220.1) *[Exophiala dermatitidis NIH/UT8656]*  *hypothetical protein* [*HMPREF1120_04204*](https://www.ncbi.nlm.nih.gov/protein/EHY56104.1) *[Exophiala dermatitidis NIH/UT8656]*  *hypothetical protein* [*HMPREF1120_04822*](https://www.ncbi.nlm.nih.gov/protein/EHY56755.1) *[Exophiala dermatitidis NIH/UT8656]*  *hypothetical protein* [*HMPREF1120_05199*](https://www.ncbi.nlm.nih.gov/protein/EHY57151.1) *[Exophiala dermatitidis NIH/UT8656]*  *hypothetical protein* [*HMPREF1120_05928*](https://www.ncbi.nlm.nih.gov/protein/EHY57907.1) *[Exophiala dermatitidis NIH/UT8656]*  *hypothetical protein* [*HMPREF1120_06924*](https://www.ncbi.nlm.nih.gov/protein/EHY58922.1) *[Exophiala dermatitidis NIH/UT8656]*  *AFG2 - ATPase of the CDC48/PAS1/SEC18 (AAA) family [Exophiala dermatitidis NIH/UT8656]*  *ALG2_2 - Alpha-1,3/1,6-mannosyltransferase [Exophiala dermatitidis NIH/UT8656]*  *DNA polymerase alpha subunit A* [*HMPREF1120_07994*](https://www.ncbi.nlm.nih.gov/protein/EHY60019.1) *[Exophiala dermatitidis NIH/UT8656]*  *GYP1 - Cis-golgi GTPase-activating protein [Exophiala dermatitidis NIH/UT8656]* |
| *Ex1 & Ex14*  *14* | *MADS-box transcription factor* [*HMPREF1120_04857*](https://www.ncbi.nlm.nih.gov/protein/EHY56791.1) *[Exophiala dermatitidis NIH/UT8656]*  *G4 quadruplex nucleic acid binding protein [Exophiala dermatitidis NIH/UT8656]*  *hypothetical protein* [*HMPREF1120_02735*](https://www.ncbi.nlm.nih.gov/protein/EHY54567.1) *[Exophiala dermatitidis NIH/UT8656]*  *hypothetical protein* [*HMPREF1120_04822*](https://www.ncbi.nlm.nih.gov/protein/EHY56755.1) *[Exophiala dermatitidis NIH/UT8656]*  *hypothetical protein* [*HMPREF1120_05199*](https://www.ncbi.nlm.nih.gov/protein/EHY57151.1) *[Exophiala dermatitidis NIH/UT8656]*  *monoamine oxidase* [*HMPREF1120_05597*](https://www.ncbi.nlm.nih.gov/protein/EHY57567.1) *[Exophiala dermatitidis NIH/UT8656]*  *biphenyl-2,3-diol 1,2-dioxygenase, variant* [*HMPREF1120_05880*](https://www.ncbi.nlm.nih.gov/protein/EHY57856.1) *[Exophiala dermatitidis NIH/UT8656]*  *hypothetical protein* [*HMPREF1120_05928*](https://www.ncbi.nlm.nih.gov/protein/EHY57907.1) *[Exophiala dermatitidis NIH/UT8656]*  *hypothetical protein* [*HMPREF1120_06924*](https://www.ncbi.nlm.nih.gov/protein/EHY58922.1) *[Exophiala dermatitidis NIH/UT8656]*  *AFG2 - ATPase of the CDC48/PAS1/SEC18 (AAA) family [Exophiala dermatitidis NIH/UT8656]*  *ALG2_2 - Alpha-1,3/1,6-mannosyltransferase [Exophiala dermatitidis NIH/UT8656]*  *DNA polymerase alpha subunit A* [*HMPREF1120_07994*](https://www.ncbi.nlm.nih.gov/protein/EHY60019.1) *[Exophiala dermatitidis NIH/UT8656]*  *hypothetical protein* [*HMPREF1120_08201*](https://www.ncbi.nlm.nih.gov/protein/EHY60233.1) *[Exophiala dermatitidis NIH/UT8656]*  *GYP1 - Cis-golgi GTPase-activating protein [Exophiala dermatitidis NIH/UT8656]* |
| *Ex2 & Ex12*  *12* | *hypothetical protein* [*HMPREF1120_00373*](https://www.ncbi.nlm.nih.gov/protein/EHY52156.1) *[Exophiala dermatitidis NIH/UT8656]*  *hypothetical protein* [*HMPREF1120_01565*](https://www.ncbi.nlm.nih.gov/protein/EHY53371.1) *[Exophiala dermatitidis NIH/UT8656]*  *MADS-box transcription factor* [*HMPREF1120_04857*](https://www.ncbi.nlm.nih.gov/protein/EHY56791.1) *[Exophiala dermatitidis NIH/UT8656]*  *G4 quadruplex nucleic acid binding protein [Exophiala dermatitidis NIH/UT8656]*  *mitogen-activated protein kinase kinase kinase* [*HMPREF1120_04310*](https://www.ncbi.nlm.nih.gov/protein/EHY56220.1) *[Exophiala dermatitidis NIH/UT8656]*  *MFS transporter, SP family, sugar:H+ symporter* [*HMPREF1120_06771*](https://www.ncbi.nlm.nih.gov/protein/EHY58768.1) *[Exophiala dermatitidis NIH/UT8656]*  *hypothetical protein* [*HMPREF1120_04204*](https://www.ncbi.nlm.nih.gov/protein/EHY56104.1) *[Exophiala dermatitidis NIH/UT8656]*  *hypothetical protein* [*HMPREF1120_04822*](https://www.ncbi.nlm.nih.gov/protein/EHY56755.1) *[Exophiala dermatitidis NIH/UT8656]*  *hypothetical protein* [*HMPREF1120_05199*](https://www.ncbi.nlm.nih.gov/protein/EHY57151.1) *[Exophiala dermatitidis NIH/UT8656]*  *BUD4 - Anillin-like protein involved in bud-site selection [Exophiala dermatitidis NIH/UT8656]*  *hypothetical protein* [*HMPREF1120_06469*](https://www.ncbi.nlm.nih.gov/protein/EHY58459.1) *[Exophiala dermatitidis NIH/UT8656]*  *hypothetical protein* [*HMPREF1120_08370*](https://www.ncbi.nlm.nih.gov/protein/EHY60408.1) *[Exophiala dermatitidis NIH/UT8656]* |
| *Ex2 & Ex14*  *9* | *MADS-box transcription factor* [*HMPREF1120_04857*](https://www.ncbi.nlm.nih.gov/protein/EHY56791.1) *[Exophiala dermatitidis NIH/UT8656]*  *G4 quadruplex nucleic acid binding protein[Exophiala dermatitidis NIH/UT8656]*  *MFS transporter, SP family, sugar:H+ symporter* [*HMPREF1120_06771*](https://www.ncbi.nlm.nih.gov/protein/EHY58768.1) *[Exophiala dermatitidis NIH/UT8656]*  *hypothetical protein* [*HMPREF1120_04822*](https://www.ncbi.nlm.nih.gov/protein/EHY56755.1) *[Exophiala dermatitidis NIH/UT8656]*  *monoamine oxidase [*[*HMPREF1120_05597*](https://www.ncbi.nlm.nih.gov/protein/EHY57567.1) *[Exophiala dermatitidis NIH/UT8656]*  *biphenyl-2,3-diol 1,2-dioxygenase, variant* [*HMPREF1120_05880*](https://www.ncbi.nlm.nih.gov/protein/EHY57856.1) *[Exophiala dermatitidis NIH/UT8656]*  *BUD4 - Anillin-like protein involved in bud-site selection [Exophiala dermatitidis NIH/UT8656]*  *hypothetical protein* [*HMPREF1120_08201*](https://www.ncbi.nlm.nih.gov/protein/EHY60233.1) *[Exophiala dermatitidis NIH/UT8656]*  *hypothetical protein* [*HMPREF1120_08370*](https://www.ncbi.nlm.nih.gov/protein/EHY60408.1) *[Exophiala dermatitidis NIH/UT8656]* |
| *Ex7 & Ex16*  *14* | *hypothetical protein* [*HMPREF1120_00271*](https://www.ncbi.nlm.nih.gov/protein/EHY52052.1) *[Exophiala dermatitidis NIH/UT8656]*  *hypothetical protein* [*HMPREF1120_00303*](https://www.ncbi.nlm.nih.gov/protein/EHY52084.1) *[Exophiala dermatitidis NIH/UT8656]*  *hypothetical protein* [*HMPREF1120_01139*](https://www.ncbi.nlm.nih.gov/protein/EHY52937.1) *[Exophiala dermatitidis NIH/UT8656]*  *hypothetical protein* [*HMPREF1120_02190*](https://www.ncbi.nlm.nih.gov/protein/EHY54013.1) *[Exophiala dermatitidis NIH/UT8656]*  *ADA HAT complex component 1* [*HMPREF1120_03635*](https://www.ncbi.nlm.nih.gov/protein/EHY55501.1) *[Exophiala dermatitidis NIH/UT8656]*  *MFS transporter, SP family, sugar:H+ symporter* [*HMPREF1120_06771*](https://www.ncbi.nlm.nih.gov/protein/EHY58768.1)  *[Exophiala dermatitidis NIH/UT8656]*  *NAD-dependent histone deacetylase SIR2* [*HMPREF1120_07852*](https://www.ncbi.nlm.nih.gov/protein/EHY59873.1)*[Exophiala dermatitidis NIH/UT8656]*  *cytochrome P450, family 7, subfamily B (oxysterol 7-alpha-hydroxylase)* [*HMPREF1120_04581*](https://www.ncbi.nlm.nih.gov/protein/XP_009156961.1) *[Exophiala dermatitidis NIH/UT8656]*  *hypothetical protein* [*HMPREF1120_04596*](https://www.ncbi.nlm.nih.gov/protein/EHY56515.1) *[Exophiala dermatitidis NIH/UT8656]*  *PAN3 - PABP1-Dependent Poly A-Specific Ribonuclease Subunit* [*HMPREF1120_05279*](https://www.ncbi.nlm.nih.gov/protein/EHY57233.1) *[Exophiala dermatitidis NIH/UT8656]*  *hypothetical protein* [*HMPREF1120_07418*](https://www.ncbi.nlm.nih.gov/protein/EHY59428.1) *[Exophiala dermatitidis NIH/UT8656]*  *queuine tRNA-ribosyltransferase* [*HMPREF1120_07977*](https://www.ncbi.nlm.nih.gov/protein/EHY60002.1)*[Exophiala dermatitidis NIH/UT8656]*  *hypothetical protein* [*HMPREF1120_08115*](https://www.ncbi.nlm.nih.gov/protein/EHY60143.1) *[Exophiala dermatitidis NIH/UT8656]*  *HemK protein* [*HMPREF1120_08430*](https://www.ncbi.nlm.nih.gov/protein/EHY60469.1)*[Exophiala dermatitidis NIH/UT8656]* |
| *Ex7 & Ex17*  *7* | *hypothetical protein* [*HMPREF1120_00271*](https://www.ncbi.nlm.nih.gov/protein/EHY52052.1) *[Exophiala dermatitidis NIH/UT8656]*  *hypothetical protein* [*HMPREF1120_01760*](https://www.ncbi.nlm.nih.gov/protein/EHY53571.1) *[Exophiala dermatitidis NIH/UT8656]*  *Strongly-conserved Zn-finger binding protein (TFIIIA) [Exophiala dermatitidis NIH/UT8656]*  *hypothetical protein* [*HMPREF1120_02190*](https://www.ncbi.nlm.nih.gov/protein/EHY54013.1) *[Exophiala dermatitidis NIH/UT8656]*  *ADA HAT complex component 1* [*HMPREF1120_03635*](https://www.ncbi.nlm.nih.gov/protein/EHY55501.1) *[Exophiala dermatitidis NIH/UT8656]*  *PAN3 - PABP1-Dependent Poly A-Specific Ribonuclease Subunit* [*HMPREF1120_05279*](https://www.ncbi.nlm.nih.gov/protein/EHY57233.1) *[Exophiala dermatitidis NIH/UT8656]*  *hypothetical protein* [*HMPREF1120_06447*](https://www.ncbi.nlm.nih.gov/protein/EHY58437.1) *[Exophiala dermatitidis NIH/UT8656]* |
| *Ex7 & Ex19*  *14* | *hypothetical protein* [*HMPREF1120_00271*](https://www.ncbi.nlm.nih.gov/protein/EHY52052.1) *[Exophiala dermatitidis NIH/UT8656]*  *hypothetical protein* [*HMPREF1120_01139*](https://www.ncbi.nlm.nih.gov/protein/EHY52937.1) *[Exophiala dermatitidis NIH/UT8656]*  *alpha-1,2-mannosyltransferase* [*HMPREF1120_07833*](https://www.ncbi.nlm.nih.gov/protein/EHY59853.1)*[Exophiala dermatitidis NIH/UT8656]*  *ribosome biosynthesis protein rrb1[Exophiala dermatitidis NIH/UT8656]*  *hypothetical protein* [*HMPREF1120_05142*](https://www.ncbi.nlm.nih.gov/protein/EHY57092.1) *[Exophiala dermatitidis NIH/UT8656]*  *ADA HAT complex component 1* [*HMPREF1120_03635*](https://www.ncbi.nlm.nih.gov/protein/EHY55501.1) *[Exophiala dermatitidis NIH/UT8656]*  *MFS transporter, SP family, sugar:H+ symporter* [*HMPREF1120_06771*](https://www.ncbi.nlm.nih.gov/protein/EHY58768.1)  *[Exophiala dermatitidis NIH/UT8656]*  *hypothetical protein* [*HMPREF1120_04204*](https://www.ncbi.nlm.nih.gov/protein/EHY56104.1) *[Exophiala dermatitidis NIH/UT8656]*  *DUS3 - Dihydrouridine synthase* [*HMPREF1120_04489*](https://www.ncbi.nlm.nih.gov/protein/EHY56407.1) *[Exophiala dermatitidis NIH/UT8656]*  *hypothetical protein* [*HMPREF1120_04676*](https://www.ncbi.nlm.nih.gov/protein/EHY56600.1) *[Exophiala dermatitidis NIH/UT8656]*  *PAN3 - PABP1-Dependent Poly A-Specific Ribonuclease Subunit* [*HMPREF1120_05279*](https://www.ncbi.nlm.nih.gov/protein/EHY57233.1) *[Exophiala dermatitidis NIH/UT8656]*  *monoamine oxidase [*[*HMPREF1120_05597*](https://www.ncbi.nlm.nih.gov/protein/EHY57567.1) *[Exophiala dermatitidis NIH/UT8656]*  *hypothetical protein* [*HMPREF1120_06612*](https://www.ncbi.nlm.nih.gov/protein/EHY58604.1) *[Exophiala dermatitidis NIH/UT8656]*  *hypothetical protein* [*HMPREF1120_08252*](https://www.ncbi.nlm.nih.gov/protein/EHY60284.1) *[Exophiala dermatitidis NIH/UT8656]* |
| *Ex7 & Ex22*  *6* | *hypothetical protein* [*HMPREF1120_00271*](https://www.ncbi.nlm.nih.gov/protein/EHY52052.1) *[Exophiala dermatitidis NIH/UT8656]*  *hypothetical protein* [*HMPREF1120_01139*](https://www.ncbi.nlm.nih.gov/protein/EHY52937.1) *[Exophiala dermatitidis NIH/UT8656]*  *hypothetical protein* [*HMPREF1120_02190*](https://www.ncbi.nlm.nih.gov/protein/EHY54013.1) *[Exophiala dermatitidis NIH/UT8656]*  *ADA HAT complex component 1* [*HMPREF1120_03635*](https://www.ncbi.nlm.nih.gov/protein/EHY55501.1) *[Exophiala dermatitidis NIH/UT8656]*  *Ring finger and CHY zinc finger domain-containing protein 1* [*HMPREF1120_04104*](https://www.ncbi.nlm.nih.gov/protein/EHY55998.1) *[Exophiala dermatitidis NIH/UT8656]*  *PAN3 - PABP1-Dependent Poly A-Specific Ribonuclease Subunit* [*HMPREF1120_05279*](https://www.ncbi.nlm.nih.gov/protein/EHY57233.1) *[Exophiala dermatitidis NIH/UT8656]* |
| *Ex7 & Ex23*  *22* | *hypothetical protein* [*HMPREF1120_00215*](https://www.ncbi.nlm.nih.gov/protein/EHY51992.1) *[Exophiala dermatitidis NIH/UT8656]*  *hypothetical protein* [*HMPREF1120_00271*](https://www.ncbi.nlm.nih.gov/protein/EHY52052.1) *[Exophiala dermatitidis NIH/UT8656]*  *hypothetical protein* [*HMPREF1120_01244*](https://www.ncbi.nlm.nih.gov/protein/EHY53043.1) *[Exophiala dermatitidis NIH/UT8656]*  *hypothetical protein* [*HMPREF1120_01346*](https://www.ncbi.nlm.nih.gov/protein/EHY53148.1) *[Exophiala dermatitidis NIH/UT8656]*  *MFS transporter, DHA1 family, multidrug resistance protein* [*HMPREF1120_09017*](https://www.ncbi.nlm.nih.gov/protein/EHY61079.1) *[Exophiala dermatitidis NIH/UT8656]*  *transcriptional regulatory protein GAL4* [*HMPREF1120_08700*](https://www.ncbi.nlm.nih.gov/protein/EHY60755.1)*[Exophiala dermatitidis NIH/UT8656]*  *G4 quadruplex nucleic acid binding protein [Exophiala dermatitidis NIH/UT8656]*  *ribosome biosynthesis protein rrb1[Exophiala dermatitidis NIH/UT8656]*  *hypothetical protein* [*HMPREF1120_02823*](https://www.ncbi.nlm.nih.gov/protein/EHY54656.1) *[Exophiala dermatitidis NIH/UT8656]*  *hypothetical protein* [*HMPREF1120_05142*](https://www.ncbi.nlm.nih.gov/protein/EHY57092.1) *[Exophiala dermatitidis NIH/UT8656]*  *2,3-bisphosphoglycerate-dependent phosphoglycerate mutase* [*HMPREF1120_02960*](https://www.ncbi.nlm.nih.gov/protein/EHY54796.1) *[Exophiala dermatitidis NIH/UT8656]*  *phosphatidylinositol-bisphosphatase* [*HMPREF1120_04965*](https://www.ncbi.nlm.nih.gov/protein/EHY56901.1)*[Exophiala dermatitidis NIH/UT8656]*  *ADA HAT complex component 1* [*HMPREF1120_03635*](https://www.ncbi.nlm.nih.gov/protein/EHY55501.1) *[Exophiala dermatitidis NIH/UT8656]*  *hypothetical protein* [*HMPREF1120_04204*](https://www.ncbi.nlm.nih.gov/protein/EHY56104.1) *[Exophiala dermatitidis NIH/UT8656]*  *hypothetical protein* [*HMPREF1120_04403*](https://www.ncbi.nlm.nih.gov/protein/EHY56319.1) *[Exophiala dermatitidis NIH/UT8656]*  *PAN3 - PABP1-Dependent Poly A-Specific Ribonuclease Subunit* [*HMPREF1120_05279*](https://www.ncbi.nlm.nih.gov/protein/EHY57233.1) *[Exophiala dermatitidis NIH/UT8656]*  *nucleolin* [*HMPREF1120_07826*](https://www.ncbi.nlm.nih.gov/protein/EHY59846.1)*[Exophiala dermatitidis NIH/UT8656]*  *hypothetical protein* [*HMPREF1120_05877*](https://www.ncbi.nlm.nih.gov/protein/EHY57853.1) *[Exophiala dermatitidis NIH/UT8656]*  *hypothetical protein* [*HMPREF1120_05976*](https://www.ncbi.nlm.nih.gov/protein/EHY57956.1) *[Exophiala dermatitidis NIH/UT8656]*  *inositol oxygenase* [*HMPREF1120_07129*](https://www.ncbi.nlm.nih.gov/protein/EHY59131.1)*[Exophiala dermatitidis NIH/UT8656]*  *hypothetical protein* [*HMPREF1120_09068*](https://www.ncbi.nlm.nih.gov/protein/EHY61132.1) *[Exophiala dermatitidis NIH/UT8656]*  *hypothetical protein* [*HMPREF1120_09075*](https://www.ncbi.nlm.nih.gov/protein/EHY61139.1) *[Exophiala dermatitidis NIH/UT8656]* |
| *Ex5 & Ex15*  *120* | *DNA repair protein RAD50 [Exophiala dermatitidis NIH/UT8656]*  *hypothetical protein* [*HMPREF1120_00457*](https://www.ncbi.nlm.nih.gov/protein/EHY52242.1) *[Exophiala dermatitidis NIH/UT8656]*  *hypothetical protein* [*HMPREF1120_00646*](https://www.ncbi.nlm.nih.gov/protein/EHY52434.1) *[Exophiala dermatitidis NIH/UT8656]*  *eukaryotic translation initiation factor 2 subunit gamma* [*HMPREF1120_00677*](https://www.ncbi.nlm.nih.gov/protein/EHY52465.1)  *[Exophiala dermatitidis NIH/UT8656]*  *cytochrome P450 oxidoreductase* [*HMPREF1120_01361*](https://www.ncbi.nlm.nih.gov/protein/EHY53163.1) *[Exophiala dermatitidis NIH/UT8656]*  *hypothetical protein* [*HMPREF1120_00830*](https://www.ncbi.nlm.nih.gov/protein/EHY52619.1) *[Exophiala dermatitidis NIH/UT8656]*  *hypothetical protein* [*HMPREF1120_01089*](https://www.ncbi.nlm.nih.gov/protein/EHY52883.1) *[Exophiala dermatitidis NIH/UT8656]*  *hypothetical protein* [*HMPREF1120_01151*](https://www.ncbi.nlm.nih.gov/protein/EHY52950.1) *[Exophiala dermatitidis NIH/UT8656]*  *sulfite oxidase* [*HMPREF1120_05227*](https://www.ncbi.nlm.nih.gov/protein/EHY57179.1) *[Exophiala dermatitidis NIH/UT8656]*  *prolyl-tRNA synthetase* [*HMPREF1120_01354*](https://www.ncbi.nlm.nih.gov/protein/EHY53156.1) *[Exophiala dermatitidis NIH/UT8656]*  *cytochrome P450 oxidoreductase* [*HMPREF1120_01361*](https://www.ncbi.nlm.nih.gov/protein/EHY53163.1) *[Exophiala dermatitidis NIH/UT8656]*  *hypothetical protein* [*HMPREF1120_01471*](https://www.ncbi.nlm.nih.gov/protein/EHY53277.1) *[Exophiala dermatitidis NIH/UT8656]*  *serine/threonine kinase 16* [*HMPREF1120_06920*](https://www.ncbi.nlm.nih.gov/protein/EHY58918.1) *[Exophiala dermatitidis NIH/UT8656]*  *hypothetical protein* [*HMPREF1120_02055*](https://www.ncbi.nlm.nih.gov/protein/EHY53875.1) *[Exophiala dermatitidis NIH/UT8656]*  *hypothetical protein* [*HMPREF1120_02083*](https://www.ncbi.nlm.nih.gov/protein/EHY53903.1) *[Exophiala dermatitidis NIH/UT8656]*  *G4 quadruplex nucleic acid binding protein [Exophiala dermatitidis NIH/UT8656]*  *hypothetical protein* [*HMPREF1120_02190*](https://www.ncbi.nlm.nih.gov/protein/EHY54013.1) *[Exophiala dermatitidis NIH/UT8656]*  *glutathione S-transferase* [*HMPREF1120_08143*](https://www.ncbi.nlm.nih.gov/protein/EHY60173.1) *[Exophiala dermatitidis NIH/UT8656]*  *nicotinate-nucleotide diphosphorylase (carboxylating)* [*HMPREF1120_02317*](https://www.ncbi.nlm.nih.gov/protein/EHY54142.1)  *[Exophiala dermatitidis NIH/UT8656]*  *phosphodiesterase/alkaline phosphatase D* [*HMPREF1120_02364*](https://www.ncbi.nlm.nih.gov/protein/EHY54190.1) *[Exophiala dermatitidis NIH/UT8656]*  *Xanthine phosphoribosyltransferase 1* [*HMPREF1120_06110*](https://www.ncbi.nlm.nih.gov/protein/EHY58092.1) *[Exophiala dermatitidis NIH/UT8656]*  *Xanthine phosphoribosyltransferase 1* [*HMPREF1120_06110*](https://www.ncbi.nlm.nih.gov/protein/EHY58092.1) *[Exophiala dermatitidis NIH/UT8656]*  *rRNA (cytosine-C5-)-methyltransferase nop2* [*HMPREF1120_02613*](https://www.ncbi.nlm.nih.gov/protein/EHY54444.1) *[Exophiala dermatitidis NIH/UT8656]*  *transformation/transcription domain-associated protein* [*HMPREF1120_02639*](https://www.ncbi.nlm.nih.gov/protein/EHY54471.1)  *[Exophiala dermatitidis NIH/UT8656]*  *Serine/threonine-protein phosphatase 2A 56 kDa regulatory subunit delta isoform* [*HMPREF1120_01344*](https://www.ncbi.nlm.nih.gov/protein/EHY53146.1) *[Exophiala dermatitidis NIH/UT8656]*  *hypothetical protein* [*HMPREF1120_02700*](https://www.ncbi.nlm.nih.gov/protein/EHY54532.1) *[Exophiala dermatitidis NIH/UT8656]*  *hypothetical protein* [*HMPREF1120_02708*](https://www.ncbi.nlm.nih.gov/protein/EHY54540.1) *[Exophiala dermatitidis NIH/UT8656]*  *hypothetical protein* [*HMPREF1120_02758*](https://www.ncbi.nlm.nih.gov/protein/EHY54590.1) *[Exophiala dermatitidis NIH/UT8656]*  *hypothetical protein* [*HMPREF1120_02784*](https://www.ncbi.nlm.nih.gov/protein/EHY54616.1) *[Exophiala dermatitidis NIH/UT8656]*  *MOT1 - TATA-binding protein-associated factor* [*HMPREF1120_06808*](https://www.ncbi.nlm.nih.gov/protein/EHY58805.1)  *[Exophiala dermatitidis NIH/UT8656]*  *ING3 - Inhibitor of growth protein 3* [*HMPREF1120_05286*](https://www.ncbi.nlm.nih.gov/protein/EHY57240.1) *[Exophiala dermatitidis NIH/UT8656]*  *hypothetical protein* [*HMPREF1120_02908*](https://www.ncbi.nlm.nih.gov/protein/EHY54743.1) *[Exophiala dermatitidis NIH/UT8656]*  *PDE2 - High-affinity cyclic AMP phosphodiesterase [Exophiala dermatitidis NIH/UT8656]*  *hypothetical protein* [*HMPREF1120_03003*](https://www.ncbi.nlm.nih.gov/protein/EHY54840.1) *[Exophiala dermatitidis NIH/UT8656]*  *hypothetical protein* [*HMPREF1120_03007*](https://www.ncbi.nlm.nih.gov/protein/EHY54845.1) *[Exophiala dermatitidis NIH/UT8656]*  *VE1 - Verticillium wilt disease resistance protein [Exophiala dermatitidis NIH/UT8656]*  *hypothetical protein* [*HMPREF1120_03359*](https://www.ncbi.nlm.nih.gov/protein/EHY55214.1) *[Exophiala dermatitidis NIH/UT8656]*  *hypothetical protein* [*HMPREF1120_03367*](https://www.ncbi.nlm.nih.gov/protein/EHY55222.1) *[Exophiala dermatitidis NIH/UT8656]*  *hypothetical protein* [*HMPREF1120_03513*](https://www.ncbi.nlm.nih.gov/protein/EHY55374.1) *[Exophiala dermatitidis NIH/UT8656]*  *gibberellin 2-oxidase* [*HMPREF1120_09208*](https://www.ncbi.nlm.nih.gov/protein/EHY61274.1)*[Exophiala dermatitidis NIH/UT8656]*  *hypothetical protein* [*HMPREF1120_03910*](https://www.ncbi.nlm.nih.gov/protein/EHY55786.1) *[Exophiala dermatitidis NIH/UT8656]*  *hypothetical protein* [*HMPREF1120_03911*](https://www.ncbi.nlm.nih.gov/protein/EHY55787.1) *[Exophiala dermatitidis NIH/UT8656]*  *hypothetical protein* [*HMPREF1120_03928*](https://www.ncbi.nlm.nih.gov/protein/EHY55806.1) *[Exophiala dermatitidis NIH/UT8656]*  *amidohydrolase* [*HMPREF1120_03964*](https://www.ncbi.nlm.nih.gov/protein/EHY55847.1)*[Exophiala dermatitidis NIH/UT8656]*  *hypothetical protein* [*HMPREF1120_04101*](https://www.ncbi.nlm.nih.gov/protein/EHY55995.1) *[Exophiala dermatitidis NIH/UT8656]*  *MFS transporter, SP family, sugar:H+ symporter* [*HMPREF1120_06771*](https://www.ncbi.nlm.nih.gov/protein/EHY58768.1)  *[Exophiala dermatitidis NIH/UT8656]*  *COP9 signalosome complex subunit 2* [*HMPREF1120_04182*](https://www.ncbi.nlm.nih.gov/protein/EHY56082.1)*[Exophiala dermatitidis NIH/UT8656]*  *hypothetical protein* [*HMPREF1120_04204*](https://www.ncbi.nlm.nih.gov/protein/EHY56104.1) *[Exophiala dermatitidis NIH/UT8656]*  *SIP3 - Putative sterol transfer protein* [*HMPREF1120_04207*](https://www.ncbi.nlm.nih.gov/protein/EHY56107.1)*[Exophiala dermatitidis NIH/UT8656]*  *hypothetical protein* [*HMPREF1120_04314*](https://www.ncbi.nlm.nih.gov/protein/EHY56224.1) *[Exophiala dermatitidis NIH/UT8656]*  *hypothetical protein* [*HMPREF1120_04349*](https://www.ncbi.nlm.nih.gov/protein/EHY56262.1) *[Exophiala dermatitidis NIH/UT8656]*  *hypothetical protein* [*HMPREF1120_04459*](https://www.ncbi.nlm.nih.gov/protein/EHY56377.1) *[Exophiala dermatitidis NIH/UT8656]*  *tyrosinase* [*HMPREF1120_04514*](https://www.ncbi.nlm.nih.gov/protein/EHY56432.1)*[Exophiala dermatitidis NIH/UT8656]*  *hypothetical protein* [*HMPREF1120_04584*](https://www.ncbi.nlm.nih.gov/protein/EHY56503.1) *[Exophiala dermatitidis NIH/UT8656]*  *ankyrin* [*HMPREF1120_08463*](https://www.ncbi.nlm.nih.gov/protein/EHY60507.1)*[Exophiala dermatitidis NIH/UT8656]*  *hypothetical protein* [*HMPREF1120_04645*](https://www.ncbi.nlm.nih.gov/protein/EHY56567.1) *[Exophiala dermatitidis NIH/UT8656]*  *adenosinetriphosphatase* [*HMPREF1120_09246*](https://www.ncbi.nlm.nih.gov/protein/EHY61312.1) *[Exophiala dermatitidis NIH/UT8656]*  *hypothetical protein* [*HMPREF1120_04673*](https://www.ncbi.nlm.nih.gov/protein/EHY56597.1) *[Exophiala dermatitidis NIH/UT8656]*  *alkanesulfonate monooxygenase* [*HMPREF1120_08264*](https://www.ncbi.nlm.nih.gov/protein/EHY60297.1)*[Exophiala dermatitidis NIH/UT8656]*  *hypothetical protein* [*HMPREF1120_04695*](https://www.ncbi.nlm.nih.gov/protein/EHY56619.1) *[Exophiala dermatitidis NIH/UT8656]*  *hypothetical protein* [*HMPREF1120_04835*](https://www.ncbi.nlm.nih.gov/protein/EHY56769.1) *[Exophiala dermatitidis NIH/UT8656]*  *hypothetical protein* [*HMPREF1120_04991*](https://www.ncbi.nlm.nih.gov/protein/EHY56927.1) *[Exophiala dermatitidis NIH/UT8656]*  *4-coumarate-CoA ligase* [*HMPREF1120_09210*](https://www.ncbi.nlm.nih.gov/protein/EHY61276.1)*[Exophiala dermatitidis NIH/UT8656]*  *hypothetical protein* [*HMPREF1120_05151*](https://www.ncbi.nlm.nih.gov/protein/EHY57101.1) *[Exophiala dermatitidis NIH/UT8656]*  *hypothetical protein* [*HMPREF1120_05400*](https://www.ncbi.nlm.nih.gov/protein/EHY57359.1) *[Exophiala dermatitidis NIH/UT8656]*  *dihydrodipicolinate synthetase* [*HMPREF1120_09161*](https://www.ncbi.nlm.nih.gov/protein/EHY61225.1)*[Exophiala dermatitidis NIH/UT8656]*  *hypothetical protein* [*HMPREF1120_05622*](https://www.ncbi.nlm.nih.gov/protein/EHY57593.1) *[Exophiala dermatitidis NIH/UT8656]*  *hypothetical protein* [*HMPREF1120_05658*](https://www.ncbi.nlm.nih.gov/protein/EHY57629.1) *[Exophiala dermatitidis NIH/UT8656]*  *PSY2 - Platinum sensitivity protein [Exophiala dermatitidis NIH/UT8656]*  *hypothetical protein* [*HMPREF1120_05728*](https://www.ncbi.nlm.nih.gov/protein/EHY57701.1) *[Exophiala dermatitidis NIH/UT8656]*  *SPT20 - Transcription factor spt20* [*HMPREF1120_05758*](https://www.ncbi.nlm.nih.gov/protein/EHY57731.1)*[Exophiala dermatitidis NIH/UT8656]*  *hypothetical protein* [*HMPREF1120_05809*](https://www.ncbi.nlm.nih.gov/protein/EHY57785.1) *[Exophiala dermatitidis NIH/UT8656]*  *ETF1 - elongation factor 2* [*HMPREF1120_05986*](https://www.ncbi.nlm.nih.gov/protein/EHY57966.1)*[Exophiala dermatitidis NIH/UT8656]*  *hypothetical protein* [*HMPREF1120_06093*](https://www.ncbi.nlm.nih.gov/protein/EHY58075.1) *[Exophiala dermatitidis NIH/UT8656]*  *RGT1 - Glucose-responsive transcription factor 1 [Exophiala dermatitidis NIH/UT8656]*  *hypothetical protein* [*HMPREF1120_06177*](https://www.ncbi.nlm.nih.gov/protein/EHY58163.1) *[Exophiala dermatitidis NIH/UT8656]*  *hypothetical protein* [*HMPREF1120_06361*](https://www.ncbi.nlm.nih.gov/protein/EHY58349.1) *[Exophiala dermatitidis NIH/UT8656]*  *37S ribosomal protein, mitochondrial [Exophiala dermatitidis NIH/UT8656]*  *hypothetical protein* [*HMPREF1120_06500*](https://www.ncbi.nlm.nih.gov/protein/EHY58490.1) *[Exophiala dermatitidis NIH/UT8656]*  *hypothetical protein* [*HMPREF1120_06642*](https://www.ncbi.nlm.nih.gov/protein/EHY58637.1) *[Exophiala dermatitidis NIH/UT8656]*  *hypothetical protein* [*HMPREF1120_06878*](https://www.ncbi.nlm.nih.gov/protein/EHY58876.1) *[Exophiala dermatitidis NIH/UT8656]*  *hypothetical protein* [*HMPREF1120_06971*](https://www.ncbi.nlm.nih.gov/protein/EHY58970.1) *[Exophiala dermatitidis NIH/UT8656]*  *transcription initiation factor TFIID subunit D2* [*MPREF1120_06984*](https://www.ncbi.nlm.nih.gov/protein/EHY58983.1)*[Exophiala dermatitidis NIH/UT8656]*  *L-galactose dehydrogenase* [*HMPREF1120_07000*](https://www.ncbi.nlm.nih.gov/protein/EHY59000.1)*[Exophiala dermatitidis NIH/UT8656]*  *hypothetical protein* [*HMPREF1120_07085*](https://www.ncbi.nlm.nih.gov/protein/EHY59086.1) *[Exophiala dermatitidis NIH/UT8656]*  *3' exoribonuclease* [*HMPREF1120_08304*](https://www.ncbi.nlm.nih.gov/protein/EHY60338.1)*[Exophiala dermatitidis NIH/UT8656]*  *hypothetical protein* [*HMPREF1120_07255*](https://www.ncbi.nlm.nih.gov/protein/EHY59262.1) *[Exophiala dermatitidis NIH/UT8656]*  *hypothetical protein* [*HMPREF1120_07291*](https://www.ncbi.nlm.nih.gov/protein/EHY59299.1) *[Exophiala dermatitidis NIH/UT8656]*  *hypothetical protein* [*HMPREF1120_07306*](https://www.ncbi.nlm.nih.gov/protein/EHY59314.1) *[Exophiala dermatitidis NIH/UT8656]*  *hypothetical protein* [*HMPREF1120_07433*](https://www.ncbi.nlm.nih.gov/protein/EHY59443.1) *[Exophiala dermatitidis NIH/UT8656]*  *RAS2 - Ras GTPase* [*HMPREF1120_01421*](https://www.ncbi.nlm.nih.gov/protein/EHY53224.1)*[Exophiala dermatitidis NIH/UT8656]*  *hypothetical protein* [*HMPREF1120_07616*](https://www.ncbi.nlm.nih.gov/protein/EHY59631.1) *[Exophiala dermatitidis NIH/UT8656]*  *D-3-phosphoglycerate dehydrogenase* [*HMPREF1120_06805*](https://www.ncbi.nlm.nih.gov/protein/EHY58802.1)*[Exophiala dermatitidis NIH/UT8656]*  *ankyrin* [*HMPREF1120_08463*](https://www.ncbi.nlm.nih.gov/protein/EHY60507.1)*[Exophiala dermatitidis NIH/UT8656]*  *nuclear transcription factor Y, alpha* [*HMPREF1120_07714*](https://www.ncbi.nlm.nih.gov/protein/EHY59731.1)*[Exophiala dermatitidis NIH/UT8656]*  *CHS3 - chitin synthase class 3* [*HMPREF1120_08776*](https://www.ncbi.nlm.nih.gov/protein/EHY60832.1) *[Exophiala dermatitidis NIH/UT8656]*  *salicylate hydroxylase* [*HMPREF1120_03459*](https://www.ncbi.nlm.nih.gov/protein/EHY55317.1)*[Exophiala dermatitidis NIH/UT8656]*  *hypothetical protein* [*HMPREF1120_06589*](https://www.ncbi.nlm.nih.gov/protein/EHY58580.1) *[Exophiala dermatitidis NIH/UT8656]*  *hypothetical protein* [*HMPREF1120_06584*](https://www.ncbi.nlm.nih.gov/protein/EHY58575.1) *[Exophiala dermatitidis NIH/UT8656]*  *queuine tRNA-ribosyltransferase* [*HMPREF1120_07977*](https://www.ncbi.nlm.nih.gov/protein/EHY60002.1)*[Exophiala dermatitidis NIH/UT8656]*  *hypothetical protein* [*HMPREF1120_07985*](https://www.ncbi.nlm.nih.gov/protein/EHY60010.1) *[Exophiala dermatitidis NIH/UT8656]*  *thiamin biosynthesis protein* [*HMPREF1120_07987*](https://www.ncbi.nlm.nih.gov/protein/EHY60012.1)*[Exophiala dermatitidis NIH/UT8656]*  *ABD1 - mRNA cap guanine-N7 methyltransferase* [*HMPREF1120_06541*](https://www.ncbi.nlm.nih.gov/protein/EHY58531.1)*[Exophiala dermatitidis NIH/UT8656]*  *PAN2 - poly(A) specific ribonuclease [Exophiala dermatitidis NIH/UT8656]*  *DOA4 - ubiquitin specific protease* [*HMPREF1120_06573*](https://www.ncbi.nlm.nih.gov/protein/EHY58564.1)*[Exophiala dermatitidis NIH/UT8656]*  *amidase* [*HMPREF1120_09153*](https://www.ncbi.nlm.nih.gov/protein/EHY61217.1)*[Exophiala dermatitidis NIH/UT8656]*  *MEF2 - Ribosome-releasing factor 2, mitochondrial[Exophiala dermatitidis NIH/UT8656]*  *RGA2 - Rho-type gtpase-activating protein[Exophiala dermatitidis NIH/UT8656]*  *hypothetical protein* [*HMPREF1120_08236*](https://www.ncbi.nlm.nih.gov/protein/EHY60268.1) *[Exophiala dermatitidis NIH/UT8656]*  *cytochrome P450 oxidoreductase* [*HMPREF1120_01361*](https://www.ncbi.nlm.nih.gov/protein/EHY53163.1) *[Exophiala dermatitidis NIH/UT8656]*  *hypothetical protein* [*HMPREF1120_08425*](https://www.ncbi.nlm.nih.gov/protein/EHY60464.1) *[Exophiala dermatitidis NIH/UT8656]*  *hydrolase* [*HMPREF1120_08460*](https://www.ncbi.nlm.nih.gov/protein/EHY60504.1)*[Exophiala dermatitidis NIH/UT8656]*  *hypothetical protein* [*HMPREF1120_08629*](https://www.ncbi.nlm.nih.gov/protein/EHY60679.1) *[Exophiala dermatitidis NIH/UT8656]*  *pyruvate carboxylase* [*HMPREF1120_09185*](https://www.ncbi.nlm.nih.gov/protein/EHY61250.1) *[Exophiala dermatitidis NIH/UT8656]*  *hypothetical protein* [*HMPREF1120_08890*](https://www.ncbi.nlm.nih.gov/protein/EHY60948.1) *[Exophiala dermatitidis NIH/UT8656]*  *hypothetical protein* [*HMPREF1120_09031*](https://www.ncbi.nlm.nih.gov/protein/EHY61093.1) *[Exophiala dermatitidis NIH/UT8656]*  *hypothetical protein* [*HMPREF1120_09084*](https://www.ncbi.nlm.nih.gov/protein/EHY61148.1) *[Exophiala dermatitidis NIH/UT8656]*  *chitin synthase* [*HMPREF1120_08777*](https://www.ncbi.nlm.nih.gov/protein/EHY60833.1) *[Exophiala dermatitidis NIH/UT8656]*  *hypothetical protein* [*HMPREF1120_09220*](https://www.ncbi.nlm.nih.gov/protein/EHY61286.1) *[Exophiala dermatitidis NIH/UT8656]*  *ribonuclease H2 subunit A* [*HMPREF1120_09244*](https://www.ncbi.nlm.nih.gov/protein/EHY61310.1) *[Exophiala dermatitidis NIH/UT8656]* |
| *Ex5 & Ex18*  *148* | *MFS transporter, DHA2 family, methylenomycin A resistance protein* [*HMPREF1120_00012*](https://www.ncbi.nlm.nih.gov/protein/EHY51785.1) *[Exophiala dermatitidis NIH/UT8656]*  *DNA repair protein RAD50* [*HMPREF1120_04505*](https://www.ncbi.nlm.nih.gov/protein/EHY56423.1) *[Exophiala dermatitidis NIH/UT8656]*  *hypothetical protein* [*HMPREF1120_00457*](https://www.ncbi.nlm.nih.gov/protein/EHY52242.1) *[Exophiala dermatitidis NIH/UT8656]*  *inositol polyphosphate 5-phosphatase* [*HMPREF1120_01286*](https://www.ncbi.nlm.nih.gov/protein/EHY53086.1)*[Exophiala dermatitidis NIH/UT8656]*  *hypothetical protein* [*HMPREF1120_00487*](https://www.ncbi.nlm.nih.gov/protein/EHY52273.1) *[Exophiala dermatitidis NIH/UT8656]*  *hypothetical protein* [*HMPREF1120_00646*](https://www.ncbi.nlm.nih.gov/protein/EHY52434.1) *[Exophiala dermatitidis NIH/UT8656]*  *DEAD/DEAH box RNA helicase* [*HMPREF1120_00651*](https://www.ncbi.nlm.nih.gov/protein/EHY52439.1)*[Exophiala dermatitidis NIH/UT8656]*  *eukaryotic translation initiation factor 2 subunit gamma* [*HMPREF1120_00677*](https://www.ncbi.nlm.nih.gov/protein/EHY52465.1)  *[Exophiala dermatitidis NIH/UT8656]*  *cytochrome P450 oxidoreductase* [*HMPREF1120_01361*](https://www.ncbi.nlm.nih.gov/protein/EHY53163.1) *[Exophiala dermatitidis NIH/UT8656]*  *sulfite reductase (ferredoxin)* [*HMPREF1120_00943*](https://www.ncbi.nlm.nih.gov/protein/EHY52734.1)*[Exophiala dermatitidis NIH/UT8656]*  *glycerol ethanol, ferric requiring protein [Exophiala dermatitidis NIH/UT8656]*  *hypothetical protein* [*HMPREF1120_01089*](https://www.ncbi.nlm.nih.gov/protein/EHY52883.1) *[Exophiala dermatitidis NIH/UT8656]*  *hypothetical protein* [*HMPREF1120_01151*](https://www.ncbi.nlm.nih.gov/protein/EHY52950.1) *[Exophiala dermatitidis NIH/UT8656]*  *sulfite oxidase* [*HMPREF1120_05227*](https://www.ncbi.nlm.nih.gov/protein/EHY57179.1) *[Exophiala dermatitidis NIH/UT8656]*  *prolyl-tRNA synthetase* [*HMPREF1120_01354*](https://www.ncbi.nlm.nih.gov/protein/EHY53156.1) *[Exophiala dermatitidis NIH/UT8656]*  *cytochrome P450 oxidoreductase* [*HMPREF1120_01361*](https://www.ncbi.nlm.nih.gov/protein/EHY53163.1) *[Exophiala dermatitidis NIH/UT8656]*  *hypothetical protein* [*HMPREF1120_01471*](https://www.ncbi.nlm.nih.gov/protein/EHY53277.1) *[Exophiala dermatitidis NIH/UT8656]*  *serine/threonine kinase 16* [*HMPREF1120_06920*](https://www.ncbi.nlm.nih.gov/protein/EHY58918.1) *[Exophiala dermatitidis NIH/UT8656]*  *hypothetical protein* [*HMPREF1120_01961*](https://www.ncbi.nlm.nih.gov/protein/EHY53777.1) *[Exophiala dermatitidis NIH/UT8656]*  *hypothetical protein* [*HMPREF1120_02083*](https://www.ncbi.nlm.nih.gov/protein/EHY53903.1) *[Exophiala dermatitidis NIH/UT8656]*  *hypothetical protein* [*HMPREF1120_02190*](https://www.ncbi.nlm.nih.gov/protein/EHY54013.1) *[Exophiala dermatitidis NIH/UT8656]*  *MFS transporter, DHA1 family, multidrug resistance protein* [*HMPREF1120_09017*](https://www.ncbi.nlm.nih.gov/protein/EHY61079.1) *[Exophiala dermatitidis NIH/UT8656]*  *glutathione S-transferase* [*HMPREF1120_08143*](https://www.ncbi.nlm.nih.gov/protein/EHY60173.1) *[Exophiala dermatitidis NIH/UT8656]*  *nicotinate-nucleotide diphosphorylase (carboxylating)* [*HMPREF1120_02317*](https://www.ncbi.nlm.nih.gov/protein/EHY54142.1)  *[Exophiala dermatitidis NIH/UT8656]*  *phosphodiesterase/alkaline phosphatase D* [*HMPREF1120_02364*](https://www.ncbi.nlm.nih.gov/protein/EHY54190.1) *[Exophiala dermatitidis NIH/UT8656]*  *hypothetical protein* [*HMPREF1120_02386*](https://www.ncbi.nlm.nih.gov/protein/EHY54214.1) *[Exophiala dermatitidis NIH/UT8656]*  *Xanthine phosphoribosyltransferase 1* [*HMPREF1120_06110*](https://www.ncbi.nlm.nih.gov/protein/EHY58092.1) *[Exophiala dermatitidis NIH/UT8656]*  *Xanthine phosphoribosyltransferase 1* [*HMPREF1120_06110*](https://www.ncbi.nlm.nih.gov/protein/EHY58092.1) *[Exophiala dermatitidis NIH/UT8656]*  *hypothetical protein* [*HMPREF1120_02597*](https://www.ncbi.nlm.nih.gov/protein/EHY54428.1) *[Exophiala dermatitidis NIH/UT8656]*  *rRNA (cytosine-C5-)-methyltransferase nop2* [*HMPREF1120_02613*](https://www.ncbi.nlm.nih.gov/protein/EHY54444.1) *[Exophiala dermatitidis NIH/UT8656]*  *transformation/transcription domain-associated protein* [*HMPREF1120_02639*](https://www.ncbi.nlm.nih.gov/protein/EHY54471.1)  *[Exophiala dermatitidis NIH/UT8656]*  *Serine/threonine-protein phosphatase 2A 56 kDa regulatory subunit delta isoform* [*HMPREF1120_01344*](https://www.ncbi.nlm.nih.gov/protein/EHY53146.1) *[Exophiala dermatitidis NIH/UT8656]*  *hypothetical protein* [*HMPREF1120_02700*](https://www.ncbi.nlm.nih.gov/protein/EHY54532.1) *[Exophiala dermatitidis NIH/UT8656]*  *hypothetical protein* [*HMPREF1120_02708*](https://www.ncbi.nlm.nih.gov/protein/EHY54540.1) *[Exophiala dermatitidis NIH/UT8656]*  *hypothetical protein* [*HMPREF1120_02758*](https://www.ncbi.nlm.nih.gov/protein/EHY54590.1) *[Exophiala dermatitidis NIH/UT8656]*  *MC family mitochondrial carrier protein* [*HMPREF1120_08788*](https://www.ncbi.nlm.nih.gov/protein/EHY60844.1)*[Exophiala dermatitidis NIH/UT8656]*  *hypothetical protein* [*HMPREF1120_02784*](https://www.ncbi.nlm.nih.gov/protein/EHY54616.1) *[Exophiala dermatitidis NIH/UT8656]*  *hypothetical protein* [*HMPREF1120_02852*](https://www.ncbi.nlm.nih.gov/protein/EHY54687.1) *[Exophiala dermatitidis NIH/UT8656]*  *MOT1 - TATA-binding protein-associated factor* [*HMPREF1120_06808*](https://www.ncbi.nlm.nih.gov/protein/EHY58805.1)  *[Exophiala dermatitidis NIH/UT8656]*  *ING3 - Inhibitor of growth protein 3* [*HMPREF1120_05286*](https://www.ncbi.nlm.nih.gov/protein/EHY57240.1) *[Exophiala dermatitidis NIH/UT8656]*  *PDE - 3',5'-cyclic-nucleotide phosphodiesterase* [*HMPREF1120_05232*](https://www.ncbi.nlm.nih.gov/protein/EHY57184.1) *[Exophiala dermatitidis NIH/UT8656]*  *hypothetical protein* [*HMPREF1120_02947*](https://www.ncbi.nlm.nih.gov/protein/EHY54783.1) *[Exophiala dermatitidis NIH/UT8656]*  *hypothetical protein* [*HMPREF1120_03007*](https://www.ncbi.nlm.nih.gov/protein/EHY54845.1) *[Exophiala dermatitidis NIH/UT8656]*  *polyketide synthase* [*HMPREF1120_06570*](https://www.ncbi.nlm.nih.gov/protein/EHY58561.1)*[Exophiala dermatitidis NIH/UT8656]*  *VEI - velvet protein* [*HMPREF1120_06091*](https://www.ncbi.nlm.nih.gov/protein/EHY58073.1)*[Exophiala dermatitidis NIH/UT8656]*  *hypothetical protein* [*HMPREF1120_03359*](https://www.ncbi.nlm.nih.gov/protein/EHY55214.1) *[Exophiala dermatitidis NIH/UT8656]*  *salicylate hydroxylase* [*HMPREF1120_03459*](https://www.ncbi.nlm.nih.gov/protein/EHY55317.1)*[Exophiala dermatitidis NIH/UT8656]*  *hypothetical protein* [*HMPREF1120_03367*](https://www.ncbi.nlm.nih.gov/protein/EHY55222.1) *[Exophiala dermatitidis NIH/UT8656]*  *CPR6 - peptidyl-prolyl cis-trans isomerase cpr6* [*HMPREF1120_08126*](https://www.ncbi.nlm.nih.gov/protein/EHY60155.1)*[Exophiala dermatitidis NIH/UT8656]*  *gibberellin 2-oxidase* [*HMPREF1120_09208*](https://www.ncbi.nlm.nih.gov/protein/EHY61274.1)*[Exophiala dermatitidis NIH/UT8656]*  *hypothetical protein* [*HMPREF1120_03804*](https://www.ncbi.nlm.nih.gov/protein/EHY55678.1) *[Exophiala dermatitidis NIH/UT8656]*  *hypothetical protein* [*HMPREF1120_03909*](https://www.ncbi.nlm.nih.gov/protein/EHY55785.1) *[Exophiala dermatitidis NIH/UT8656]*  *hypothetical protein* [*HMPREF1120_03910*](https://www.ncbi.nlm.nih.gov/protein/EHY55786.1) *[Exophiala dermatitidis NIH/UT8656]*  *hypothetical protein* [*HMPREF1120_03928*](https://www.ncbi.nlm.nih.gov/protein/EHY55806.1) *[Exophiala dermatitidis NIH/UT8656]*  *amidohydrolase* [*HMPREF1120_03964*](https://www.ncbi.nlm.nih.gov/protein/EHY55847.1)*[Exophiala dermatitidis NIH/UT8656]*  *hypothetical protein* [*HMPREF1120_04101*](https://www.ncbi.nlm.nih.gov/protein/EHY55995.1) *[Exophiala dermatitidis NIH/UT8656]*  *MFS transporter, SP family, sugar:H+ symporter* [*HMPREF1120_06771*](https://www.ncbi.nlm.nih.gov/protein/EHY58768.1) *[Exophiala dermatitidis NIH/UT8656]*  *COP9 signalosome complex subunit 2* [*HMPREF1120_04182*](https://www.ncbi.nlm.nih.gov/protein/EHY56082.1)*[Exophiala dermatitidis NIH/UT8656]*  *hypothetical protein* [*HMPREF1120_04204*](https://www.ncbi.nlm.nih.gov/protein/EHY56104.1) *[Exophiala dermatitidis NIH/UT8656]*  *SIP3 - Putative sterol transfer protein* [*HMPREF1120_04207*](https://www.ncbi.nlm.nih.gov/protein/EHY56107.1)*[Exophiala dermatitidis NIH/UT8656]*  *hypothetical protein* [*HMPREF1120_04232*](https://www.ncbi.nlm.nih.gov/protein/EHY56135.1) *[Exophiala dermatitidis NIH/UT8656]*  *hypothetical protein* [*HMPREF1120_04314*](https://www.ncbi.nlm.nih.gov/protein/EHY56224.1) *[Exophiala dermatitidis NIH/UT8656]*  *hypothetical protein* [*HMPREF1120_04349*](https://www.ncbi.nlm.nih.gov/protein/EHY56262.1) *[Exophiala dermatitidis NIH/UT8656]*  *KAR3 - kinesin-like nuclear fusion protein [Exophiala dermatitidis NIH/UT8656]*  *hypothetical protein* [*HMPREF1120_04459*](https://www.ncbi.nlm.nih.gov/protein/EHY56377.1) *[Exophiala dermatitidis NIH/UT8656]*  *tyrosinase* [*HMPREF1120_04514*](https://www.ncbi.nlm.nih.gov/protein/EHY56432.1)*[Exophiala dermatitidis NIH/UT8656]*  *hypothetical protein* [*HMPREF1120_04584*](https://www.ncbi.nlm.nih.gov/protein/EHY56503.1) *[Exophiala dermatitidis NIH/UT8656]*  *ankyrin* [*HMPREF1120_08463*](https://www.ncbi.nlm.nih.gov/protein/EHY60507.1)*[Exophiala dermatitidis NIH/UT8656]*  *hypothetical protein* [*HMPREF1120_04645*](https://www.ncbi.nlm.nih.gov/protein/EHY56567.1) *[Exophiala dermatitidis NIH/UT8656]*  *hypothetical protein* [*HMPREF1120_04673*](https://www.ncbi.nlm.nih.gov/protein/EHY56597.1) *[Exophiala dermatitidis NIH/UT8656]*  *alkanesulfonate monooxygenase* [*HMPREF1120_08264*](https://www.ncbi.nlm.nih.gov/protein/EHY60297.1)*[Exophiala dermatitidis NIH/UT8656]*  *hypothetical protein* [*HMPREF1120_04695*](https://www.ncbi.nlm.nih.gov/protein/EHY56619.1) *[Exophiala dermatitidis NIH/UT8656]*  *hypothetical protein* [*HMPREF1120_04835*](https://www.ncbi.nlm.nih.gov/protein/EHY56769.1) *[Exophiala dermatitidis NIH/UT8656]*  *hypothetical protein* [*HMPREF1120_04991*](https://www.ncbi.nlm.nih.gov/protein/EHY56927.1) *[Exophiala dermatitidis NIH/UT8656]*  *4-coumarate-CoA ligase* [*HMPREF1120_09210*](https://www.ncbi.nlm.nih.gov/protein/EHY61276.1)*[Exophiala dermatitidis NIH/UT8656]*  *hypothetical protein* [*HMPREF1120_05054*](https://www.ncbi.nlm.nih.gov/protein/EHY56998.1) *[Exophiala dermatitidis NIH/UT8656]*  *KU70 - ATP-dependent DNA helicase II subunit 1* [*HMPREF1120_05117*](https://www.ncbi.nlm.nih.gov/protein/EHY57067.1)*[Exophiala dermatitidis NIH/UT8656]*  *hypothetical protein* [*HMPREF1120_05153*](https://www.ncbi.nlm.nih.gov/protein/EHY57103.1) *[Exophiala dermatitidis NIH/UT8656]*  *hypothetical protein* [*HMPREF1120_05151*](https://www.ncbi.nlm.nih.gov/protein/EHY57101.1) *[Exophiala dermatitidis NIH/UT8656]*  *hypothetical protein* [*HMPREF1120_05181*](https://www.ncbi.nlm.nih.gov/protein/EHY57132.1) *[Exophiala dermatitidis NIH/UT8656]*  *hypothetical protein* [*HMPREF1120_05217*](https://www.ncbi.nlm.nih.gov/protein/EHY57169.1) *[Exophiala dermatitidis NIH/UT8656]*  *dimethylaniline monooxygenase (N-oxide forming)* [*HMPREF1120_08742*](https://www.ncbi.nlm.nih.gov/protein/EHY60798.1)*[Exophiala dermatitidis NIH/UT8656]*  *hypothetical protein* [*HMPREF1120_05400*](https://www.ncbi.nlm.nih.gov/protein/EHY57359.1) *[Exophiala dermatitidis NIH/UT8656]*  *dihydrodipicolinate synthetase* [*HMPREF1120_09161*](https://www.ncbi.nlm.nih.gov/protein/EHY61225.1)*[Exophiala dermatitidis NIH/UT8656]*  *DNA repair protein RAD50* [*HMPREF1120_04505*](https://www.ncbi.nlm.nih.gov/protein/EHY56423.1) *[Exophiala dermatitidis NIH/UT8656]*  *hypothetical protein* [*HMPREF1120_05622*](https://www.ncbi.nlm.nih.gov/protein/EHY57593.1) *[Exophiala dermatitidis NIH/UT8656]*  *hypothetical protein* [*HMPREF1120_05658*](https://www.ncbi.nlm.nih.gov/protein/EHY57629.1) *[Exophiala dermatitidis NIH/UT8656]*  *PSY2 - Platinum sensitivity protein [Exophiala dermatitidis NIH/UT8656]*  *hypothetical protein* [*HMPREF1120_05728*](https://www.ncbi.nlm.nih.gov/protein/EHY57701.1) *[Exophiala dermatitidis NIH/UT8656]*  *SPT20 - Transcription factor spt20* [*HMPREF1120_05758*](https://www.ncbi.nlm.nih.gov/protein/EHY57731.1)*[Exophiala dermatitidis NIH/UT8656]*  *hypothetical protein* [*HMPREF1120_05809*](https://www.ncbi.nlm.nih.gov/protein/EHY57785.1) *[Exophiala dermatitidis NIH/UT8656]*  *ETF1 - elongation factor 2* [*HMPREF1120_05986*](https://www.ncbi.nlm.nih.gov/protein/EHY57966.1)*[Exophiala dermatitidis NIH/UT8656]*  *hypothetical protein* [*HMPREF1120_06093*](https://www.ncbi.nlm.nih.gov/protein/EHY58075.1) *[Exophiala dermatitidis NIH/UT8656]*  *RGT1 - Glucose-responsive transcription factor 1 [Exophiala dermatitidis NIH/UT8656]*  *hypothetical protein* [*HMPREF1120_06177*](https://www.ncbi.nlm.nih.gov/protein/EHY58163.1) *[Exophiala dermatitidis NIH/UT8656]*  *hypothetical protein* [*HMPREF1120_06361*](https://www.ncbi.nlm.nih.gov/protein/EHY58349.1) *[Exophiala dermatitidis NIH/UT8656]*  *37S ribosomal protein, mitochondrial [Exophiala dermatitidis NIH/UT8656]*  *hypothetical protein* [*HMPREF1120_06500*](https://www.ncbi.nlm.nih.gov/protein/EHY58490.1) *[Exophiala dermatitidis NIH/UT8656]*  *hypothetical protein* [*HMPREF1120_06642*](https://www.ncbi.nlm.nih.gov/protein/EHY58637.1) *[Exophiala dermatitidis NIH/UT8656]*  *MFS transporter, SP family, solute carrier family 2 (facilitated glucose transporter), member 2* [*HMPREF1120_06771*](https://www.ncbi.nlm.nih.gov/protein/EHY58768.1)  *[Exophiala dermatitidis NIH/UT8656]*  *hypothetical protein* [*HMPREF1120_06878*](https://www.ncbi.nlm.nih.gov/protein/EHY58876.1) *[Exophiala dermatitidis NIH/UT8656]*  *hypothetical protein* [*HMPREF1120_06971*](https://www.ncbi.nlm.nih.gov/protein/EHY58970.1) *[Exophiala dermatitidis NIH/UT8656]*  *transcription initiation factor TFIID subunit D2* [*MPREF1120_06984*](https://www.ncbi.nlm.nih.gov/protein/EHY58983.1)*[Exophiala dermatitidis NIH/UT8656]*  *L-galactose dehydrogenase* [*HMPREF1120_07000*](https://www.ncbi.nlm.nih.gov/protein/EHY59000.1)*[Exophiala dermatitidis NIH/UT8656]*  *FAS1_2 - beta subunit of fatty acid synthetase* [*HMPREF1120_07065*](https://www.ncbi.nlm.nih.gov/protein/EHY59066.1)*[Exophiala dermatitidis NIH/UT8656]*  *hypothetical protein* [*HMPREF1120_07085*](https://www.ncbi.nlm.nih.gov/protein/EHY59086.1) *[Exophiala dermatitidis NIH/UT8656]*  *3' exoribonuclease* [*HMPREF1120_08304*](https://www.ncbi.nlm.nih.gov/protein/EHY60338.1)*[Exophiala dermatitidis NIH/UT8656]*  *hypothetical protein* [*HMPREF1120_07255*](https://www.ncbi.nlm.nih.gov/protein/EHY59262.1) *[Exophiala dermatitidis NIH/UT8656]*  *hypothetical protein* [*HMPREF1120_07291*](https://www.ncbi.nlm.nih.gov/protein/EHY59299.1) *[Exophiala dermatitidis NIH/UT8656]*  *hypothetical protein* [*HMPREF1120_07306*](https://www.ncbi.nlm.nih.gov/protein/EHY59314.1) *[Exophiala dermatitidis NIH/UT8656]*  *hypothetical protein* [*HMPREF1120_07433*](https://www.ncbi.nlm.nih.gov/protein/EHY59443.1) *[Exophiala dermatitidis NIH/UT8656]*  *RAS2 - Ras GTPase* [*HMPREF1120_01421*](https://www.ncbi.nlm.nih.gov/protein/EHY53224.1)*[Exophiala dermatitidis NIH/UT8656]*  *hypothetical protein* [*HMPREF1120_07616*](https://www.ncbi.nlm.nih.gov/protein/EHY59631.1) *[Exophiala dermatitidis NIH/UT8656]*  *D-3-phosphoglycerate dehydrogenase* [*HMPREF1120_06805*](https://www.ncbi.nlm.nih.gov/protein/EHY58802.1)*[Exophiala dermatitidis NIH/UT8656]*  *ankyrin* [*HMPREF1120_08463*](https://www.ncbi.nlm.nih.gov/protein/EHY60507.1)*[Exophiala dermatitidis NIH/UT8656]*  *nuclear transcription factor Y, alpha* [*HMPREF1120_07714*](https://www.ncbi.nlm.nih.gov/protein/EHY59731.1)*[Exophiala dermatitidis NIH/UT8656]*  *CHS3 - chitin synthase class 3 [Exophiala dermatitidis NIH/UT8656]*  *salicylate hydroxylase* [*HMPREF1120_03459*](https://www.ncbi.nlm.nih.gov/protein/EHY55317.1)*[Exophiala dermatitidis NIH/UT8656]*  *hypothetical protein* [*HMPREF1120_06589*](https://www.ncbi.nlm.nih.gov/protein/EHY58580.1) *[Exophiala dermatitidis NIH/UT8656]*  *hypothetical protein* [*HMPREF1120_06584*](https://www.ncbi.nlm.nih.gov/protein/EHY58575.1) *[Exophiala dermatitidis NIH/UT8656]*  *queuine tRNA-ribosyltransferase* [*HMPREF1120_07977*](https://www.ncbi.nlm.nih.gov/protein/EHY60002.1)*[Exophiala dermatitidis NIH/UT8656]*  *hypothetical protein* [*HMPREF1120_07985*](https://www.ncbi.nlm.nih.gov/protein/EHY60010.1) *[Exophiala dermatitidis NIH/UT8656]*  *thiamin biosynthesis protein* [*HMPREF1120_07987*](https://www.ncbi.nlm.nih.gov/protein/EHY60012.1)*[Exophiala dermatitidis NIH/UT8656]*  *ABD1 - mRNA cap guanine-N7 methyltransferase* [*HMPREF1120_06541*](https://www.ncbi.nlm.nih.gov/protein/EHY58531.1)*[Exophiala dermatitidis NIH/UT8656]*  *hypothetical protein* [*HMPREF1120_08063*](https://www.ncbi.nlm.nih.gov/protein/EHY60091.1) *[Exophiala dermatitidis NIH/UT8656]*  *PAN2 - poly(A) specific ribonuclease [Exophiala dermatitidis NIH/UT8656]*  *DOA4 - ubiquitin specific protease* [*HMPREF1120_06573*](https://www.ncbi.nlm.nih.gov/protein/EHY58564.1)*[Exophiala dermatitidis NIH/UT8656]*  *amidase* [*HMPREF1120_09153*](https://www.ncbi.nlm.nih.gov/protein/EHY61217.1)*[Exophiala dermatitidis NIH/UT8656]*  *MEF2 - Ribosome-releasing factor 2, mitochondrial [Exophiala dermatitidis NIH/UT8656]*  *RGA2 - Rho-type gtpase-activating protein [Exophiala dermatitidis NIH/UT8656]*  *hypothetical protein* [*HMPREF1120_08236*](https://www.ncbi.nlm.nih.gov/protein/EHY60268.1) *[Exophiala dermatitidis NIH/UT8656]*  *hypothetical protein* [*HMPREF1120_08238*](https://www.ncbi.nlm.nih.gov/protein/EHY60270.1) *[Exophiala dermatitidis NIH/UT8656]*  *hypothetical protein* [*HMPREF1120_08408*](https://www.ncbi.nlm.nih.gov/protein/EHY60446.1) *[Exophiala dermatitidis NIH/UT8656]*  *cytochrome P450 oxidoreductase* [*HMPREF1120_01361*](https://www.ncbi.nlm.nih.gov/protein/EHY53163.1) *[Exophiala dermatitidis NIH/UT8656]*  *hypothetical protein* [*HMPREF1120_08425*](https://www.ncbi.nlm.nih.gov/protein/EHY60464.1) *[Exophiala dermatitidis NIH/UT8656]*  *hydrolase* [*HMPREF1120_08460*](https://www.ncbi.nlm.nih.gov/protein/EHY60504.1)*[Exophiala dermatitidis NIH/UT8656]*  *hypothetical protein* [*HMPREF1120_08629*](https://www.ncbi.nlm.nih.gov/protein/EHY60679.1) *[Exophiala dermatitidis NIH/UT8656]*  *pyruvate carboxylase* [*HMPREF1120_09185*](https://www.ncbi.nlm.nih.gov/protein/EHY61250.1) *[Exophiala dermatitidis NIH/UT8656]*  *branchpoint-bridging protein* [*HMPREF1120_08884*](https://www.ncbi.nlm.nih.gov/protein/EHY60941.1)*[Exophiala dermatitidis NIH/UT8656]*  *hypothetical protein* [*HMPREF1120_08890*](https://www.ncbi.nlm.nih.gov/protein/EHY60948.1) *[Exophiala dermatitidis NIH/UT8656]*  *hypothetical protein* [*HMPREF1120_09031*](https://www.ncbi.nlm.nih.gov/protein/EHY61093.1) *[Exophiala dermatitidis NIH/UT8656]*  *hypothetical protein* [*HMPREF1120_09084*](https://www.ncbi.nlm.nih.gov/protein/EHY61148.1) *[Exophiala dermatitidis NIH/UT8656]*  *chitin synthase* [*HMPREF1120_08777*](https://www.ncbi.nlm.nih.gov/protein/EHY60833.1)  *[Exophiala dermatitidis NIH/UT8656]*  *hypothetical protein* [*HMPREF1120_09220*](https://www.ncbi.nlm.nih.gov/protein/EHY61286.1) *[Exophiala dermatitidis NIH/UT8656]*  *ribonuclease H2 subunit A* [*HMPREF1120_09244*](https://www.ncbi.nlm.nih.gov/protein/EHY61310.1) *[Exophiala dermatitidis NIH/UT8656]* |
| *Ex5 & Ex20*  *111* | *DNA repair protein RAD50* [*HMPREF1120_04505*](https://www.ncbi.nlm.nih.gov/protein/EHY56423.1) *[Exophiala dermatitidis NIH/UT8656]*  *hypothetical protein* [*HMPREF1120_00144*](https://www.ncbi.nlm.nih.gov/protein/EHY51921.1) *[Exophiala dermatitidis NIH/UT8656]*  *hypothetical protein* [*HMPREF1120_00457*](https://www.ncbi.nlm.nih.gov/protein/EHY52242.1) *[Exophiala dermatitidis NIH/UT8656]*  *hypothetical protein* [*HMPREF1120_00646*](https://www.ncbi.nlm.nih.gov/protein/EHY52434.1) *[Exophiala dermatitidis NIH/UT8656]*  *eukaryotic translation initiation factor 2 subunit gamma* [*HMPREF1120_00677*](https://www.ncbi.nlm.nih.gov/protein/EHY52465.1)  *[Exophiala dermatitidis NIH/UT8656]*  *cytochrome P450 oxidoreductase* [*HMPREF1120_01361*](https://www.ncbi.nlm.nih.gov/protein/EHY53163.1) *[Exophiala dermatitidis NIH/UT8656]*  *hypothetical protein* [*HMPREF1120_00830*](https://www.ncbi.nlm.nih.gov/protein/EHY52619.1) *[Exophiala dermatitidis NIH/UT8656]*  *hypothetical protein* [*HMPREF1120_01089*](https://www.ncbi.nlm.nih.gov/protein/EHY52883.1) *[Exophiala dermatitidis NIH/UT8656]*  *hypothetical protein* [*HMPREF1120_01151*](https://www.ncbi.nlm.nih.gov/protein/EHY52950.1) *[Exophiala dermatitidis NIH/UT8656]*  *prolyl-tRNA synthetase* [*HMPREF1120_01354*](https://www.ncbi.nlm.nih.gov/protein/EHY53156.1) *[Exophiala dermatitidis NIH/UT8656]*  *hypothetical protein* [*HMPREF1120_01471*](https://www.ncbi.nlm.nih.gov/protein/EHY53277.1) *[Exophiala dermatitidis NIH/UT8656]*  *serine/threonine kinase 16* [*HMPREF1120_06920*](https://www.ncbi.nlm.nih.gov/protein/EHY58918.1) *[Exophiala dermatitidis NIH/UT8656]*  *hypothetical protein* [*HMPREF1120_02083*](https://www.ncbi.nlm.nih.gov/protein/EHY53903.1) *[Exophiala dermatitidis NIH/UT8656]*  *hypothetical protein* [*HMPREF1120_02190*](https://www.ncbi.nlm.nih.gov/protein/EHY54013.1) *[Exophiala dermatitidis NIH/UT8656]*  *glutathione S-transferase* [*HMPREF1120_08143*](https://www.ncbi.nlm.nih.gov/protein/EHY60173.1) *[Exophiala dermatitidis NIH/UT8656]*  *nicotinate-nucleotide diphosphorylase (carboxylating)* [*HMPREF1120_02317*](https://www.ncbi.nlm.nih.gov/protein/EHY54142.1)  *[Exophiala dermatitidis NIH/UT8656]*  *phosphodiesterase/alkaline phosphatase D* [*HMPREF1120_02364*](https://www.ncbi.nlm.nih.gov/protein/EHY54190.1) *[Exophiala dermatitidis NIH/UT8656]*  *Xanthine phosphoribosyltransferase 1* [*HMPREF1120_06110*](https://www.ncbi.nlm.nih.gov/protein/EHY58092.1) *[Exophiala dermatitidis NIH/UT8656]*  *Xanthine phosphoribosyltransferase 1* [*HMPREF1120_06110*](https://www.ncbi.nlm.nih.gov/protein/EHY58092.1) *[Exophiala dermatitidis NIH/UT8656]*  *hypothetical protein* [*HMPREF1120_02700*](https://www.ncbi.nlm.nih.gov/protein/EHY54532.1) *[Exophiala dermatitidis NIH/UT8656]*  *hypothetical protein* [*HMPREF1120_02708*](https://www.ncbi.nlm.nih.gov/protein/EHY54540.1) *[Exophiala dermatitidis NIH/UT8656]*  *hypothetical protein* [*HMPREF1120_02758*](https://www.ncbi.nlm.nih.gov/protein/EHY54590.1) *[Exophiala dermatitidis NIH/UT8656]*  *hypothetical protein* [*HMPREF1120_02784*](https://www.ncbi.nlm.nih.gov/protein/EHY54616.1) *[Exophiala dermatitidis NIH/UT8656]*  *MOT1 - TATA-binding protein-associated factor* [*HMPREF1120_06808*](https://www.ncbi.nlm.nih.gov/protein/EHY58805.1)  *[Exophiala dermatitidis NIH/UT8656]*  *ING3 - Inhibitor of growth protein 3* [*HMPREF1120_05286*](https://www.ncbi.nlm.nih.gov/protein/EHY57240.1) *[Exophiala dermatitidis NIH/UT8656]*  *hypothetical protein* [*HMPREF1120_02908*](https://www.ncbi.nlm.nih.gov/protein/EHY54743.1) *[Exophiala dermatitidis NIH/UT8656]*  *PDE2 - High-affinity cyclic AMP phosphodiesterase [Exophiala dermatitidis NIH/UT8656]*  *hypothetical protein* [*HMPREF1120_03003*](https://www.ncbi.nlm.nih.gov/protein/EHY54840.1) *[Exophiala dermatitidis NIH/UT8656]*  *hypothetical protein* [*HMPREF1120_03007*](https://www.ncbi.nlm.nih.gov/protein/EHY54845.1) *[Exophiala dermatitidis NIH/UT8656]*  *VEI - velvet protein* [*HMPREF1120_06091*](https://www.ncbi.nlm.nih.gov/protein/EHY58073.1)*[Exophiala dermatitidis NIH/UT8656]*  *hypothetical protein* [*HMPREF1120_03359*](https://www.ncbi.nlm.nih.gov/protein/EHY55214.1) *[Exophiala dermatitidis NIH/UT8656]*  *hypothetical protein* [*HMPREF1120_03513*](https://www.ncbi.nlm.nih.gov/protein/EHY55374.1) *[Exophiala dermatitidis NIH/UT8656]*  *gibberellin 2-oxidase* [*HMPREF1120_09208*](https://www.ncbi.nlm.nih.gov/protein/EHY61274.1)*[Exophiala dermatitidis NIH/UT8656]*  *hypothetical protein* [*HMPREF1120_03910*](https://www.ncbi.nlm.nih.gov/protein/EHY55786.1) *[Exophiala dermatitidis NIH/UT8656]*  *hypothetical protein* [*HMPREF1120_03911*](https://www.ncbi.nlm.nih.gov/protein/EHY55787.1) *[Exophiala dermatitidis NIH/UT8656]*  *hypothetical protein* [*HMPREF1120_03928*](https://www.ncbi.nlm.nih.gov/protein/EHY55806.1) *[Exophiala dermatitidis NIH/UT8656]*  *amidohydrolase* [*HMPREF1120_03964*](https://www.ncbi.nlm.nih.gov/protein/EHY55847.1)*[Exophiala dermatitidis NIH/UT8656]*  *hypothetical protein* [*HMPREF1120_04101*](https://www.ncbi.nlm.nih.gov/protein/EHY55995.1) *[Exophiala dermatitidis NIH/UT8656]*  *MFS transporter, SP family, sugar:H+ symporter* [*HMPREF1120_06771*](https://www.ncbi.nlm.nih.gov/protein/EHY58768.1)  *[Exophiala dermatitidis NIH/UT8656]*  *COP9 signalosome complex subunit 2* [*HMPREF1120_04182*](https://www.ncbi.nlm.nih.gov/protein/EHY56082.1)*[Exophiala dermatitidis NIH/UT8656]*  *hypothetical protein* [*HMPREF1120_04204*](https://www.ncbi.nlm.nih.gov/protein/EHY56104.1) *[Exophiala dermatitidis NIH/UT8656]*  *SIP3 - Putative sterol transfer protein* [*HMPREF1120_04207*](https://www.ncbi.nlm.nih.gov/protein/EHY56107.1)*[Exophiala dermatitidis NIH/UT8656]*  *hypothetical protein* [*HMPREF1120_04349*](https://www.ncbi.nlm.nih.gov/protein/EHY56262.1) *[Exophiala dermatitidis NIH/UT8656]*  *hypothetical protein* [*HMPREF1120_04459*](https://www.ncbi.nlm.nih.gov/protein/EHY56377.1) *[Exophiala dermatitidis NIH/UT8656]*  *tyrosinase* [*HMPREF1120_04514*](https://www.ncbi.nlm.nih.gov/protein/EHY56432.1)*[Exophiala dermatitidis NIH/UT8656]*  *hypothetical protein* [*HMPREF1120_04584*](https://www.ncbi.nlm.nih.gov/protein/EHY56503.1) *[Exophiala dermatitidis NIH/UT8656]*  *ankyrin* [*HMPREF1120_08463*](https://www.ncbi.nlm.nih.gov/protein/EHY60507.1)*[Exophiala dermatitidis NIH/UT8656]*  *hypothetical protein* [*HMPREF1120_04645*](https://www.ncbi.nlm.nih.gov/protein/EHY56567.1) *[Exophiala dermatitidis NIH/UT8656]*  *adenosinetriphosphatase* [*HMPREF1120_09246*](https://www.ncbi.nlm.nih.gov/protein/EHY61312.1) *[Exophiala dermatitidis NIH/UT8656]*  *hypothetical protein* [*HMPREF1120_04673*](https://www.ncbi.nlm.nih.gov/protein/EHY56597.1) *[Exophiala dermatitidis NIH/UT8656]*  *alkanesulfonate monooxygenase* [*HMPREF1120_08264*](https://www.ncbi.nlm.nih.gov/protein/EHY60297.1)*[Exophiala dermatitidis NIH/UT8656]*  *hypothetical protein* [*HMPREF1120_04835*](https://www.ncbi.nlm.nih.gov/protein/EHY56769.1) *[Exophiala dermatitidis NIH/UT8656]*  *hypothetical protein* [*HMPREF1120_04991*](https://www.ncbi.nlm.nih.gov/protein/EHY56927.1) *[Exophiala dermatitidis NIH/UT8656]*  *4-coumarate-CoA ligase* [*HMPREF1120_09210*](https://www.ncbi.nlm.nih.gov/protein/EHY61276.1)*[Exophiala dermatitidis NIH/UT8656]*  *hypothetical protein* [*HMPREF1120_05087*](https://www.ncbi.nlm.nih.gov/protein/EHY57035.1) *[Exophiala dermatitidis NIH/UT8656]*  *hypothetical protein* [*HMPREF1120_05151*](https://www.ncbi.nlm.nih.gov/protein/EHY57101.1) *[Exophiala dermatitidis NIH/UT8656]*  *hypothetical protein* [*HMPREF1120_05217*](https://www.ncbi.nlm.nih.gov/protein/EHY57169.1) *[Exophiala dermatitidis NIH/UT8656]*  *dimethylaniline monooxygenase (N-oxide forming)* [*HMPREF1120_08742*](https://www.ncbi.nlm.nih.gov/protein/EHY60798.1)*[Exophiala dermatitidis NIH/UT8656]*  *hypothetical protein* [*HMPREF1120_05393*](https://www.ncbi.nlm.nih.gov/protein/EHY57352.1) *[Exophiala dermatitidis NIH/UT8656]*  *hypothetical protein* [*HMPREF1120_05400*](https://www.ncbi.nlm.nih.gov/protein/EHY57359.1) *[Exophiala dermatitidis NIH/UT8656]*  *dihydrodipicolinate synthetase* [*HMPREF1120_09161*](https://www.ncbi.nlm.nih.gov/protein/EHY61225.1)*[Exophiala dermatitidis NIH/UT8656]*  *hypothetical protein* [*HMPREF1120_05622*](https://www.ncbi.nlm.nih.gov/protein/EHY57593.1) *[Exophiala dermatitidis NIH/UT8656]*  *hypothetical protein* [*HMPREF1120_05658*](https://www.ncbi.nlm.nih.gov/protein/EHY57629.1) *[Exophiala dermatitidis NIH/UT8656]*  *PSY2 - platinum sensitivity protein [Exophiala dermatitidis NIH/UT8656]*  *hypothetical protein* [*HMPREF1120_05728*](https://www.ncbi.nlm.nih.gov/protein/EHY57701.1) *[Exophiala dermatitidis NIH/UT8656]*  *SPT20 - Transcription factor spt20* [*HMPREF1120_05758*](https://www.ncbi.nlm.nih.gov/protein/EHY57731.1)*[Exophiala dermatitidis NIH/UT8656]*  *hypothetical protein* [*HMPREF1120_05809*](https://www.ncbi.nlm.nih.gov/protein/EHY57785.1) *[Exophiala dermatitidis NIH/UT8656]*  *ETF1 - elongation factor 2* [*HMPREF1120_05986*](https://www.ncbi.nlm.nih.gov/protein/EHY57966.1)*[Exophiala dermatitidis NIH/UT8656]*  *hypothetical protein* [*HMPREF1120_06093*](https://www.ncbi.nlm.nih.gov/protein/EHY58075.1) *[Exophiala dermatitidis NIH/UT8656]*  *hypothetical protein* [*HMPREF1120_06177*](https://www.ncbi.nlm.nih.gov/protein/EHY58163.1) *[Exophiala dermatitidis NIH/UT8656]*  *hypothetical protein* [*HMPREF1120_06361*](https://www.ncbi.nlm.nih.gov/protein/EHY58349.1) *[Exophiala dermatitidis NIH/UT8656]*  *37S ribosomal protein, mitochondrial [Exophiala dermatitidis NIH/UT8656]*  *hypothetical protein* [*HMPREF1120_06500*](https://www.ncbi.nlm.nih.gov/protein/EHY58490.1) *[Exophiala dermatitidis NIH/UT8656]*  *hypothetical protein* [*HMPREF1120_06642*](https://www.ncbi.nlm.nih.gov/protein/EHY58637.1) *[Exophiala dermatitidis NIH/UT8656]*  *hypothetical protein* [*HMPREF1120_06971*](https://www.ncbi.nlm.nih.gov/protein/EHY58970.1) *[Exophiala dermatitidis NIH/UT8656]*  *transcription initiation factor TFIID subunit D2* [*MPREF1120_06984*](https://www.ncbi.nlm.nih.gov/protein/EHY58983.1)*[Exophiala dermatitidis NIH/UT8656]*  *L-galactose dehydrogenase* [*HMPREF1120_07000*](https://www.ncbi.nlm.nih.gov/protein/EHY59000.1)*[Exophiala dermatitidis NIH/UT8656]*  *hypothetical protein* [*HMPREF1120_07085*](https://www.ncbi.nlm.nih.gov/protein/EHY59086.1) *[Exophiala dermatitidis NIH/UT8656]*  *3' exoribonuclease* [*HMPREF1120_08304*](https://www.ncbi.nlm.nih.gov/protein/EHY60338.1)*[Exophiala dermatitidis NIH/UT8656]*  *hypothetical protein* [*HMPREF1120_07255*](https://www.ncbi.nlm.nih.gov/protein/EHY59262.1) *[Exophiala dermatitidis NIH/UT8656]*  *hypothetical protein* [*HMPREF1120_07291*](https://www.ncbi.nlm.nih.gov/protein/EHY59299.1) *[Exophiala dermatitidis NIH/UT8656]*  *hypothetical protein* [*HMPREF1120_07306*](https://www.ncbi.nlm.nih.gov/protein/EHY59314.1) *[Exophiala dermatitidis NIH/UT8656]*  *hypothetical protein* [*HMPREF1120_07433*](https://www.ncbi.nlm.nih.gov/protein/EHY59443.1) *[Exophiala dermatitidis NIH/UT8656]*  *RAS2 - Ras GTPase* [*HMPREF1120_01421*](https://www.ncbi.nlm.nih.gov/protein/EHY53224.1)*[Exophiala dermatitidis NIH/UT8656]*  *hypothetical protein* [*HMPREF1120_07616*](https://www.ncbi.nlm.nih.gov/protein/EHY59631.1) *[Exophiala dermatitidis NIH/UT8656]*  *D-3-phosphoglycerate dehydrogenase* [*HMPREF1120_06805*](https://www.ncbi.nlm.nih.gov/protein/EHY58802.1)*[Exophiala dermatitidis NIH/UT8656]*  *ankyrin* [*HMPREF1120_08463*](https://www.ncbi.nlm.nih.gov/protein/EHY60507.1)*[Exophiala dermatitidis NIH/UT8656]*  *nuclear transcription factor Y, alpha* [*HMPREF1120_07714*](https://www.ncbi.nlm.nih.gov/protein/EHY59731.1)*[Exophiala dermatitidis NIH/UT8656]*  *salicylate hydroxylase* [*HMPREF1120_03459*](https://www.ncbi.nlm.nih.gov/protein/EHY55317.1)*[Exophiala dermatitidis NIH/UT8656]*  *hypothetical protein* [*HMPREF1120_06584*](https://www.ncbi.nlm.nih.gov/protein/EHY58575.1) *[Exophiala dermatitidis NIH/UT8656]*  *queuine tRNA-ribosyltransferase* [*HMPREF1120_07977*](https://www.ncbi.nlm.nih.gov/protein/EHY60002.1)*[Exophiala dermatitidis NIH/UT8656]*  *hypothetical protein* [*HMPREF1120_07985*](https://www.ncbi.nlm.nih.gov/protein/EHY60010.1) *[Exophiala dermatitidis NIH/UT8656]*  *thiamin biosynthesis protein* [*HMPREF1120_07987*](https://www.ncbi.nlm.nih.gov/protein/EHY60012.1)*[Exophiala dermatitidis NIH/UT8656]*  *ABD1 - mRNA cap guanine-N7 methyltransferase* [*HMPREF1120_06541*](https://www.ncbi.nlm.nih.gov/protein/EHY58531.1)*[Exophiala dermatitidis NIH/UT8656]*  *PAN2 - poly(A) specific ribonuclease [Exophiala dermatitidis NIH/UT8656]*  *DOA4 - ubiquitin specific protease* [*HMPREF1120_06573*](https://www.ncbi.nlm.nih.gov/protein/EHY58564.1)*[Exophiala dermatitidis NIH/UT8656]*  *VPS27 - vacuolar protein sorting associated protein 27 [Exophiala dermatitidis NIH/UT8656]*  *MEF2 - Ribosome-releasing factor 2, mitochondrial [Exophiala dermatitidis NIH/UT8656]*  *RGA2 - Rho-type gtpase-activating protein [Exophiala dermatitidis NIH/UT8656]*  *hypothetical protein* [*HMPREF1120_08236*](https://www.ncbi.nlm.nih.gov/protein/EHY60268.1) *[Exophiala dermatitidis NIH/UT8656]*  *cytochrome P450 oxidoreductase* [*HMPREF1120_01361*](https://www.ncbi.nlm.nih.gov/protein/EHY53163.1) *[Exophiala dermatitidis NIH/UT8656]*  *hypothetical protein* [*HMPREF1120_08425*](https://www.ncbi.nlm.nih.gov/protein/EHY60464.1) *[Exophiala dermatitidis NIH/UT8656]*  *hydrolase* [*HMPREF1120_08460*](https://www.ncbi.nlm.nih.gov/protein/EHY60504.1)*[Exophiala dermatitidis NIH/UT8656]*  *hypothetical protein* [*HMPREF1120_08629*](https://www.ncbi.nlm.nih.gov/protein/EHY60679.1) *[Exophiala dermatitidis NIH/UT8656]*  *pyruvate carboxylase* [*HMPREF1120_09185*](https://www.ncbi.nlm.nih.gov/protein/EHY61250.1) *[Exophiala dermatitidis NIH/UT8656]*  *hypothetical protein* [*HMPREF1120_08890*](https://www.ncbi.nlm.nih.gov/protein/EHY60948.1) *[Exophiala dermatitidis NIH/UT8656]*  *hypothetical protein* [*HMPREF1120_09084*](https://www.ncbi.nlm.nih.gov/protein/EHY61148.1) *[Exophiala dermatitidis NIH/UT8656]*  *chitin synthase* [*HMPREF1120_08777*](https://www.ncbi.nlm.nih.gov/protein/EHY60833.1) *[Exophiala dermatitidis NIH/UT8656]*  *hypothetical protein* [*HMPREF1120_09220*](https://www.ncbi.nlm.nih.gov/protein/EHY61286.1) *[Exophiala dermatitidis NIH/UT8656]*  *ribonuclease H2 subunit A* [*HMPREF1120_09244*](https://www.ncbi.nlm.nih.gov/protein/EHY61310.1)*[Exophiala dermatitidis NIH/UT8656]* |
| *Ex5 & Ex21*  *72* | *hypothetical protein* [*HMPREF1120_00144*](https://www.ncbi.nlm.nih.gov/protein/EHY51921.1) *[Exophiala dermatitidis NIH/UT8656]*  *hypothetical protein* [*HMPREF1120_00457*](https://www.ncbi.nlm.nih.gov/protein/EHY52242.1) *[Exophiala dermatitidis NIH/UT8656]*  *hypothetical protein* [*HMPREF1120_00646*](https://www.ncbi.nlm.nih.gov/protein/EHY52434.1) *[Exophiala dermatitidis NIH/UT8656]*  *eukaryotic translation initiation factor 2 subunit gamma* [*HMPREF1120_00677*](https://www.ncbi.nlm.nih.gov/protein/EHY52465.1)  *[Exophiala dermatitidis NIH/UT8656]*  *cytochrome P450 oxidoreductase* [*HMPREF1120_01361*](https://www.ncbi.nlm.nih.gov/protein/EHY53163.1) *[Exophiala dermatitidis NIH/UT8656]*  *H(+)-transporting V1 sector ATPase subunit H* [*HMPREF1120_00992*](https://www.ncbi.nlm.nih.gov/protein/EHY52784.1)*[Exophiala dermatitidis NIH/UT8656]*  *hypothetical protein* [*HMPREF1120_01089*](https://www.ncbi.nlm.nih.gov/protein/EHY52883.1) *[Exophiala dermatitidis NIH/UT8656]*  *sulfite oxidase* [*HMPREF1120_05227*](https://www.ncbi.nlm.nih.gov/protein/EHY57179.1) *[Exophiala dermatitidis NIH/UT8656]*  *prolyl-tRNA synthetase* [*HMPREF1120_01354*](https://www.ncbi.nlm.nih.gov/protein/EHY53156.1) *[Exophiala dermatitidis NIH/UT8656]*  *cytochrome P450 oxidoreductase* [*HMPREF1120_01361*](https://www.ncbi.nlm.nih.gov/protein/EHY53163.1) *[Exophiala dermatitidis NIH/UT8656]*  *hypothetical protein* [*HMPREF1120_01814*](https://www.ncbi.nlm.nih.gov/protein/EHY53626.1) *[Exophiala dermatitidis NIH/UT8656]*  *serine/threonine kinase 16* [*HMPREF1120_06920*](https://www.ncbi.nlm.nih.gov/protein/EHY58918.1) *[Exophiala dermatitidis NIH/UT8656]*  *G4 quadruplex nucleic acid binding protein [Exophiala dermatitidis NIH/UT8656]*  *hypothetical protein* [*HMPREF1120_02190*](https://www.ncbi.nlm.nih.gov/protein/EHY54013.1) *[Exophiala dermatitidis NIH/UT8656]*  *nicotinate-nucleotide diphosphorylase (carboxylating)* [*HMPREF1120_02317*](https://www.ncbi.nlm.nih.gov/protein/EHY54142.1)  *[Exophiala dermatitidis NIH/UT8656]*  *phosphodiesterase/alkaline phosphatase D* [*HMPREF1120_02364*](https://www.ncbi.nlm.nih.gov/protein/EHY54190.1) *[Exophiala dermatitidis NIH/UT8656]*  *Xanthine phosphoribosyltransferase 1* [*HMPREF1120_06110*](https://www.ncbi.nlm.nih.gov/protein/EHY58092.1) *[Exophiala dermatitidis NIH/UT8656]*  *Xanthine phosphoribosyltransferase 1* [*HMPREF1120_06110*](https://www.ncbi.nlm.nih.gov/protein/EHY58092.1) *[Exophiala dermatitidis NIH/UT8656]*  *Serine/threonine-protein phosphatase 2A 56 kDa regulatory subunit delta isoform* [*HMPREF1120_01344*](https://www.ncbi.nlm.nih.gov/protein/EHY53146.1) *[Exophiala dermatitidis NIH/UT8656]*  *hypothetical protein* [*HMPREF1120_02784*](https://www.ncbi.nlm.nih.gov/protein/EHY54616.1) *[Exophiala dermatitidis NIH/UT8656]*  *ING3 - Inhibitor of growth protein 3* [*HMPREF1120_05286*](https://www.ncbi.nlm.nih.gov/protein/EHY57240.1) *[Exophiala dermatitidis NIH/UT8656]*  *hypothetical protein* [*HMPREF1120_02908*](https://www.ncbi.nlm.nih.gov/protein/EHY54743.1) *[Exophiala dermatitidis NIH/UT8656]*  *PDE2 - High-affinity cyclic AMP phosphodiesterase [Exophiala dermatitidis NIH/UT8656]*  *gibberellin 2-oxidase* [*HMPREF1120_09208*](https://www.ncbi.nlm.nih.gov/protein/EHY61274.1)*[Exophiala dermatitidis NIH/UT8656]*  *hypothetical protein* [*HMPREF1120_04101*](https://www.ncbi.nlm.nih.gov/protein/EHY55995.1) *[Exophiala dermatitidis NIH/UT8656]*  *MFS transporter, SP family, sugar:H+ symporter* [*HMPREF1120_06771*](https://www.ncbi.nlm.nih.gov/protein/EHY58768.1) *[Exophiala dermatitidis NIH/UT8656]*  *SIP3 - Putative sterol transfer protein* [*HMPREF1120_04207*](https://www.ncbi.nlm.nih.gov/protein/EHY56107.1)*[Exophiala dermatitidis NIH/UT8656]*  *hypothetical protein* [*HMPREF1120_04349*](https://www.ncbi.nlm.nih.gov/protein/EHY56262.1) *[Exophiala dermatitidis NIH/UT8656]*  *hypothetical protein* [*HMPREF1120_04459*](https://www.ncbi.nlm.nih.gov/protein/EHY56377.1) *[Exophiala dermatitidis NIH/UT8656]*  *ankyrin* [*HMPREF1120_08463*](https://www.ncbi.nlm.nih.gov/protein/EHY60507.1)*[Exophiala dermatitidis NIH/UT8656]*  *hypothetical protein* [*HMPREF1120_04991*](https://www.ncbi.nlm.nih.gov/protein/EHY56927.1) *[Exophiala dermatitidis NIH/UT8656]*  *arginyl-tRNA synthetase* [*HMPREF1120_08595*](https://www.ncbi.nlm.nih.gov/protein/EHY60643.1)*[Exophiala dermatitidis NIH/UT8656]*  *hypothetical protein* [*HMPREF1120_05087*](https://www.ncbi.nlm.nih.gov/protein/EHY57035.1) *[Exophiala dermatitidis NIH/UT8656]*  *hypothetical protein* [*HMPREF1120_05151*](https://www.ncbi.nlm.nih.gov/protein/EHY57101.1) *[Exophiala dermatitidis NIH/UT8656]*  *hypothetical protein* [*HMPREF1120_05217*](https://www.ncbi.nlm.nih.gov/protein/EHY57169.1) *[Exophiala dermatitidis NIH/UT8656]*  *dihydrodipicolinate synthetase* [*HMPREF1120_09161*](https://www.ncbi.nlm.nih.gov/protein/EHY61225.1)*[Exophiala dermatitidis NIH/UT8656]*  *hypothetical protein* [*HMPREF1120_05658*](https://www.ncbi.nlm.nih.gov/protein/EHY57629.1) *[Exophiala dermatitidis NIH/UT8656]*  *ETF1 - elongation factor 2* [*HMPREF1120_05986*](https://www.ncbi.nlm.nih.gov/protein/EHY57966.1)*[Exophiala dermatitidis NIH/UT8656]*  *hypothetical protein* [*HMPREF1120_06093*](https://www.ncbi.nlm.nih.gov/protein/EHY58075.1) *[Exophiala dermatitidis NIH/UT8656]*  *hypothetical protein* [*HMPREF1120_06177*](https://www.ncbi.nlm.nih.gov/protein/EHY58163.1) *[Exophiala dermatitidis NIH/UT8656]*  *hypothetical protein* [*HMPREF1120_06361*](https://www.ncbi.nlm.nih.gov/protein/EHY58349.1) *[Exophiala dermatitidis NIH/UT8656]*  *hypothetical protein* [*HMPREF1120_06500*](https://www.ncbi.nlm.nih.gov/protein/EHY58490.1) *[Exophiala dermatitidis NIH/UT8656]*  *hypothetical protein* [*HMPREF1120_06642*](https://www.ncbi.nlm.nih.gov/protein/EHY58637.1) *[Exophiala dermatitidis NIH/UT8656]*  *hypothetical protein* [*HMPREF1120_06852*](https://www.ncbi.nlm.nih.gov/protein/EHY58850.1) *[Exophiala dermatitidis NIH/UT8656]*  *transcription initiation factor TFIID subunit D2* [*MPREF1120_06984*](https://www.ncbi.nlm.nih.gov/protein/EHY58983.1)*[Exophiala dermatitidis NIH/UT8656]*  *L-galactose dehydrogenase* [*HMPREF1120_07000*](https://www.ncbi.nlm.nih.gov/protein/EHY59000.1)*[Exophiala dermatitidis NIH/UT8656]*  *hypothetical protein* [*HMPREF1120_07085*](https://www.ncbi.nlm.nih.gov/protein/EHY59086.1) *[Exophiala dermatitidis NIH/UT8656]*  *3' exoribonuclease* [*HMPREF1120_08304*](https://www.ncbi.nlm.nih.gov/protein/EHY60338.1)*[Exophiala dermatitidis NIH/UT8656]*  *hypothetical protein* [*HMPREF1120_07255*](https://www.ncbi.nlm.nih.gov/protein/EHY59262.1) *[Exophiala dermatitidis NIH/UT8656]*  *hypothetical protein* [*HMPREF1120_07291*](https://www.ncbi.nlm.nih.gov/protein/EHY59299.1) *[Exophiala dermatitidis NIH/UT8656]*  *hypothetical protein* [*HMPREF1120_07306*](https://www.ncbi.nlm.nih.gov/protein/EHY59314.1) *[Exophiala dermatitidis NIH/UT8656]*  *hypothetical protein* [*HMPREF1120_07433*](https://www.ncbi.nlm.nih.gov/protein/EHY59443.1) *[Exophiala dermatitidis NIH/UT8656]*  *hypothetical protein* [*HMPREF1120_07571*](https://www.ncbi.nlm.nih.gov/protein/EHY59586.1) *[Exophiala dermatitidis NIH/UT8656]*  *hypothetical protein* [*HMPREF1120_07589*](https://www.ncbi.nlm.nih.gov/protein/EHY59604.1) *[Exophiala dermatitidis NIH/UT8656]*  *D-3-phosphoglycerate dehydrogenase* [*HMPREF1120_06805*](https://www.ncbi.nlm.nih.gov/protein/EHY58802.1)*[Exophiala dermatitidis NIH/UT8656]*  *ABD1 - mRNA cap guanine-N7 methyltransferase* [*HMPREF1120_06541*](https://www.ncbi.nlm.nih.gov/protein/EHY58531.1)*[Exophiala dermatitidis NIH/UT8656]*  *PAN2 - poly(A) specific ribonuclease [Exophiala dermatitidis NIH/UT8656]*  *DOA4 - ubiquitin specific protease* [*HMPREF1120_06573*](https://www.ncbi.nlm.nih.gov/protein/EHY58564.1)*[Exophiala dermatitidis NIH/UT8656]*  *RGA2 - Rho-type gtpase-activating protein [Exophiala dermatitidis NIH/UT8656]*  *hypothetical protein* [*HMPREF1120_08236*](https://www.ncbi.nlm.nih.gov/protein/EHY60268.1) *[Exophiala dermatitidis NIH/UT8656]*  *cytochrome P450 oxidoreductase* [*HMPREF1120_01361*](https://www.ncbi.nlm.nih.gov/protein/EHY53163.1) *[Exophiala dermatitidis NIH/UT8656]*  *hypothetical protein* [*HMPREF1120_08425*](https://www.ncbi.nlm.nih.gov/protein/EHY60464.1) *[Exophiala dermatitidis NIH/UT8656]*  *hydrolase* [*HMPREF1120_08460*](https://www.ncbi.nlm.nih.gov/protein/EHY60504.1)*[Exophiala dermatitidis NIH/UT8656]*  *hypothetical protein* [*HMPREF1120_08629*](https://www.ncbi.nlm.nih.gov/protein/EHY60679.1) *[Exophiala dermatitidis NIH/UT8656]*  *VMA2 - Vacuolar ATP synthase subunit B* [*HMPREF1120_08721*](https://www.ncbi.nlm.nih.gov/protein/EHY60777.1) *[Exophiala dermatitidis NIH/UT8656]*  *pyruvate carboxylase* [*HMPREF1120_09185*](https://www.ncbi.nlm.nih.gov/protein/EHY61250.1)*[Exophiala dermatitidis NIH/UT8656]*  *hypothetical protein* [*HMPREF1120_08890*](https://www.ncbi.nlm.nih.gov/protein/EHY60948.1) *[Exophiala dermatitidis NIH/UT8656]*  *hypothetical protein HMPREF1120_09050 [Exophiala dermatitidis NIH/UT8656]*  *hypothetical protein* [*HMPREF1120_09084*](https://www.ncbi.nlm.nih.gov/protein/EHY61148.1) *[Exophiala dermatitidis NIH/UT8656]*  *chitin synthase* [*HMPREF1120_08777*](https://www.ncbi.nlm.nih.gov/protein/EHY60833.1) *[Exophiala dermatitidis NIH/UT8656]*  *hypothetical protein* [*HMPREF1120_09220*](https://www.ncbi.nlm.nih.gov/protein/EHY61286.1) *[Exophiala dermatitidis NIH/UT8656]* |
| *Ex9 & Ex13*  *27* | *APL5 - AP-3 complex subunit delta [Exophiala dermatitidis NIH/UT8656]*  *G2/mitotic-specific cyclin 3/4 [Exophiala dermatitidis NIH/UT8656]*  *cytochrome P450 oxidoreductase* [*HMPREF1120_01361*](https://www.ncbi.nlm.nih.gov/protein/EHY53163.1) *[Exophiala dermatitidis NIH/UT8656]*  *DEAD box RNA helicase HelA* [*HMPREF1120_02010*](https://www.ncbi.nlm.nih.gov/protein/EHY53828.1) *[Exophiala dermatitidis NIH/UT8656]*  *hypothetical protein* [*HMPREF1120_02190*](https://www.ncbi.nlm.nih.gov/protein/EHY54013.1) *[Exophiala dermatitidis NIH/UT8656]*  *MFS transporter, DHA1 family, multidrug resistance protein* [*HMPREF1120_09017*](https://www.ncbi.nlm.nih.gov/protein/EHY61079.1)  *[Exophiala dermatitidis NIH/UT8656]*  *hypothetical protein* [*HMPREF1120_02556*](https://www.ncbi.nlm.nih.gov/protein/EHY54387.1) *[Exophiala dermatitidis NIH/UT8656]*  *hypothetical protein* [*HMPREF1120_02949*](https://www.ncbi.nlm.nih.gov/protein/EHY54785.1) *[Exophiala dermatitidis NIH/UT8656]*  *hypothetical protein* [*HMPREF1120_03367*](https://www.ncbi.nlm.nih.gov/protein/EHY55222.1) *[Exophiala dermatitidis NIH/UT8656]*  *VPS41 - Vacuolar protein sorting-associated protein 41[Exophiala dermatitidis NIH/UT8656]*  *MFS transporter, SP family, sugar:H+ symporter* [*HMPREF1120_06771*](https://www.ncbi.nlm.nih.gov/protein/EHY58768.1) *[Exophiala dermatitidis NIH/UT8656]*  *hypothetical protein* [*HMPREF1120_04659*](https://www.ncbi.nlm.nih.gov/protein/EHY56583.1) *[Exophiala dermatitidis NIH/UT8656]*  *DNA repair protein RAD50* [*HMPREF1120_04505*](https://www.ncbi.nlm.nih.gov/protein/EHY56423.1) *[Exophiala dermatitidis NIH/UT8656]*  *biphenyl-2,3-diol 1,2-dioxygenase, variant* [*HMPREF1120_05880*](https://www.ncbi.nlm.nih.gov/protein/EHY57856.1) *[Exophiala dermatitidis NIH/UT8656]*  *chloride channel 3 [Exophiala dermatitidis NIH/UT8656]*  *hypothetical protein* [*HMPREF1120_06002*](https://www.ncbi.nlm.nih.gov/protein/EHY57982.1) *[Exophiala dermatitidis NIH/UT8656]*  *hypothetical protein* [*HMPREF1120_06122*](https://www.ncbi.nlm.nih.gov/protein/EHY58104.1) *[Exophiala dermatitidis NIH/UT8656]*  *hypothetical protein* [*HMPREF1120_06336*](https://www.ncbi.nlm.nih.gov/protein/EHY58324.1) *[Exophiala dermatitidis NIH/UT8656]*  *ISU1 - iron-binding protein* [*HMPREF1120_06751*](https://www.ncbi.nlm.nih.gov/protein/EHY58748.1) *[Exophiala dermatitidis NIH/UT8656]*  *LBA1 - Regulator of nonsense transcripts 1-like protein [Exophiala dermatitidis NIH/UT8656]*  *hypothetical protein* [*HMPREF1120_07292*](https://www.ncbi.nlm.nih.gov/protein/EHY59300.1) *[Exophiala dermatitidis NIH/UT8656]*  *adenosinetriphosphatase* [*HMPREF1120_09246*](https://www.ncbi.nlm.nih.gov/protein/EHY61312.1) *[Exophiala dermatitidis NIH/UT8656]*  *TPC1 - mitochondrial thiamine pyrophosphate transporter* [*HMPREF1120_07619*](https://www.ncbi.nlm.nih.gov/protein/EHY59634.1) *[Exophiala dermatitidis NIH/UT8656]*  *MFS transporter, SIT family, siderophore-iron:H+ symporter* [*HMPREF1120_07838*](https://www.ncbi.nlm.nih.gov/protein/EHY59858.1) *[Exophiala dermatitidis NIH/UT8656]*  *hypothetical protein* [*HMPREF1120_08665*](https://www.ncbi.nlm.nih.gov/protein/EHY60717.1) *[Exophiala dermatitidis NIH/UT8656]* |
| *Ex11 & Ex13*  *27* | *APL5 - AP-3 complex subunit delta* [*HMPREF1120_04507*](https://www.ncbi.nlm.nih.gov/protein/EHY56425.1) *[Exophiala dermatitidis NIH/UT8656]*  *G2/mitotic-specific cyclin 3/4* [*HMPREF1120_00797*](https://www.ncbi.nlm.nih.gov/protein/EHY52586.1) *[Exophiala dermatitidis NIH/UT8656]*  *cytochrome P450 oxidoreductase* [*HMPREF1120_01361*](https://www.ncbi.nlm.nih.gov/protein/EHY53163.1) *[Exophiala dermatitidis NIH/UT8656]*  *DEAD box RNA helicase HelA* [*HMPREF1120_02010*](https://www.ncbi.nlm.nih.gov/protein/EHY53828.1) *[Exophiala dermatitidis NIH/UT8656]*  *hypothetical protein* [*HMPREF1120_02190*](https://www.ncbi.nlm.nih.gov/protein/EHY54013.1) *[Exophiala dermatitidis NIH/UT8656]*  *MFS transporter, DHA1 family, multidrug resistance protein* [*HMPREF1120_09017*](https://www.ncbi.nlm.nih.gov/protein/EHY61079.1) *[Exophiala dermatitidis NIH/UT8656]*  *hypothetical protein* [*HMPREF1120_02556*](https://www.ncbi.nlm.nih.gov/protein/EHY54387.1) *[Exophiala dermatitidis NIH/UT8656]*  *hypothetical protein* [*HMPREF1120_02949*](https://www.ncbi.nlm.nih.gov/protein/EHY54785.1) *[Exophiala dermatitidis NIH/UT8656]*  *hypothetical protein* [*HMPREF1120_03367*](https://www.ncbi.nlm.nih.gov/protein/EHY55222.1) *[Exophiala dermatitidis NIH/UT8656]*  *VPS41 - Vacuolar protein sorting-associated protein 41* [*HMPREF1120_06737*](https://www.ncbi.nlm.nih.gov/protein/EHY58734.1) *[Exophiala dermatitidis NIH/UT8656]*  *MFS transporter, SP family, sugar:H+ symporter* [*HMPREF1120_09186*](https://www.ncbi.nlm.nih.gov/protein/EHY61252.1)  *[Exophiala dermatitidis NIH/UT8656]*  *hypothetical protein* [*HMPREF1120_04659*](https://www.ncbi.nlm.nih.gov/protein/EHY56583.1) *[Exophiala dermatitidis NIH/UT8656]*  *DNA repair protein RAD50* [*HMPREF1120_04505*](https://www.ncbi.nlm.nih.gov/protein/EHY56423.1) *[Exophiala dermatitidis NIH/UT8656]*  *biphenyl-2,3-diol 1,2-dioxygenase, variant* [*HMPREF1120_05880*](https://www.ncbi.nlm.nih.gov/protein/EHY57856.1)  *[Exophiala dermatitidis NIH/UT8656]*  *chloride channel 3* [*HMPREF1120_05920*](https://www.ncbi.nlm.nih.gov/protein/EHY57899.1) *[Exophiala dermatitidis NIH/UT8656]*  *hypothetical protein* [*HMPREF1120_06002*](https://www.ncbi.nlm.nih.gov/protein/EHY57982.1) *[Exophiala dermatitidis NIH/UT8656]*  *hypothetical protein* [*HMPREF1120_06122*](https://www.ncbi.nlm.nih.gov/protein/EHY58104.1) *[Exophiala dermatitidis NIH/UT8656]*  *hypothetical protein* [*HMPREF1120_06336*](https://www.ncbi.nlm.nih.gov/protein/EHY58324.1) *[Exophiala dermatitidis NIH/UT8656]*  *ISU1 - iron-binding protein* [*HMPREF1120_06751*](https://www.ncbi.nlm.nih.gov/protein/EHY58748.1) *[Exophiala dermatitidis NIH/UT8656]*  *LBA1 - Regulator of nonsense transcripts 1-like protein [Exophiala dermatitidis NIH/UT8656]*  *hypothetical protein* [*HMPREF1120_07292*](https://www.ncbi.nlm.nih.gov/protein/EHY59300.1) *[Exophiala dermatitidis NIH/UT8656]*  *adenosinetriphosphatase* [*HMPREF1120_09246*](https://www.ncbi.nlm.nih.gov/protein/EHY61312.1) *[Exophiala dermatitidis NIH/UT8656]*  *hypothetical protein A1O3_09068 [Capronia epimyces CBS 606.96]*  *TPC1 - mitochondrial thiamine pyrophosphate transporter* [*HMPREF1120_07619*](https://www.ncbi.nlm.nih.gov/protein/EHY59634.1) *[Exophiala dermatitidis NIH/UT8656]*  *MFS transporter, SIT family, siderophore-iron:H+ symporter* [*HMPREF1120_07838*](https://www.ncbi.nlm.nih.gov/protein/EHY59858.1) *[Exophiala dermatitidis NIH/UT8656]*  *hypothetical protein* [*HMPREF1120_08665*](https://www.ncbi.nlm.nih.gov/protein/EHY60717.1) *[Exophiala dermatitidis NIH/UT8656]* |
